# The Cytomegalovirus M35 Protein Modulates Transcription of *Ifnb1* and Other IRF3-Driven Genes by Direct Promoter Binding

**DOI:** 10.1101/2023.03.21.533612

**Authors:** Hella Schwanke, Vladimir Gonçalves Magalhães, Stefan Schmelz, Emanuel Wyler, Thomas Hennig, Thomas Günther, Adam Grundhoff, Lars Dölken, Markus Landthaler, Marco van Ham, Lothar Jänsch, Konrad Büssow, Joop van den Heuvel, Wulf Blankenfeldt, Caroline C. Friedel, Florian Erhard, Melanie M. Brinkmann

**Author notes:** These authors contributed equally (alphabetical order) (Vladimir Gonçalves Magalhães, Stefan Schmelz).

## Abstract

Induction of type I interferon (IFN) gene expression is among the first lines of cellular defence a virus encounters during primary infection. We previously identified the tegument protein M35 of murine cytomegalovirus (MCMV) as an essential antagonist of this antiviral system. M35 localizes to the nucleus and interferes with type I IFN induction downstream of pattern-recognition receptor (PRR) activation. Here, we report structural and mechanistic details of M35’s function. Using electrophoretic mobility shift assays (EMSA), we demonstrate that purified M35 protein specifically binds to the regulatory DNA element that governs transcription of the first type I IFN gene induced in non-immune cells, *Ifnb1*. Determination of M35’s crystal structure combined with reverse genetics revealed that homodimerisation is a key feature for M35’s immunomodulatory activity. DNA-binding sites of M35 overlapped with the recognition elements of interferon regulatory factor 3 (IRF3), a key transcription factor activated by PRR signalling. Chromatin immunoprecipitation (ChIP) showed reduced binding of IRF3 to the host *Ifnb1* promoter in the presence of M35. We furthermore defined the IRF3-dependent and the type I IFN signalling-responsive genes in murine fibroblasts by RNA sequencing of metabolically labelled transcripts (SLAM-seq), and assessed M35’s global effect on gene expression. Stable expression of M35 broadly influenced the transcriptome in untreated cells and specifically down-regulated basal expression of IRF3-dependent genes, and during MCMV infection, M35 impaired expression of IRF3-responsive genes aside of *Ifnb1*. Our results suggest that M35-DNA binding directly antagonises gene induction by IRF3 and impairs the antiviral response more broadly than formerly recognised.

**Importance:** Replication of the ubiquitous human cytomegalovirus (CMV) in healthy individuals mostly goes unnoticed, but can impair foetal development or cause life-threatening symptoms in immunosuppressed or -deficient patients. Like other herpesviruses, CMV extensively manipulates its hosts and establishes lifelong latent infections. Murine CMV (MCMV) presents an important model system as it allows the study of CMV infection in the host organism. We previously showed that during entry, MCMV virions release the evolutionary conserved protein M35 protein to immediately dampen the antiviral type I interferon (IFN) response induced by pathogen detection. Here we show that M35 dimers bind to regulatory DNA elements and interfere with recruitment of interferon regulatory factor 3 (IRF3), a key factor for antiviral gene expression. Thereby, M35 interferes with expression of type I IFNs and other IRF3-dependent genes. Unrelated proteins from other herpesviruses employ the same mechanism, reflecting the importance for herpesviruses to avoid IRF3-mediated gene induction.

## Introduction

Upon host cell infection, viruses promptly encounter the first line of immune defence intrinsic to all nucleated cells, the type I interferon (IFN) response (1). As integral part of the innate immune system, type I IFN production is activated within a few hours of infection and links detection of a pathogen to induction of an antiviral state in infected and neighbouring cells, and ultimately in the entire organism. Stimulation of a host cell with type I IFNs invokes a broad transcriptional response that induces a cell-intrinsic defence programme including specific antiviral mechanisms, induction of pro-apoptotic and anti-proliferative pathways, and activation of the adaptive immune system. The type I IFN response thereby interferes with viral replication early on and is essential for the host organism to control infection (reviewed in (2, 3)).

Expression of type I IFNs is induced upon detection of pathogen-associated molecular patterns (PAMPs) such as aberrantly structured or localised nucleic acids by an array of pattern recognition receptors (PRRs) (4). The activation signal is subsequently relayed through adaptor proteins and kinases to the transcription factors activator protein 1 (AP-1), nuclear factor κB (NF-κB), interferon regulatory factor 3 (IRF3) and 7 (IRF7). Activation enables the transcription factors to enter the nucleus and induce expression of specific sets of genes: AP-1 dimers regulate various genes involved in cell proliferation, differentiation, and apoptosis (5), NF-κB activates proinflammatory gene expression (6), IRF3 and IRF7 together regulate expression of IFNα subtypes (7–9), and NF-κB and IRF3 or IRF7 together with an AP-1 heterodimer of ATF2 and c-Jun are required to activate transcription of the gene encoding IFNβ (*Ifnb1*) (10–13). Cells typically secrete IFNβ as the very first response to infection, and immune cells also produce specific subtypes of IFNα. These type I IFNs are in turn recognised by two type I interferon α/β receptor (IFNAR) subunits (IFNAR1 and IFNAR2) at the surface of infected and neighbouring cells. The activated IFNAR stimulates a second signalling cascade that induces assembly of signal transducer and activator of transcription 1 (STAT1) and 2 (STAT2) and IRF9 to transcription factor complexes, mainly interferon-stimulated gene factor 3 (ISGF3), and finally culminates in induction of hundreds of interferon-stimulated genes (ISGs) (14, 15). An intricate network of feed-back loops and signalling cross-talk sustains and diversifies type I IFN activity through multiple rounds of signalling, inducing the appropriate immune responses to eliminate the intruding virus (15–18). In addition, transcription of a small set of ISGs including *Isg15*, *Ifit1, Ifit2, Ifit3*, *Mx1, Mx2*, and *Rsad2*, is directly activated by IRF3, giving rise to their designation as IRF3-dependent ISGs (19, 20). During viral infection, this IRF3-mediated shortcut in the type I IFN-mediated antiviral response enables induction of gene expression before or in the absence of IFNAR activation (9, 21–23). Thus, IRF3-dependent gene expression provides the host cell with the ability to immediately deploy some of the best-studied ISGs to counter the commencing viral infection (24–27).

The ubiquitously expressed IRF3 is critical to initiate the very first round of type I IFN signalling (8, 28, 29). In contrast, IRF7 is an ISG itself and crucial for inducing high levels of ISGs and appropriate diversification of the immune response in later rounds of type I IFN signalling, including the upregulation of *Ifna* genes (7, 8, 29–31). The expression of *Ifnb1* is first induced by IRF3 and then maintained by IRF7, as these two IRFs can equally trans-activate the enhancer element that regulates induction of *Ifnb1* (8, 9). Upon PRR signalling, four transcription factor dimers together with co-factors bind to this IFNβ enhancer and co-operatively induce *Ifnb1* expression (32, 33): One AP-1 heterodimer of ATF-2 and c-Jun, two dimers of IRF3 and/or IRF7, and one NF-κB heterodimer of p50 and p65 (see scheme in Figure 6A). The two IRF3 and/or IRF7 dimers bind to four overlapping IRF-recognition elements (IREs) in the centre of the IFNβ enhancer, with each of the four DNA-binding domain contacting one 5’-GAAA-3’ consensus core element (34–36). The precise sequence arrangement of the IFNβ enhancer together with the structural orientation of the DNA-binding domains bound to this sequence indicate that the two IRF3/7 dimers bind from opposite sites to the DNA helix to overlapping parts of the sequence (32, 37).

To successfully infect and propagate in their respective host, viruses have evolved a multitude of mechanisms to inhibit, circumvent or modulate the type I IFN response. From the induction of the PRR signalling cascade to the activity of individual ISGs, all levels have been reported to be targeted by viral proteins (reviewed in (33, 38, 39)). The DNA virus family of *Herpesviridae* is especially well adapted to the host, employing many gene products that manipulate the host cells at various levels and enable the establishment of life-long infections. Human cytomegalovirus (HCMV) of the *Betaherpesvirinae* subfamily infects most humans early in life and reaches a seroprevalence of about 83% in the global adult population (40). Primary infection usually goes unnoticed in healthy patients, though it can result in a mild mononucleosis-like syndrome and has been associated with the development of chronic inflammatory diseases (41, 42). Different organs and cell types including fibroblasts, monocytes, endothelial, and epithelial cells can be affected during primary infection with CMV, with fibroblasts representing the standard cell culture model (43, 44). The type I IFN response is critical to control CMV infection and to protect the host from progression of pathogenesis (45–48). However, type I IFNs do not suffice to eliminate CMV from the organism, because the virus extensively manipulates the host (49, 50), and finally enters a latent state in specific cells of the myeloid lineage (51). Under certain conditions, such as a weakened immune system, CMV can reactivate (52). In immunocompromised patients, lytic replication of CMV after primary infection or reactivation can lead to life-threatening symptoms (53). Moreover, an active infection during pregnancy can be transmitted to the foetus and severely impair development of the unborn child, making congenital CMV infection the leading viral cause of birth defects worldwide (54).

To complement cell culture studies of strictly species-specific HCMV, murine CMV (MCMV) presents a well-established model system that enables characterisation of immune responses in the host organism (reviewed in (55, 56)). The first identified antagonist of the type I IFN response of MCMV was M27, which impairs IFNAR signalling by targeting STAT2 (57, 58). However, M27 alone does not suffice to efficiently shut off the type I IFN response in macrophages, indicating that MCMV harbours additional modulators (59). We identified the tegument protein M35 as the first MCMV antagonist of PRR-mediated *Ifnb1* transcription (60), and later on the MCMV m152 protein as a modulator of the adapter protein stimulator of interferon genes (STING) of the DNA-sensing PRR cyclic GMP-AMP synthase (cGAS) (61). In addition, we studied M35’s homologue in HCMV, UL35, and identified its immunomodulatory activity. Both UL35 and M35 are packaged into the virus particles as part of the tegument and therefore enter the host cell directly during infection, and both inhibit type I IFN signalling downstream of cGAS as well as of the RNA sensor retinoid acid inducible gene I (RIG-I), but upstream of IFNAR signalling (60, 62, 63). While UL35 impairs signalling at the level of the Tank-binding kinase 1 (TBK1) upstream of transcription factor activation (63), presence of M35 neither impairs phosphorylation-mediated activation and nuclear translocation of IRF3 nor of the NF-κB subunit p65 (60). By creating viruses deficient for production of UL35 or M35, we determined that these proteins are required for viral control the type I IFN response and efficient replication in cell culture (60, 63). Consistently, M35-deficient MCMV replicates to lower titres than wild-type (WT) MCMV in mice and does not reach the salivary glands, the organ from where MCMV would spread to the next host (60).

This underlines the critical role of M35 for successful viral replication and suggests similar importance for the homologous proteins in other herpesviruses, like UL35 of HCMV. However, the exact mechanism of action of M35 remained to be determined. After ectopic expression, M35 was detected in the nuclear fraction of the cell, and MCMV-delivered M35 entered the nucleus prior to activated p65 during infection (60). Considering that p65 has been reported to be the first and a rate-limiting transcription factor recruited to the IFNβ enhancer after induction of PRR signalling (64, 65), these observations emphasize how fast M35 reaches the nucleus. Moreover, the M35-mediated inhibition of *Ifnb1* expression was observable both in the context of infection and upon ectopic expression of M35, implying that no further viral factors were required for M35’s immunomodulatory activity (60).

Here, we report on the structural and mechanistic details of M35’s immunomodulatory activity. Using purified M35 protein, we demonstrate a direct interaction of M35 with a DNA probe containing the sequence of the IFNβ enhancer. Determination of the crystallographic structure of the major N-terminal portion of M35 (residues 7 to 334 and 376 to 441 of 519 amino acids (aa)) at 1.96 Å revealed dimerisation of M35, similar to the homologous U14 protein of human herpesvirus 6B (HHV6B) (66). We further verify the homotypic interaction of M35 in cells and show that dimers exist independently of PRR signalling activity. Characterisation of the M35 structure by reverse genetics suggests that homodimerisation is an essential feature for M35’s mechanism of action. Moreover, we report that M35-DNA interaction requires multiple consecutive core motifs of the IREs in the IFNβ enhancer sequence. Consistently, presence of M35 impairs recruitment of IRF3 to the *Ifnb1* promoter in the host cell. Furthermore, we applied metabolic labelling of RNAs and sequencing (SLAM-seq) upon PRR or type I IFN stimulation and defined the genes regulated in fibroblasts dependent on IRF3 or canonical IFNAR signalling. Comparison with SLAM-seq in M35-expressing cells suggested that the presence of M35 alters the expression of a substantial number of IRF3-dependent genes besides *Ifnb1*. Finally, we validate that M35 directly inhibits expression of several IRF3-regulated genes early during MCMV infection of macrophages. Our results show that by deploying M35 as a DNA-binding protein, MCMV specifically antagonises IRF3-mediated induction of host genes and impairs the immediate antiviral response more broadly than previously recognised.

## Results

### Purified M35 specifically binds to the sequence of the IFNβ enhancer in vitro

Based on our previous findings showing that M35 localises in the nucleus and antagonises PRR signalling downstream of transcription factor activation, we hypothesised that M35 might affect *Ifnb1* induction by direct binding to the IFNβ enhancer. To test a DNA interaction in vitro, and potentially learn more about M35’s structural features, we purified the M35 protein. Since previous analyses of HHV6B U14, a homologue of M35, indicated that the C-terminal part of the proteins was disordered (66), we generated expression constructs for purification of full-length M35 (amino acids (aa) 2-519; M35_FL) and a short version of M35 (aa 2-452; M35_S) corresponding to the structured U14 N-terminal domain (aa 2-458). The M35 coding regions were N-terminally fused to a Twin-Strep tag via a TEV protease cleavage sequence (NStr-; Figure 1A) for removal of the tag after primary protein purification. Comparable amounts of NStr-M35_S and NStr-M35_FL were obtained from transiently transfected High-Five insect cells (Figure 1B), however, the NStr-M35_FL eluates contained a second, slightly lower band (Figure 1B, lanes 7-8). As expected, this indicated that the full-length protein could not be purified to homogeneity due to a cleavage site or breakage point. In addition, NStr-M35_FL and M35_FL displayed a strong tendency to precipitate, especially at temperatures below 4°C. In contrast, M35_S could readily be purified after removal of the N-terminal tag. To confirm that the absence of the C-terminus did not hinder the immunomodulatory activity of the M35_S protein, we assessed the effect of M35_S on the induction of the *Ifnb1* promoter (Figure 1C, D). We co-transfected an expression plasmid encoding the adaptor protein of the RNA sensor RIG-I, mitochondrial antiviral-signalling protein (MAVS), to stimulate expression of a reporter plasmid expressing a luciferase gene under control of the *Ifnb1* promoter (Figure 1D, EV). As demonstrated before (60), co-expression of full-length M35 with a C-terminal V5/His epitope tag (M35-V5/His) strongly inhibited the induction of the *Ifnb1* reporter, and so did co-expression of M35_S (Figure 1D). This indicates that the C-terminal part was not required for the immunomodulatory activity of M35 after ectopic expression, and we further focused on M35_S.

**Figure 1:**
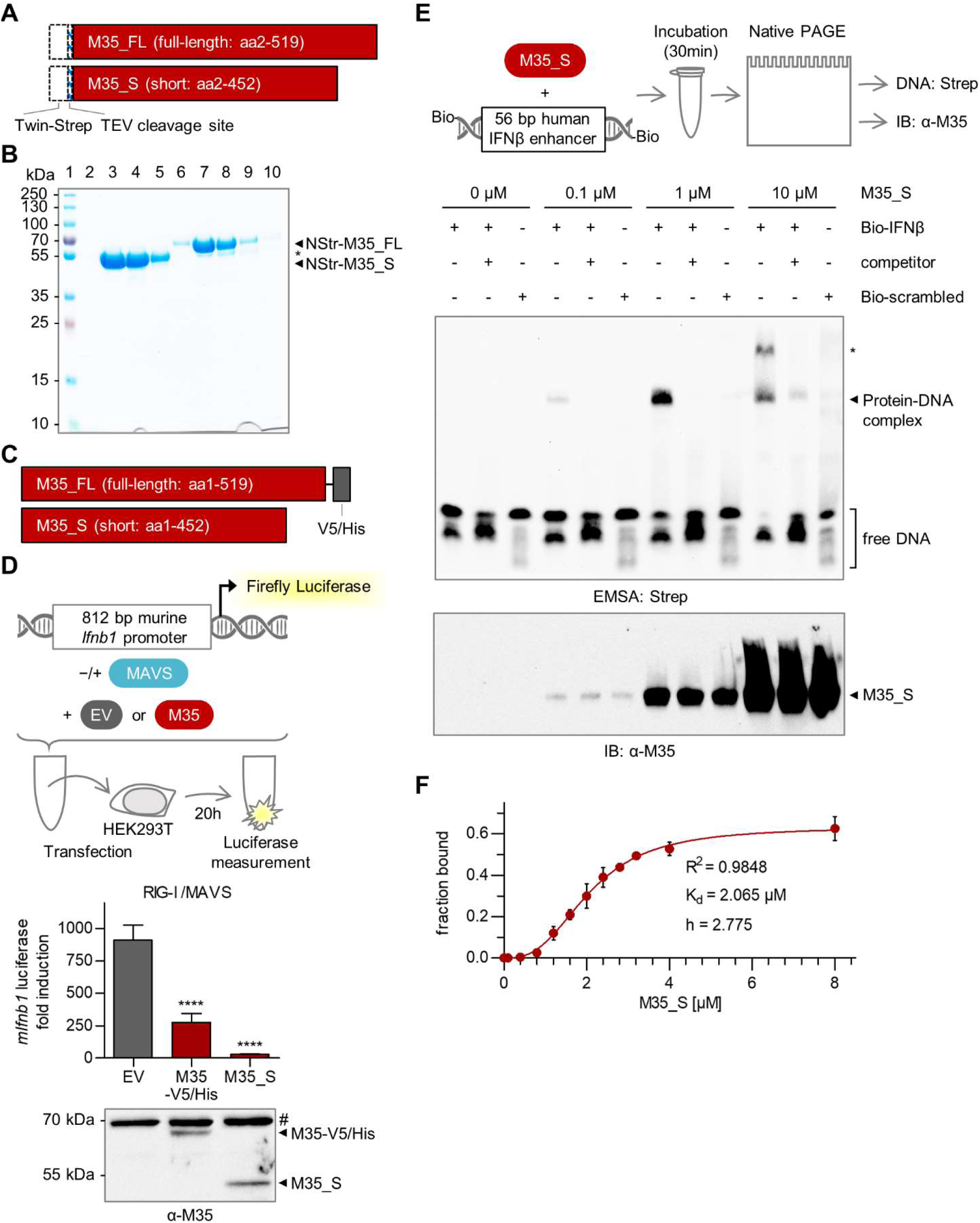
M35 specifically binds to the IFNβ enhancer sequence in vitro. (A) Expression constructs for purification of full-length (M35_FL) and short (M35_S) M35. Dashed boxes: N-terminal Twin-Strep tag and TEV protease cleavage sequence (NStr-). (B) Purification of NStr-M35_S and NStr-M35_FL proteins. NStr-M35_S and NStr-M35_FL proteins were purified from transiently transfected High-Five insect cells by StrepTactin affinity chromatography. Elution fractions were analysed by SDS-PAGE and Coomassie staining. Lane 1: Marker, lane 2-5: eluate fractions 2-5 of NStr-M35_S, lane 6-10: eluate fractions 2-6 of NStr-M35_FL. * degradation product of NStr-M35_FL. (C) Constructs for transient expression of M35 with C-terminal V5/His tag (M35-V5/His) and of untagged M35_S. (D) Analysis of M35-mediated inhibition of *Ifnb1* transcription in a luciferase reporter assay. HEK293T cells were co-transfected with the 812 bp murine *Ifnb1* luciferase reporter (*mIfnb1*-FLuc), a *Renilla* luciferase control (TK-RLuc), expression plasmids for Flag-MAVS (stimulated conditions) or a respective empty vector (EV; unstimulated conditions), and the indicated expression plasmid for M35-V5/His or M35_S or corresponding EV. Dual-luciferase measurement was performed after 20 h. Luciferase fold induction was calculated based on firefly luciferase values normalized to *Renilla* luciferase values from stimulated samples divided by corresponding values from unstimulated samples. Data are represented as mean ±SD combined from three independent experiments. Significance compared to EV was calculated by Student’s *t*-test (unpaired, two-tailed), **** p < 0.0001. Lysates were analysed by immunoblotting (IB) with an M35-specific antibody. # unspecific signal. (E) Analysis of M35-DNA binding by electrophoretic mobility shift assay (EMSA). Increasing amounts of M35_S protein were mixed with a 5’-biotinylated double-stranded 56 bp oligonucleotide probe containing the sequence of the human IFNβ enhancer (Bio-IFNβ) or a random sequence (Bio-scrambled) for control, respectively. Samples were subjected to native PAGE followed by blotting and detection of the biotinylated probes with a Streptavidin-peroxidase conjugate (Strep). Unlabelled competitor (IFNβ sequence) was added in 100x excess. A second EMSA gel was immunoblotted and analysed with an M35-specific antibody. * second protein-DNA complex. One representative of three independent experiments is shown. (F) Determination of the binding affinity of M35_S by EMSA. EMSA was performed as described in (E) with a titration series of 0.1 to 8 µM M35_S protein incubated with the murine IFNβ enhancer probe. The band intensities of bound and free probe per lane were quantified using Fiji to calculate the bound probe fraction. Values are plotted as mean ±SD of three independent experiments. The curve was fitted in GraphPad Prism using the Binding-Saturation module for specific binding with Hill slope (h) to determine the dissociation constant (K_d_).

Next, we assessed the ability of M35_S to bind to DNA in an electrophoretic mobility shift assay (EMSA), using dsDNA probes with 5’-biotin labels for detection (Figure 1E). We have previously shown that M35 inhibits induction of the human as well as of the murine *Ifnb1* promoter (60), which suggests recognition of both sequences in the case of direct M35-DNA interaction. Since the precise contact sites of the different transcription factors with the DNA nucleotides are known in the human IFNβ enhancer (37, 67), this sequence served as main probe to study specific binding (Bio-IFNβ). Incubation of increasing amounts of M35_S (0.1, 1, 10 µM) with the Bio-IFNβ probe led to a dose-dependent mobility shift, reflecting formation of a protein-DNA complex (Figure 1E). At 10 µM of M35_S protein, a second band with a lower electrophoretic mobility appeared.

Addition of a 100-fold excess of an unlabelled competitor greatly reduced the signal of the M35-DNA complex, indicating sequence-specificity. Incubation of M35_S with a biotinylated control probe harbouring a random sequence with the same GC content (Bio-scrambled) did not detectably shift the biotin signal, further confirming specificity of M35 binding to the IFNβ enhancer sequence. To determine the binding affinity of M35_S for DNA, we performed a more detailed titration series of M35_S in the EMSA using the murine IFNβ enhancer sequence as biotinylated probe to provide the natural target sequence M35 encounters in infection (Figure S1A). Quantification of the probe signals at increasing concentrations of M35_S (Table S1) and fitting of the data to a saturation model for specific binding returned a dissociation constant (K_d_) of 2.056 µM, with a Hill coefficient (h) of 2.775 suggesting cooperativity (Figure 1F).

From these data, we conclude that the first 452 amino acids part of M35 are sufficient to inhibit induction of the *Ifnb1* reporter and specifically recognise the essential enhancer sequence of the *Ifnb1* promoter in vitro. Binding of the IFNβ enhancer sequence by proteins of other herpesviruses has been suggested to inhibit *Ifnb1* induction by interfering with association of the host transcription factors (68–70). One of these proteins, K-bZIP of the Kaposi’s sarcoma-associated herpesvirus (KSHV), was initially identified as stimulator of basal *Ifnb1* promoter activity in the absence of PRR signalling, but inhibited *Ifnb1* promoter activity after induction of PRR signalling (68). Therefore, we tested whether M35 potentially activates the *Ifnb1* reporter in the absence of PRR stimulation. Similar to stimulated conditions though, M35 slightly inhibited (25%) *Ifnb1* promoter activity also in unstimulated conditions (Figure S1B).

### Structure determination reveals formation of M35 homodimers

The purified M35_S protein could be crystallised and the three-dimensional structure was determined at 1.94 Å resolution (Figure 2A, Table S2), using the homologous HHV6B U14 N-terminal domain ((66), PDB 5B1Q) as search model for molecular replacement. Similar to U14, two M35_S chains form an antiparallel homodimer with an extended interface along the long protein axis. Comparing the individual chains in the dimer to each other yielded a root-mean-square deviation (RMSD) of main-chain atoms of 0.438 Å. Most of the residues of M35_S could be located in the electron density, with the exception of the most N-terminal residues 1-6, the most C-terminal residues 442-452, and a fragment of 34 aa in M35 chain A from position 344 to 376, and 31 aa in chain B from position 346 to 375, respectively (Figure 2A). The individual M35 moieties are comprised of 14 α-helixes creating an elongated main body with two protuberant β strands forming a hairpin (Figure 2A, central panel). The β-hairpin of one monomer reaches out to the β-hairpin of the second M35 protomer of the M35 dimer (Figure 2A, top panel), constituting a prominent part of the dimer interface. At the opposite site from the β-hairpins, a groove bends along the interface (Figure 2A, bottom panel).

**Figure 2:**
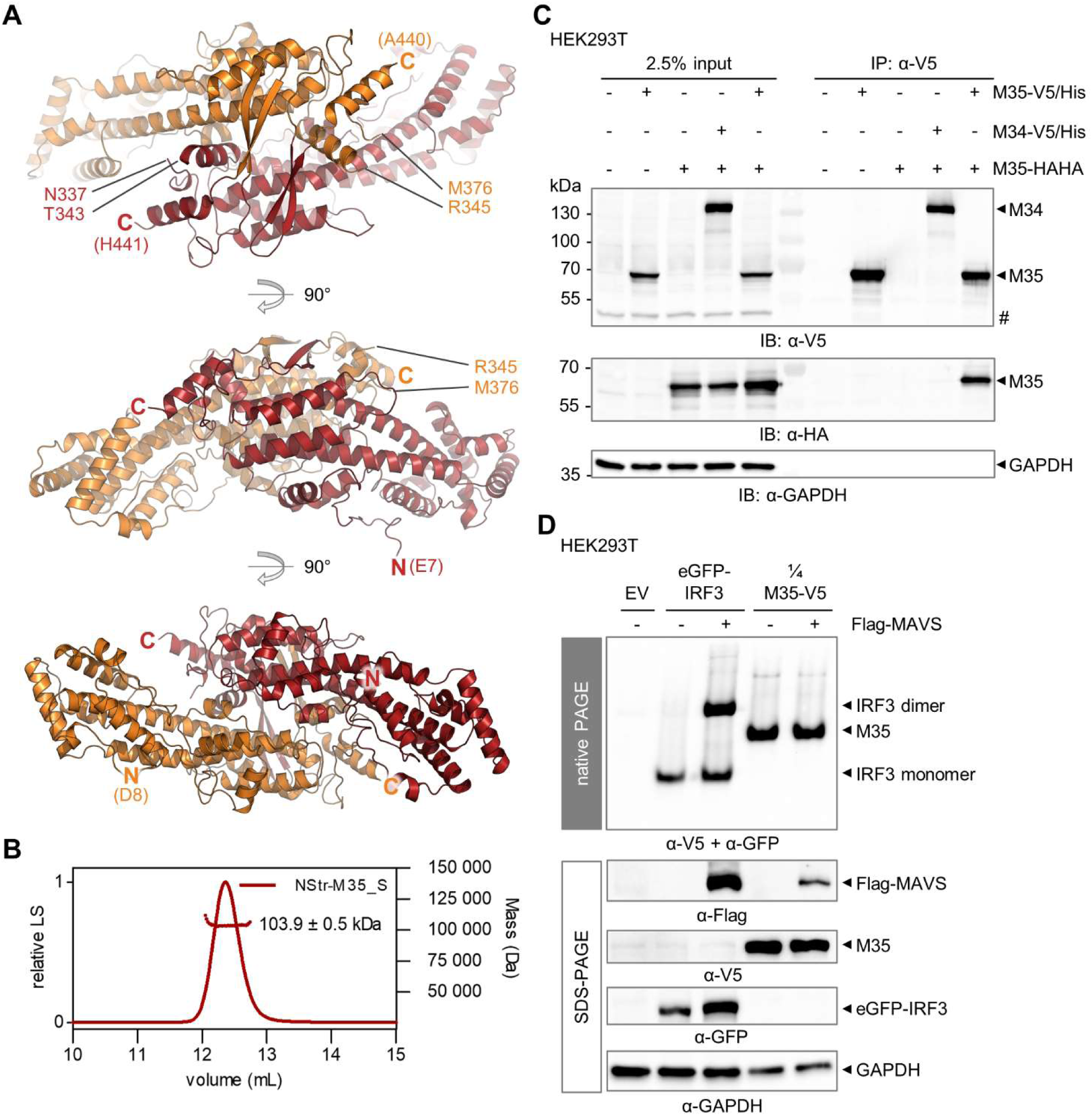
The M35 protein forms homodimers after crystallisation and in cell lysates. (A) Ribbon representations of the M35 protein crystal structure. M35_S (aa 2-452) was crystallised and its structure solved at 1.94 Å. M35 monomers are depicted in red (aa 7-343 and aa 377-441) or orange (aa 8-345 and aa 376-440), respectively. Visible N and C termini (bold) and the ends of each protein chain are labelled accordingly. The structure is depicted from three perspectives. (B) Size-exclusion chromatography followed by multi-angle light scattering (SEC-MALS) of purified NStr-M35_S protein. LS: light scattering. (C) Co-immunoprecipitation of M35 in cell lysates. HEK293T were co-transfected with indicated expression plasmids for M35-V5/His and M35-HAHA, M34-V5/His and M35-HAHA (negative control), or single constructs filled up with EV. An anti-V5-immunoprecipitation (IP) was performed 24 h later. Input and IP samples were analysed by SDS-PAGE and immunoblotting with HA- and V5-specific antibodies. Detection of GAPDH served as loading control, # unspecific band. One representative of two independent experiments is shown. (D) Native PAGE of M35 in cell lysates. HEK293T cells were co-transfected with expression plasmids for eGFP-IRF3 (control) or M35-V5/His or the corresponding EV, and for Flag-MAVS (stimulated conditions) or the respective EV (unstimulated conditions). Cells were lysed 20 h later and analysed in parallel by native (upper panel) or SDS-PAGE (lower panel) followed by immunoblotting and detection with GFP-, V5-, Flag- and GAPDH-specific antibodies as indicated. Lysates of M35-V5/His-expressing cells were diluted 1:4 in lysis buffer to adjust the signal strength in the native immunoblot. One representative of three independent experiments is shown.

Analysis of the purified NStr-M35_S protein by size-exclusion chromatography followed by multi-angle light scattering (SEC-MALS) confirmed uniform particles of 103.9 ±0.5 kDa in solution (Figure 2B). This is about twice the theoretical molecular weight of an NStr-M35_S monomer (54.5 kDa), suggesting dimerisation. Next, we studied the homodimerisation of M35 in lysates of eukaryotic cells. Co-expression of M35-V5/His and C-terminally HAHA-tagged M35 (M35-HAHA) in HEK293T cells followed by immunoprecipitation for the V5 epitope showed that M35-HAHA readily co-precipitated with M35-V5/His (Figure 2C), supporting a homotypic interaction. In contrast, M35-HAHA did not co-precipitate with a different nuclear V5/His-tagged protein of MCMV, M34 (71). Further, analysis of M35 in HEK293T lysates by native PAGE and immunoblot showed that M35 forms one defined species (Figure 2D). An eGFP-IRF3 fusion protein was included as control and as expected dimerised upon PRR signalling activation by overexpression of MAVS. Unlike eGFP-IRF3 dimers, the oligomerisation status of M35 was independent of MAVS co-expression.

Taken together, we here present the crystal structure of the domain of M35 which harbours its immunomodulatory activity. M35 forms homodimers and our results confirm that this is most likely the native state of M35 in cells and independent of PRR signalling.

### Protein folding is conserved between M35 of MCMV and U14 of HHV6B, two members of the pp85 protein superfamily of betaherpesviruses, but not their function

Based on homology to the 85 kDa phospho-protein U14 of human herpesvirus 7 (HHV7), MCMV M35 is grouped in the pp85 protein superfamily that is conserved within the *Betaherpesvirinae*, but not the *Alpha*- or *Gammaherpesvirinae* (72–74). For closer inspection, we performed multiple and pairwise sequence alignments of the members of the pp85 superfamily (Table S3, File S1). The resulting phylogenetic tree precisely mirrors the division of the betaherpesviruses into different genera, and pairwise sequence comparisons of all proteins to MCMV M35 yielded amino acid identities from up to 50% for the most closely related *Muromegalovirus* homologues to about 20% for the U14 proteins from the genus *Roseolovirus* (Figure 3A). Aside from M35, only the crystal structure of U14 of HHV6B has been reported so far from the pp85 protein superfamily (66). Superposition of the dimer structures of M35 and U14 clearly reflects their structural similarity (Figure 3B), yielding an RMSD of main chain C_α_ atoms of 2.51 Å for the superposition of dimers, and superposing the individual chains yielded even closer overlaps, with RMSDs of 1.96 Å for comparison of A chains and 2.15 Å for B chains of M35 and U14, respectively.

**Figure 3:**
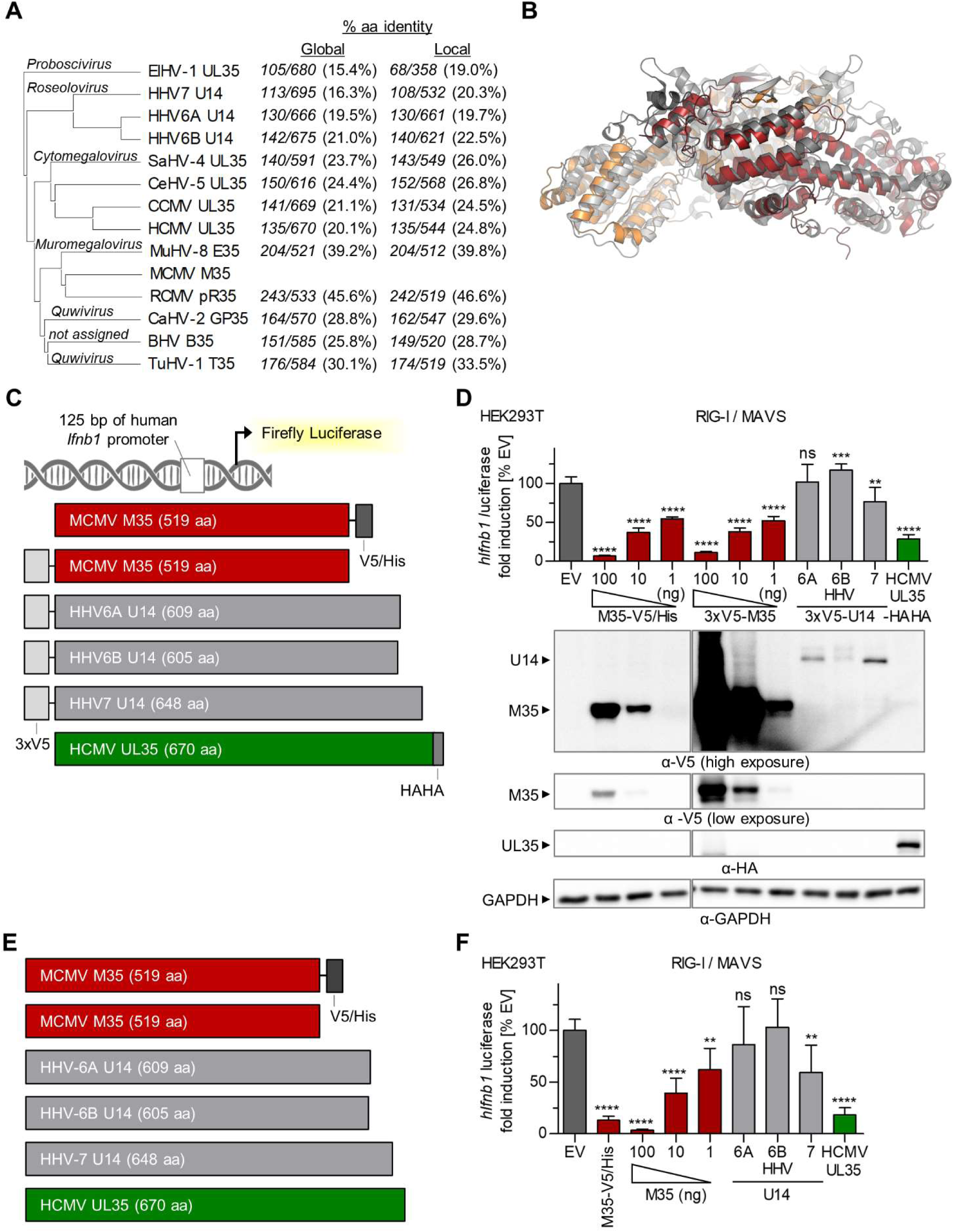
Comparison of MCMV M35 with the homologous U14 proteins of HHV6A, HHV6B and HHV7 of the *Betaherpesvirinae* pp85 protein superfamily regarding a potential inhibition of *Ifnb1* promoter induction. (A) Phylogenetic tree of the pp85 protein superfamily of the *Betaherpesvirinae*. Included are homologous proteins from all betaherpesviruses infecting humans and at least one species per genus. The table indicates the percentage of identical amino acids (% aa) as well as aligned and total sequence lengths for all proteins compared to M35 after alignment of sequences from end-to-end (global), or of the most similar regions (local). (B) Superposition of the dimers of MCMV M35 (orange-red) with the homologous HHV6B U14 (grey shades; PDB 5B1Q (66)). (C, E) Schemes of the firefly luciferase reporter construct controlled by the 125 bp human *Ifnb1* promoter (*hIfnb1*-FLuc), and of expression constructs for M35 of MCMV, U14 of HHV6A, HHV6B or HHV7, and UL35 of HCMV (C) with or (E) without tags. (D, F) Analysis of HHV6A, HHV6B, and HHV7 U14 proteins for inhibition of *Ifnb1* transcription in the luciferase reporter assay. Luciferase reporter assays were performed as described before (Figure 1D) by transfection of HEK293T cells with a Flag-MAVS-expressing plasmid for stimulation, *hIfnb1*-FLuc, and indicated expression plasmids for M35, UL35, and U14 proteins (D) with N- or C-terminal or (F) without tags. Expression constructs for (D) M35-V5/His and 3xV5-M35 or (F) untagged M35 were applied in a titration series (100, 10, 1 ng), filled up with EV to 100 ng. Data were normalised to EV samples and are represented as mean ±SD combined from three independent experiments. Significance compared to EV was calculated by Student’s *t*-test (unpaired, two-tailed) comparing M35 derivatives to EV, ns not significant, ** p < 0.01, *** p < 0.001, **** p < 0.0001. (D) Lysates were analysed by immunoblotting using V5- and HA-specific antibodies. Detection of GAPDH served as loading control.

Little is known to date about the functions of the *Roseolovirus* U14 proteins, and to our knowledge, no U14-mediated inhibition of the type I IFN response was reported. We generated expression constructs of the HHV6A, HHV6B, and HHV7 U14 ORFs, adding an N-terminal triple V5-epitope tag (3xV5-) for detection (Figure 3C), and assessed their effects in the *Ifnb1* luciferase reporter assay. Since these proteins are expected to target the human IFNβ enhancer, we co-transfected a reporter containing the human *Ifnb1* promoter sequence, again adding the MAVS expression plasmid for stimulation (Figure 3C). Similar to M35-V5/His, co-expression of analogously designed 3xV5-M35 efficiently inhibited *Ifnb1* promoter activation (Figure 3D) and so did co-expression of the C-terminal tagged HCMV homologue of M35 and U14, UL35-HAHA, consistent with our previous work (63). Out of the three U14 proteins, only HHV7 U14 downmodulated induction of the human *Ifnb1* promoter (p < 0.01). Notably, immunoblots suggested overall lower protein levels of the 3xV5-U14 proteins relative to 3xV5-M35. Still, 3xV5-U14 proteins could be detected, in contrast to M35-V5/His after transfection of only 1% of plasmid, which sufficed for significant (p < 0.0001) inhibition. To rule out that the epitope tag interfered with the putative function of U14 proteins, additional expression constructs were generated to study U14, UL35 and M35 without any modification (Figure 3E), and similar results were obtained (Figure 3F).

All in all, these data support the notion that at least some features are conserved within the pp85 protein superfamily. On the one hand, we found that neither the HHV6B nor the HHV6A U14 proteins inhibited induction of the *Ifnb1* promoter, despite HHV6B U14’s high structural similarity to M35, suggesting that the property to specifically bind DNA is not conserved in the overall fold. On the other hand, the U14 protein of HHV7 reduced induction of the human *Ifnb1* promoter, though considerably less than M35 or UL35, revealing a potential parallel between M35 and another homologue in the *Betaherpesvirinae*. From a structural perspective, it thus remains unclear why HHV6B or HHV6A U14 did not exhibit inhibitory activity in our reporter assay, and additional studies will be required to shed light on these functional differences.

### Identification of loss-of-function mutants of M35 by reverse genetics

The crystallographic structure of M35_S provided a basis to dissect the contribution of individual structural features to the immunomodulatory activity of the M35 protein. In particular, identification of residues essential for M35’s activity could potentially allow us to connect the molecular function with a structural feature, such as a putative DNA-binding site. Aiming to disrupt the function of M35, we focused mutagenesis on prominent surface features, and used the MAVS-stimulated *Ifnb1* reporter assay to screen for loss-of-function derivatives. The WT M35-V5/His protein served as basis to generate mutants and was included as control.

Firstly, we deleted the β-hairpins (aa 406-424; Δβ) or replaced them with a single proline (Δβ+P) or glycine (Δβ+G) residue to bridge the distance to the continuing protein chain (Figure 4A). All three Δβ derivatives lost the ability to inhibit induction of the *Ifnb1* luciferase reporter, indicating that the β-hairpins are an important feature of the M35 protein. Compared to WT M35-V5/His, the mutants yielded slightly reduced protein levels in control immunoblots, but were still readily detectable (Figure 4A).

**Figure 4:**
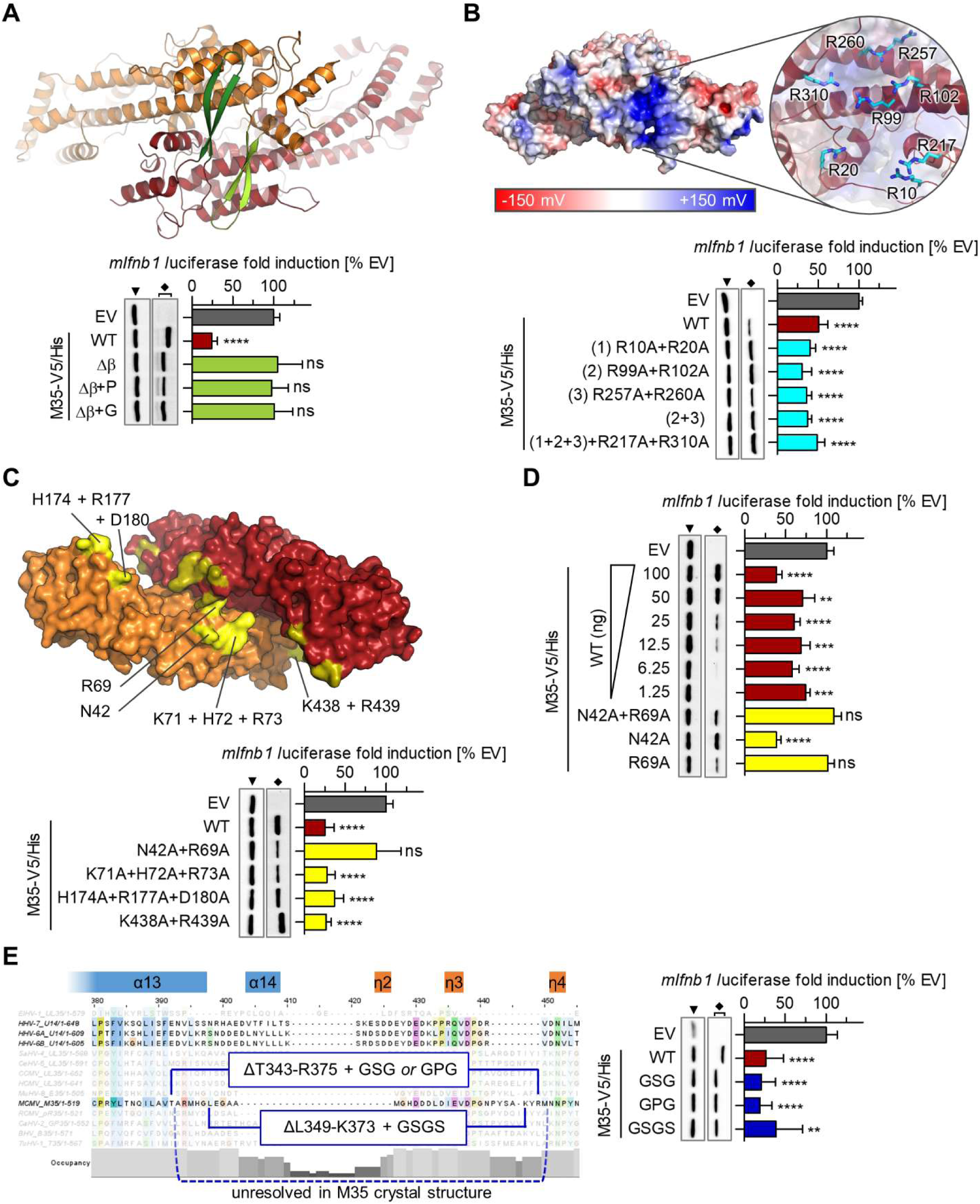
Identification of loss-of-function mutants of M35 by reverse genetics. Prominent surface features of the M35 structure and screening for of M35 mutants for a loss of inhibition of *Ifnb1* transcription in a luciferase reporter assay. (A) Top-view of the M35 dimer as ribbon representation, with the two protruding β-hairpins (aa406-424, green shades). The β- hairpins were deleted entirely (Δβ) or substituted by a single proline (Δβ+P) or glycine (Δβ+G). (B) Side-view of the M35 dimer, with depiction of the electrostatic surface potential (left; red: negative, blue: positive) and close-up on the underlying cluster of arginine residues (cyan sticks; right). Indicated arginine were mutated to alanine: 10 and 20 (1), or 99 and 102 (2), or 257 and 260 (3), combination of R10A, R20A, R99A, R102A, R257A and R260A (1+2+3), combination of (1+2+3) with R217A and R310A. (C) Bottom-view of the M35 dimer as surface representation, showing the groove formed along the dimer interface. Clusters of surface-exposed residues (yellow shades) were mutated to alanine, creating N42A+R69A, K17A+H72A+R73A, H174A+R177A+D180A, K438A+R439A (indicated on one M35 monomer). (D) The positions of the double mutant N42A+R69A were analysed individually (N42A, R69A, respectively) alongside a titration series of the wild-type (WT) M35 protein, filled up with EV to 100 ng. (E) Annotated structural elements of HHV6B U14 ((66); α and ƞ helixes) and the alignment of the pp85 protein superfamily served as basis to design mutants (navy) of the unresolved loop of M35, replacing the whole unresolved region (T343-R375) with a GSG or GPG linker, or L349-K373 with a GSGS linker. (A-E) Luciferase reporter assays were performed as described before (Figure 1D) by transfection of HEK293T cells with a Flag-MAVS-expressing plasmid for stimulation, *mIfnb1*-FLuc, and indicated expression plasmids. Data were normalised to EV samples and are represented as mean ±SD combined from three independent experiments. Significance compared to EV was calculated by Student’s *t*-test (unpaired, two-tailed) comparing M35 derivatives to EV, ns not significant, ** p < 0.01, *** p < 0.001, **** p < 0.0001. Lysates were analysed by immunoblotting with V5- and GAPDH-specific antibodies, filled arrow heads mark GAPDH, diamonds mark M35-V5/His derivatives.

Secondly, we assessed the electron surface potential of the dimer and identified a positive surface patch at the side of each M35 monomer formed by eight arginine residues (R10, R20, R99, R102, R217, R257, R260, R310; Figure 4B). Since this could provide a site for DNA interaction, we exchanged these residues in different combinations for alanine residues. However, even the exchange of all residues did not impair the inhibitory effect by these M35 derivatives, suggesting that this feature is not critical for the assayed activity.

Thirdly, we inspected the groove that runs along the dimer interface. Due to its bend and asymmetric elevations of the walls at the interface creating deep and shallow stretches, we approximated the size of the groove with a width of about 20 Å from wall to wall (Figure S2A), and roughly 83 Å from one end to the other (Figure S2B). These dimensions could accommodate a B-DNA double helix at a length of approximately 21 base pairs (75), indicating this as a candidate site for DNA binding. We exchanged neighbouring positions with surface-exposed hydrophilic residues along the groove for alanine residues, generating four mutants (N42A+R69A, K71A+H72A+R73A, H174A+R177A+D180A, K438A+R439A; Figure 4C). The double mutation N42A+R69A abrogated the inhibitory effect of M35, and again, the loss-of-function derivative yielded reduced protein levels compared to WT M35 (Figure 4C). Individual exchange of the two positions showed that the mutation R69A alone was sufficient to disrupt M35’s activity (Figure 4D). A titration of WT M35-V5/His was included to demonstrate that co-transfection of a hundredth of the standard amount (100 ng) of expression construct for M35-V5/His WT protein sufficed for significant (p < 0.001) downmodulation of *Ifnb1* promoter induction despite undetectable protein levels in the immunoblot (Figure 4D). In contrast, M35-V5/His R69A was detectable but did not notably influence luciferase induction, indicating that its loss-of-function was not or not alone due to the reduced protein level.

Fourthly, we characterised the part of M35 that was not resolved in the crystal structure (M35 aa position 344 to 376 of chain A and position 346 to 375 of chain B). In the structure of HHV6B U14 (66), the first part of the corresponding segment constitutes the end of an α-helix (α13) and then forms a loop containing small helix elements (α14, 3_10_-helixes ƞ2 to ƞ5) that reaches back to the bulk structure close to where it reached out (Figure 4E, Figure S2C). Based on the superposition of M35 and U14 (Figure S2C) and the alignment of the pp85 superfamily (Figure 4E), we replaced the unresolved residues T343 to R375 of M35 with (i) GSG or (ii) GPG linkers, or substituted only the segment L349 to K373 starting after the potentially continuing α13 helix with a (iii) GSGS linker. However, despite about 30 amino acids lacking from M35, all derivatives still inhibited induction of the *Ifnb1* reporter, indicating that the loop is not critical for the immunomodulatory activity of M35 (Figure 4E).

In sum, reverse genetic characterisation of M35 led to the identification of the β-hairpins at one side of the structure and the surface-exposed R69 at the opposite site as critical parts for the inhibitory function. Interestingly, similar to the β-hairpins, R69 is also located directly at the dimer interface, and faces the residue R69 of the second M35 moiety in the homodimer (Figure S2D). Closer inspection of the electron density revealed that each R69 residue adopts two conformations with similar occupancy, potentially allowing for π-stacking with the opposite R69 residue, or for interaction with D44 of the opposite M35 chain, respectively. In this way, interaction of the M35 chains via R69 might contribute to the homodimerisation.

### Loss-of-function mutants suggest that dimerisation is a critical feature of the M35 protein

As we generated loss-of-function mutants aiming to identify a position that specifically and directly contributes to M35’s immunomodulatory function, we further characterised M35 Δβ and M35 R69A. After immunolabelling of transfected HEK293T cells, both M35-V5/His Δβ and M35-V5/His R69A displayed a nuclear localisation (Figure 5A). Similar to WT M35 (60), the R69A mutant was dispersed throughout the nucleus, while the Δβ mutation led to the formation of distinct speckles. The overall signal for R69A was weaker compared to WT M35, corresponding to the protein levels detected by immunoblot (Figure 4D). Further, analysis of the M35 derivatives by native PAGE and immunoblot revealed that the two loss-of-function mutants of M35 were markedly different from the WT protein: Only a small fraction of M35-V5/His Δβ protein displayed the same running distance as WT M35-V5/His, and the major share moved faster through the gel creating an additional band. M35-V5/His R69A gave also rise to the faster migrating protein species, with comparable amounts for this and the WT-like species (Figure 5B). Based on our description of the WT M35 protein as a dimer (Figure 2) and the distinct shift between the WT-like and the faster moving species, we propose that the latter represents M35 monomers. This observation indicates that mutations Δβ and R69A severely impaired homodimerisation of M35. Thus, M35 Δβ and M35 R69A lost their ability to inhibit the *Ifnb1* promoter due to the impact on their overall integrity. Though we did not directly determine the DNA-binding site, this finding highlights the importance of homodimerisation for M35’s activity.

**Figure 5:**
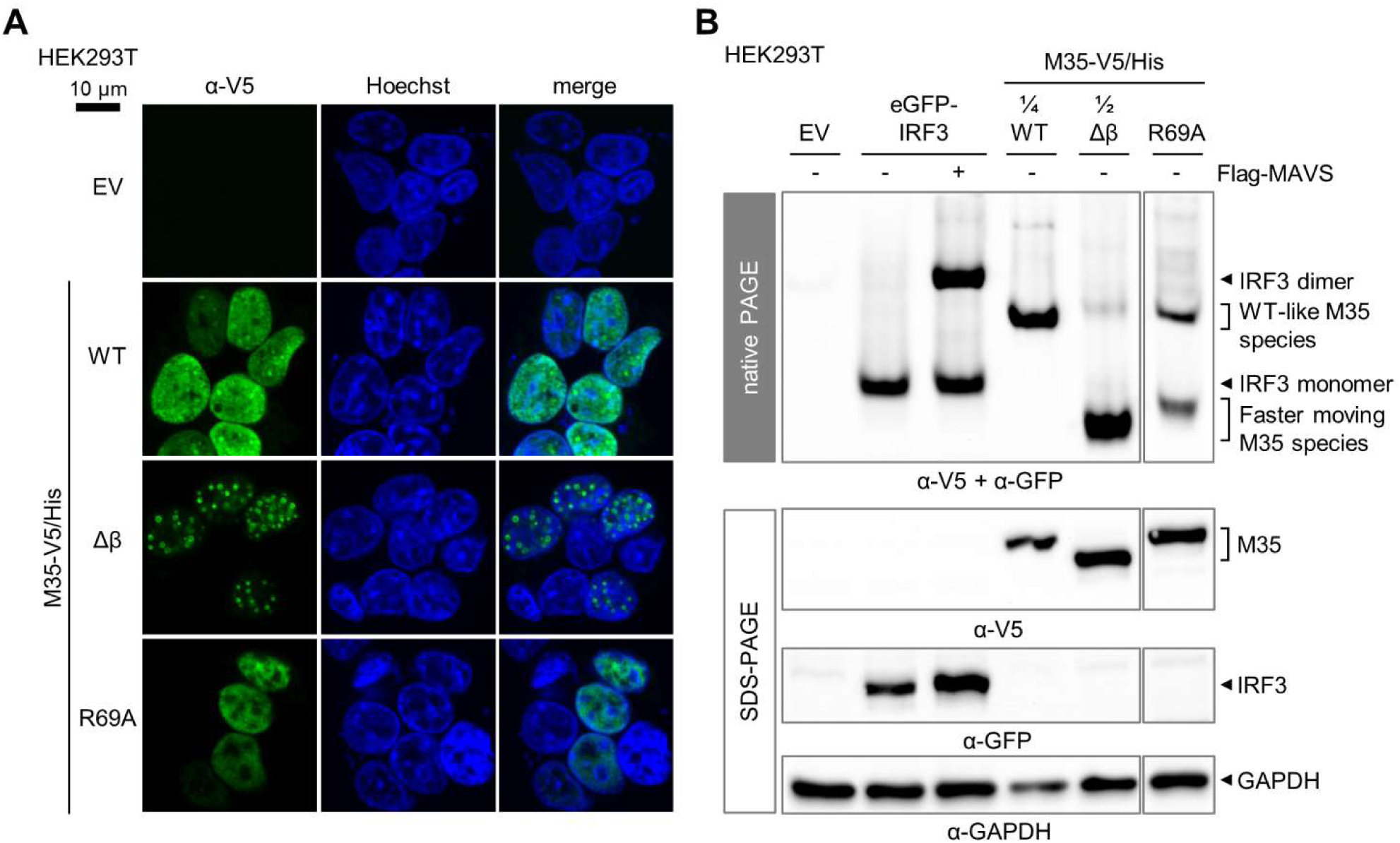
Identified M35 loss-of-function mutations impair the homodimerization of M35. (A) Immunofluorescence assay of M35 derivatives. HEK293T cells transfected with expression constructs for M35-V5/His WT, Δβ, or R69A or the corresponding EV were subjected to immunofluorescence labelling with a V5-specific antibody. Nuclei were stained with Hoechst. The scale bar represents 10 µm. Images are representative of at least two independent experiments. (B) Native PAGE of M35 derivatives. Native (upper panel) and SDS-PAGE (lower panel) followed by immunoblotting were performed as described before (Figure 2D) by co-transfecting HEK293T with expression plasmids for eGFP-IRF3, or M35-V5/His WT, Δβ, or R69A or the respective EV, and for Flag-MAVS (stimulated conditions) or the respective EV (unstimulated conditions), and analysis with GFP-, V5-, Flag- and GAPDH-specific antibodies. Lysates with M35-WT and M35-Δβ were diluted as indicated in lysis buffer to adjust the signal strength in the native immunoblot. One representative of three independent experiments is shown.

### M35-DNA recognition requires successive core motifs of IRF recognition elements

To study the protein-DNA interaction further, we next dissected the sequence requirements for M35-DNA binding by EMSA. Using the human IFNβ enhancer as a blueprint (Figure 6A), we replaced specific binding elements while keeping the probe length constant. Note that though the scheme in Figure 6A indicates alternating occupation of IREs by IRF3 and IRF7 according to the report by Panne and colleagues (37), due to the lack of steady state IRF7 expression all IREs will initially be occupied by IRF3 upon primary infection of non-immune cells.

**Figure 6:**
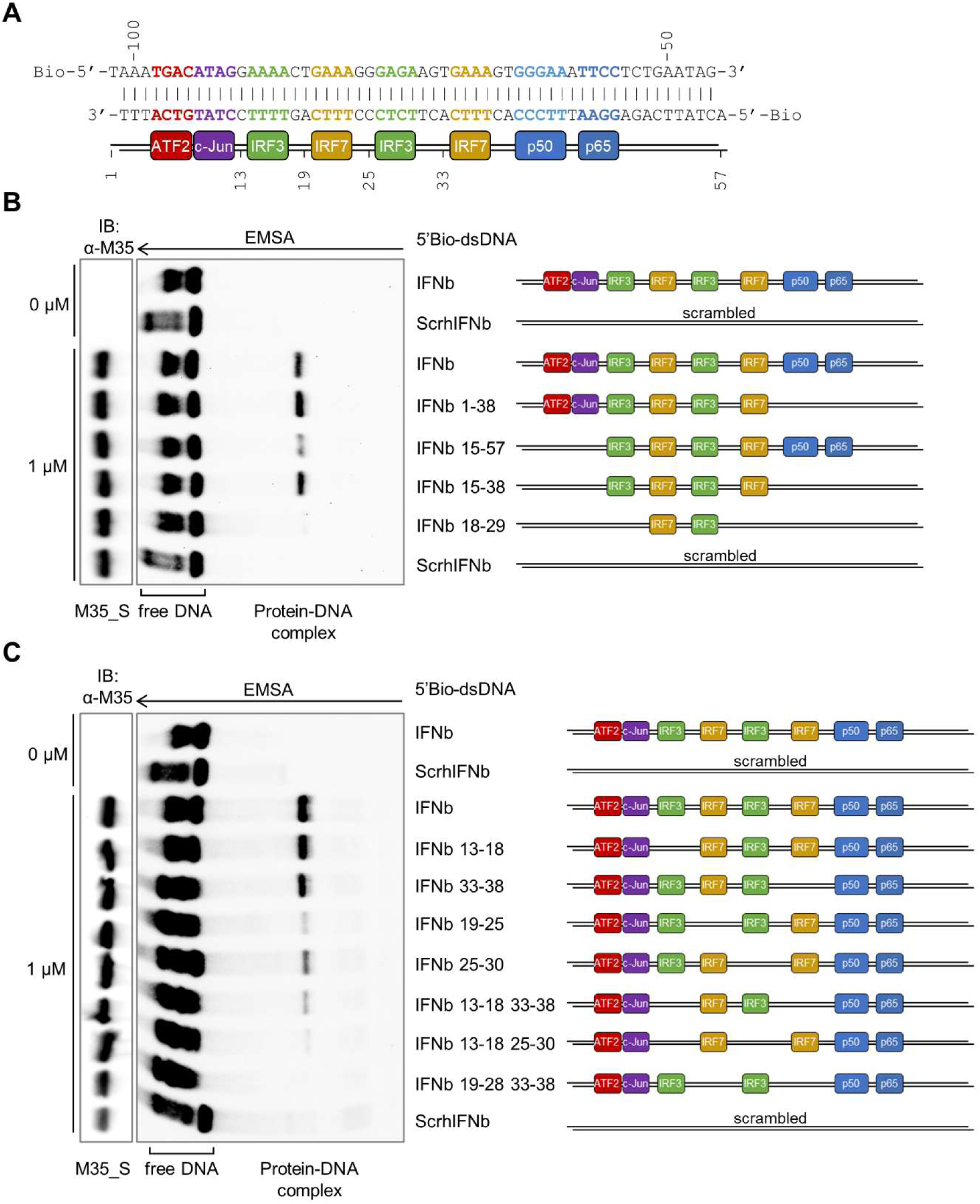
The consecutive core motifs of the IRF recognition elements in the IFNβ enhancer are required for M35-DNA binding. (A) DNA sequence of the human IFNβ enhancer upstream of the *Ifnb1* gene with positions relative to the transcription start site in bp indicated above the sequence. Both strands were 5’- biotinylated for detection of EMSA probes. Core sequence elements interacting with individual DNA-binding domains are highlighted in the same colours as the respectively bound dimeric transcription factors (based on Panne et al., 2007 (37)). The positions within the probe are indicated below. (B-C) EMSAs for M35 binding to specific sections of the IFNβ enhancer. EMSAs were performed as described before (Figure 1E) with 1 µM of purified M35_S and indicated 5’- biotinylated 56 bp dsDNA probes based on the sequence of the human IFNβ enhancer. A probe with a random sequence (scrambled: ScrhIFNb) served as negative control. (B) The sequence sections marked in the probe label were retained and remaining flanking sections replaced for random sequences. (C) The 6 bp sequence sections marked in the probe label were replaced for random sequences to mutate individual parts of the IRF recognition elements. The arrow marks the running direction of the EMSA gel. One representative of three independent experiments is shown.

First, we studied contribution of the different transcription factors binding motifs by scrambling the recognition elements individually or in combination (Figure 6B). Lack of the NF-κB (IFNb 1-38) or both ATF2/c-Jun and NF-κB binding motifs (IFNb 15-38) still allowed for formation of the M35-DNA complex, though less signal of a protein-DNA complex was observed for the probe lacking only the ATF2/c-Jun motif (IFNb 15-57). Overall, this narrowed down the M35-bound sequence to the central repeat of IREs, and congruently, the signal of DNA-protein complex was drastically reduced when only two of the four core IRE motifs were intact (IFNb 18-29). We additionally sought to narrow down if individual IREs enable M35-DNA recognition with probes in which different combinations of the core 5’-GAAA-3’ motifs were scrambled. Analysis of M35-DNA binding with the yielded array of probes revealed that the signal of the protein-DNA complex gradually decreased with fewer immediately successive core IRE motifs (Figure 6C). This suggests that instead of contacting a short sequence, the M35 binding site broadly overlaps with the binding site of IRF3/7 dimers.

These results show that the M35 binding sequence coincides with the recognition elements for IRF3/7 binding, whereas motifs for NF-κB or ATF2/c-Jun binding were not essential. This finding was in contrast to our previous report, which had suggested that M35 targeted NF-κB-mediated transcription (60). We therefore tested if M35 inhibits *Ifnb1* promoter induction when activation was directly dependent on either IRF3 or NF-κB. In agreement with the EMSA data, co-expression of M35-V5/His, but not by the IFNAR-signalling antagonist M27, significantly (p < 0.0001) reduced *Ifnb1* luciferase reporter activity induced by transient expression of constitutively active IRF3-5D (Figure S3A). In contrast, induction of the *Ifnb1* luciferase reporter by transient expression of the intrinsically active NF-κB subunit p65 was not impaired by M35-V5/His (Figure S3B). This supports the results here, which indicate that the immunomodulatory activity of M35 is independent of NF-κB or its binding motifs.

### Presence of M35 impairs binding of IRF3 to the host’s IFNβ enhancer upon stimulation of PRR signalling

Since M35-DNA binding requires IREs, we next asked whether M35 would impair recruitment of IRF3 to the IFNβ enhancer. To address this, we used immortalised mouse embryonic fibroblasts (iMEFs) that stably express the previously characterised M35-myc/His (60) to perform chromatin immunoprecipitation (ChIP) with an IRF3-specific antibody (Figure 7A). Immunoblotting of chromatin fractions validated expression of M35-myc/His and phosphorylation of IRF3 after transfection with the dsRNA mimetic poly(I:C) (Figure 7B). Enrichment of the *Ifnb1* promoter sequence in the ChIP eluates was measured by qPCR, and as expected, the fraction of *Ifnb1* promoter sequences bound by IRF3 was greatly increased in control cells upon PRR stimulation compared to mock-treatment (Figure 7C). Strikingly, stimulation-induced enrichment of IRF3 at this promoter was significantly decreased in iMEFs stably expressing M35-myc/His (p < 0.01). This suggests that the presence of M35 in host cells impairs the binding of endogenous IRF3 to its target sequence in the *Ifnb1* promoter upon PRR signalling.

**Figure 7:**
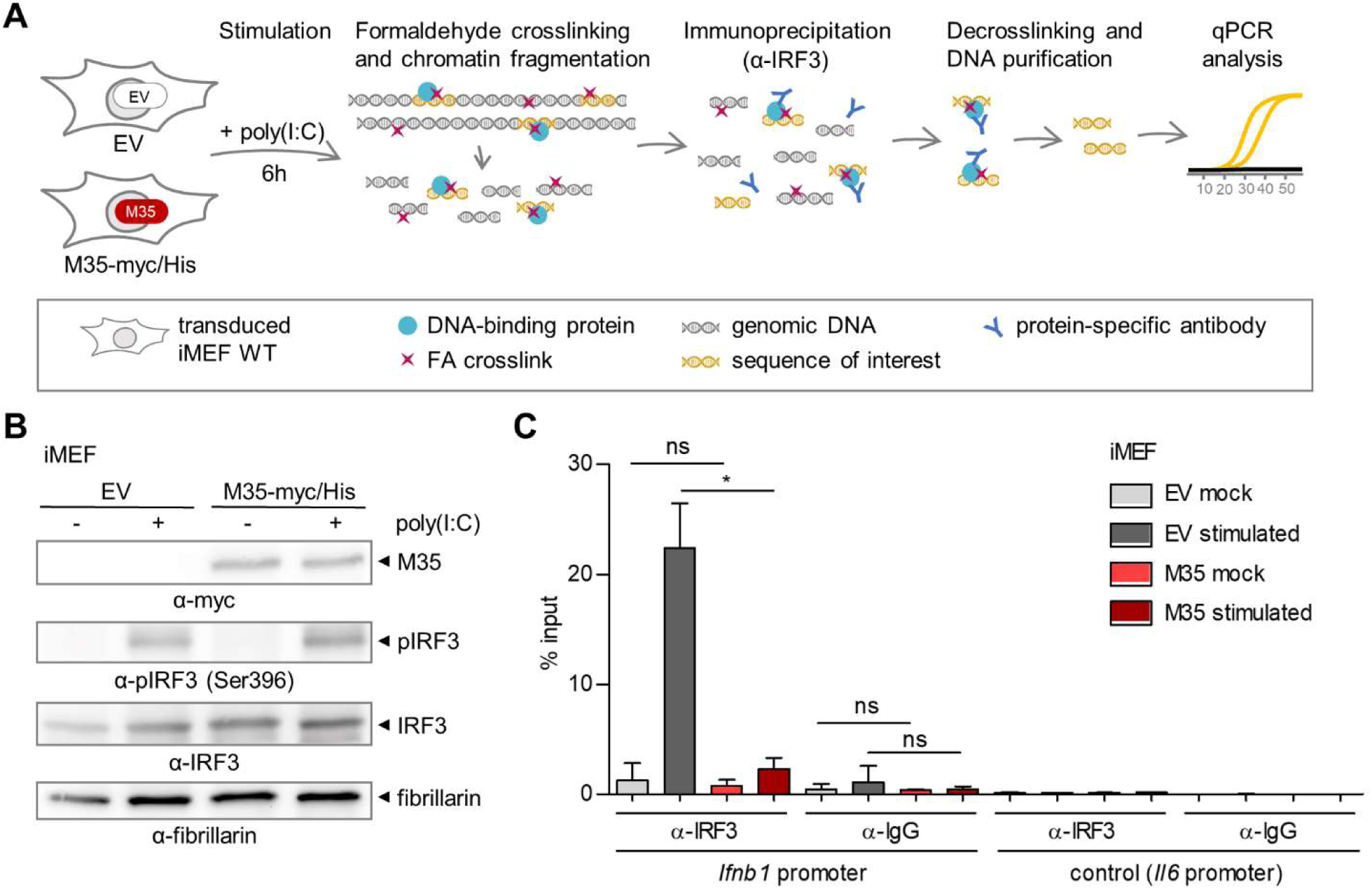
Presence of M35 impairs binding of IRF3 to the host’s IFNβ enhancer upon stimulation of PRR signalling. (A) Chromatin immunoprecipitation (ChIP) assay. iMEFs stably expressing M35-myc/His or the corresponding EV were stimulated by transfection of poly(I:C) or mock-treated. After 6 h, formaldehyde (FA) was applied to cross-link interactions and cells were harvested. Chromatin was isolated, fragmented for processing, and subjected to immunoprecipitation with an IRF3- specific antibody. The precipitated material was decrosslinked, DNA was purified and analysed by qPCR alongside 1% of input material. (B) Immunoblot of chromatin samples from iMEFs. iMEFs were processed as described in (A) and analysed by immunoblotting with myc-, pIRF3-, and IRF3- specific antibodies, and fibrillarin-specific antibodies. Fibrillarin served as a loading control for the nuclear fraction. Shown is one representative of three independent experiments. (C) ChIP for recruitment of IRF3 to the *Ifnb1* promoter in presence or absence of M35. ChIP was performed as described in (A) with an IRF3-specific and an IgG control antibody, and samples were analysed for enrichment of the IFNβ enhancer sequence by qPCR. A primer set targeting the promoter of *Il6* upstream of a predicted IRF3 binding site was used as negative control. Shown are combined data from two independent experiments.

Taken together, our data indicate that the viral protein M35 localises to the nucleus where it binds to specific host DNA sequences by recognition of motifs in IRF3/7 binding sites. As presence of M35 does not influence activation, total or stimulus-induced nuclear levels of the transcription factors NF-κB or IRF3 (60), we conclude that binding of M35 to the IFNβ enhancer competitively impairs binding of IRF3 to the same site and thus antagonises induction of *Ifnb1* transcription.

### Dissection of the contribution of IRF3- versus type I IFN signalling-mediated induction of antiviral genes in murine fibroblasts

While type I IFNs represent a major target of IRF3-mediated gene regulation, several reports have demonstrated that during viral infection, IRF3 also regulates expression of a subset of ISGs (22–24, 76). During HCMV infection of fibroblasts, some IRF3-dependent ISGs are upregulated to a similar extent by IRF3 and type I IFN-IFNAR signalling, and others are only fully induced when both pathways are activated (24). Since the M35 recognition site overlapped with IREs and M35’s presence impaired binding of IRF3 to the *Ifnb1* promoter after PRR stimulation, we wondered whether other IRF3-regulated transcripts are influenced by M35.

The direct induction of specific ISGs by IRF3 was reported by several groups studying human cells (20, 77), but is to our knowledge less well characterised in murine cells. Aiming to obtain a full picture of M35’s effect on mRNA transcription in the host cell, we applied RNA sequencing of metabolically-labelled transcripts (SLAM-seq) (78, 79). In this method, the nucleotide analogue 4-thiouridine (4sU) is incorporated into nascent RNA for a defined time, and after sequencing this enables quantification of transcripts synthesised in this time window. For direct comparison of transcripts affected by M35 with those regulated by IRF3 or type I IFN-IFNAR signalling, we firstly used SLAM-seq to characterise the IRF3-dependet versus type I IFN signalling-responsive genes in murine fibroblasts. Comparison of the responses in primary WT MEFs with IRF3^-/-^ or IFNAR1^-/-^ MEFs allowed us to differentiate gene regulation dependent on the activation of IRF3 downstream of PRR activation versus in response to type I IFN signalling downstream of IFNAR1/IFNAR2 activation (Figure 8A). DNA sensing has previously been reported as the most biologically relevant pathway in immune control of initial CMV infection (80). Accordingly, MEFs were stimulated for 4 h by transfection of immunostimulatory DNA (ISD) to detect IRF3-mediated regulation (*Ifnb1*) before production of type I IFNs would upregulate ISG expression, or for 3 h by treatment with murine IFNβ to detect the peak of the first transcriptional response to canonical IFNAR signalling (*Ifit1*, *Rsad2*, *Stat1*; Figure S4A). Co-incubation with 200 µM of the nucleoside analogue 4-thiouridine for 2 h yielded a good incorporation rate (∼5%, Figure S4B, C) and did not change gene expression (Figure S4D, E), and was used in all conditions for metabolic labelling of newly synthesized RNA.

**Figure 8:**
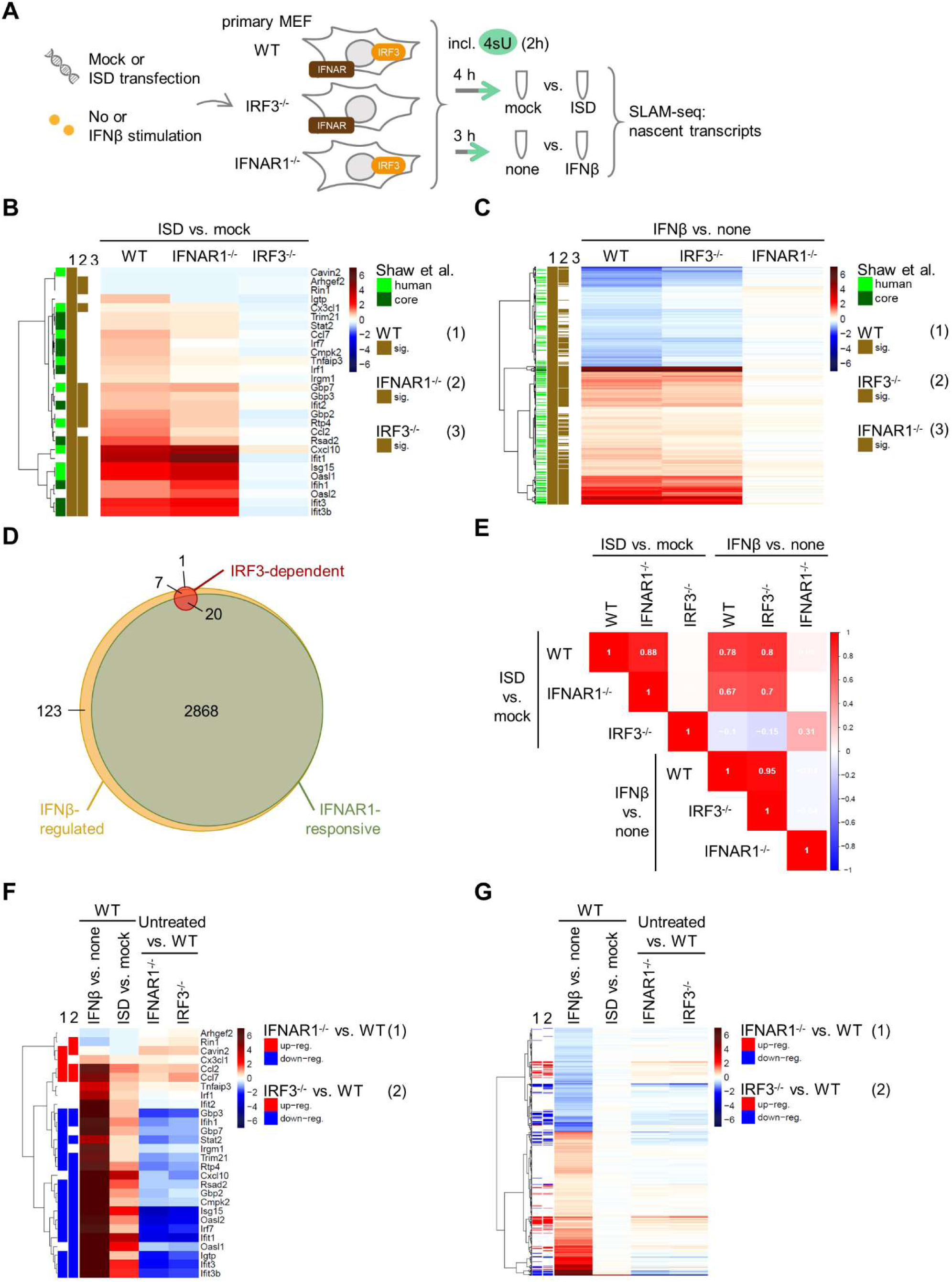
SLAM-seq for characterisation of the dependency of ISD-stimulated transcripts on IRF3 or of IFNβ-stimulated transcripts on canonical type I IFN-IFNAR1/IFNAR2 signalling in MEFs. (A) Determination of IRF3-dependent and IFNAR1-responsive transcripts. Primary MEFs of WT, IRF3^-/-^ or IFNAR1^-/-^ mice were stimulated by transfection of 5 µg/mL ISD for 4 h or mock- transfected, or stimulated with 100 U/mL of murine IFNβ for 3 h or left untreated. Transcripts were labelled in the last 2 h of stimulation by incubation with 200 µM of 4-thiouridine (4sU) and analysed by SLAM-seq. Samples were prepared and analysed in quadruplicate. (B-C) Heatmaps showing the log_2_ fold-changes (log_2_FC; blue: down-, red: up-regulation) in the indicated cell lines for (B) the 28 genes IRF3-dependent genes detected after ISD stimulation or (C) the 2,888 IFNAR1-responsive genes detected after IFNβ treatment. Transcripts with an FDR ≤ 0.01 were considered statistically significant. The green marks on the left indicate overlaps with IFNα- responsive genes in human fibroblasts or conserved (core) between ten different species (81), brown marks show significant regulation in the different cells. Genes were clustered according to Euclidean distances with Ward’s clustering criterion. (D) Venn diagram showing overlaps of genes regulated in an IRF3-specific manner in response to ISD treatment (IRF3-dependent genes), regulated upon IFNβ in WT MEFs (independent of regulation in IFNAR1^-/-^ cells) or regulated by IFNβ only in WT but not IFNAR1^-/-^ MEFs (IFNAR1-responsive genes). (E) Correlation plot showing spearman correlation (blue: negative, red: positive correlation) for pairwise comparisons of log_2_FC for indicated treatments and cell lines for IRF3-dependent genes. (F-G) Heatmaps showing the log_2_FC (blue: down-, red: up-regulation) of (F) IRF3-dependent or (G) IFNAR1-responsive genes in WT cells after IFNβ or ISD treatment compared to controls, and in untreated knockout cell lines compared to WT. Genes significantly differentially expressed (FDR ≤ 0.01) in the (1) IFNAR1^-/-^ or (2) IRF3^-/-^ compared to WT MEFs are marked on the left (blue: down-, red: up- regulation).

In total, 10,616 transcripts were detected across all samples. In response to ISD transfection, 28 transcripts were significantly (FDR ≤ 0.01) up- or down-regulated in WT cells, and consistently, none of these were induced after ISD stimulation of IRF3^-/-^ MEFs (defined as IRF3-dependent genes; Figure S5A). Transcripts of type I IFNs themselves were not detected at sufficient levels for quantification, but we could validate IRF3-dependent induction of *Ifnb1* and *Ifna4* by RT-qPCR (Figure S5C, D). By comparing the response to IFNβ treatment between WT and IFNAR1^-/-^ MEFs, we determined 2,888 transcripts that were significantly up- or down-regulated dependent on type I IFN-mediated IFNAR1/IFNAR2 activation (defined as IFNAR1-dependent type I IFN-responsive [or short: IFNAR1-responsive] genes; Figure S5B). Interestingly, another 130 transcripts were regulated by treatment with IFNβ also in the IFNAR1^-/-^ cells and thus independently of canonical type I IFN signalling (Figure S5B). These 130 transcripts included well-known NF-κB targets such as *Nfkbia*, *Tnfaip3*, and *Cxcl5* (Table S4), highlighting the importance to define ISG induction based on required signalling components, such as INFAR1. Regulation of IRF3-dependent genes upon ISD transfection was comparable between WT and IFNAR1^-/-^ cells (Figure 8B), and vice versa for IFNAR1-responsive genes stimulated by IFNβ treatment between WT and IRF3^-/-^ cells (Figure 8C). In addition, both the IRF3-dependent and the IFNAR1-responsive murine genes overlapped significantly (Fisher’s exact test, p < 0.0001) with IFNα-responsive genes previously determined in human fibroblasts, as well as a conserved ‘core’ of genes in human and nine further vertebrate species (81) (Figure 8B, C).

Next, we examined the IRF3-dependent genes more closely. Comparing the IRF3-dependent and IFNAR1-responsive groups revealed that 20 of the 28 IRF3-dependent genes were also responsive to IFNAR1/IFNAR2 activation (Figure 8D; Fisher’s exact test, p < 0.0001). Moreover, of the remaining 8 IRF3-dependent genes, another 7 responded to IFNβ treatment, though both in WT and IFNAR1^-/-^ cells. Overall, the induction of IRF3-dependent genes was even more pronounced after stimulation via IFNAR activation than via PRR signalling (Figure S5E). Thus, IRF3-dependent genes represent a small subset within the > 100x bigger group of IFNAR1-responsive genes (Figure S5F). Accordingly, expression of IFNAR1-responsive genes is well correlated between IFNβ treatment of WT and IRF3^-/-^ MEFs (Spearman correlation r=0.95), but not between ISD stimulation of WT and IFNAR1^-/-^ cells (r=0.06). In contrast, the regulation of IRF3-dependent gene expression correlated well between ISD stimulation of WT and IFNAR1^-/-^ cells (r=0.88) as well as between IFNβ treatment of WT and IRF3^-/-^ MEFs (r=0.95, Figure 8E).

Furthermore, we observed that the absence of IRF3 or IFNAR1, two key components of the type I signalling system, markedly influenced the basal levels of many transcripts. Interestingly, while signalling via these factors resulted in vastly different numbers of induced genes (28 for IRF3, 2,888 for IFNAR1-responsive activation), similar numbers of transcripts were influenced by both knockouts (1,323 and 1,255 significantly de-regulated transcripts in IRF3^-/-^ and IFNAR1^-/-^, respectively, compared to WT). While the transcriptional profiles in untreated IRF3^-/-^ and IFNAR1^-/-^ cells were distinct from WT cells, expression changes compared to WT were highly similar between the two knockouts (Figure S6A, B). Especially the basal levels of most IRF3-dependent genes were evidently affected by absence of IRF3 or IFNAR1, with most transcripts showing lower basal levels in the IRF3^-/-^ or IFNAR1^-/-^ cells compared to WT cells (Figure 8F). Similarly, the knockouts affected the basal expression of many IFNAR1-responsive genes, again with a similar outcome (Figure 8G). In addition, analysis of the transcripts differentially regulated in IRF3^-/-^ or IFNAR1^-/-^ compared to WT MEFs for enriched biological processes based on gene ontology (GO) indicated that de-regulation of the type I IFN signalling system affects not only immune system processes and the response to stimulation, but also further processes in multicellular organisms like the regulation of cell motility, cell adhesion, and vasculature development (Table S5).

Overall, we identified 28 IRF3-dependent genes in MEFs and observed that almost all of these were also inducible by canonical type I IFN signalling. Of note, the absence of critical components of the type I IFN response greatly impacted the basal levels of many transcripts, including a large fraction of ISGs.

### M35 modulates expression of several IRF3-dependent genes

Having defined the IRF3-dependent and IFNAR1-responsive genes in MEF, we next addressed the effect of M35 on cellular gene expression after PRR stimulation, with special regard to IRF3-driven genes. For this, we generated immortalised MEFs (iMEFs) that constitutively express M35-HAHA. As expected, these cells showed reduced induction of *Ifnb1* transcription upon PRR stimulation compared to an empty vector (EV) control cell line (Figure S7A-D).

Based on the kinetics of *Ifnb1* expression in EV iMEFs upon stimulation with Alexa488-labelled ISD (Figure S7E), cells were transfected with ISD or mock-transfected for 2, 4, or 6 h and analysed by SLAM-seq alongside untreated cells (Figure 9A). Since application of the labelling protocol established before (200 µM of 4sU applied to label RNA for 90 minutes before harvest) did not achieve sufficient incorporation in this experiment, total RNA transcript numbers were analysed instead. As expected from the *Ifnb1* expression kinetic, 2 h was too early to observe a major response (Figure 9B, left panel). To our surprise, there was no striking difference in the gene induction in M35-HAHA compared to EV iMEFs at the peak of *Ifnb1* transcription (Figure S7E) after 4 h of stimulation (Figure 9B, middle panel). Only 6 h after stimulation, several transcripts were more strongly induced in EV compared to M35-expressing cells (Figure 9B, right panel).

**Figure 9:**
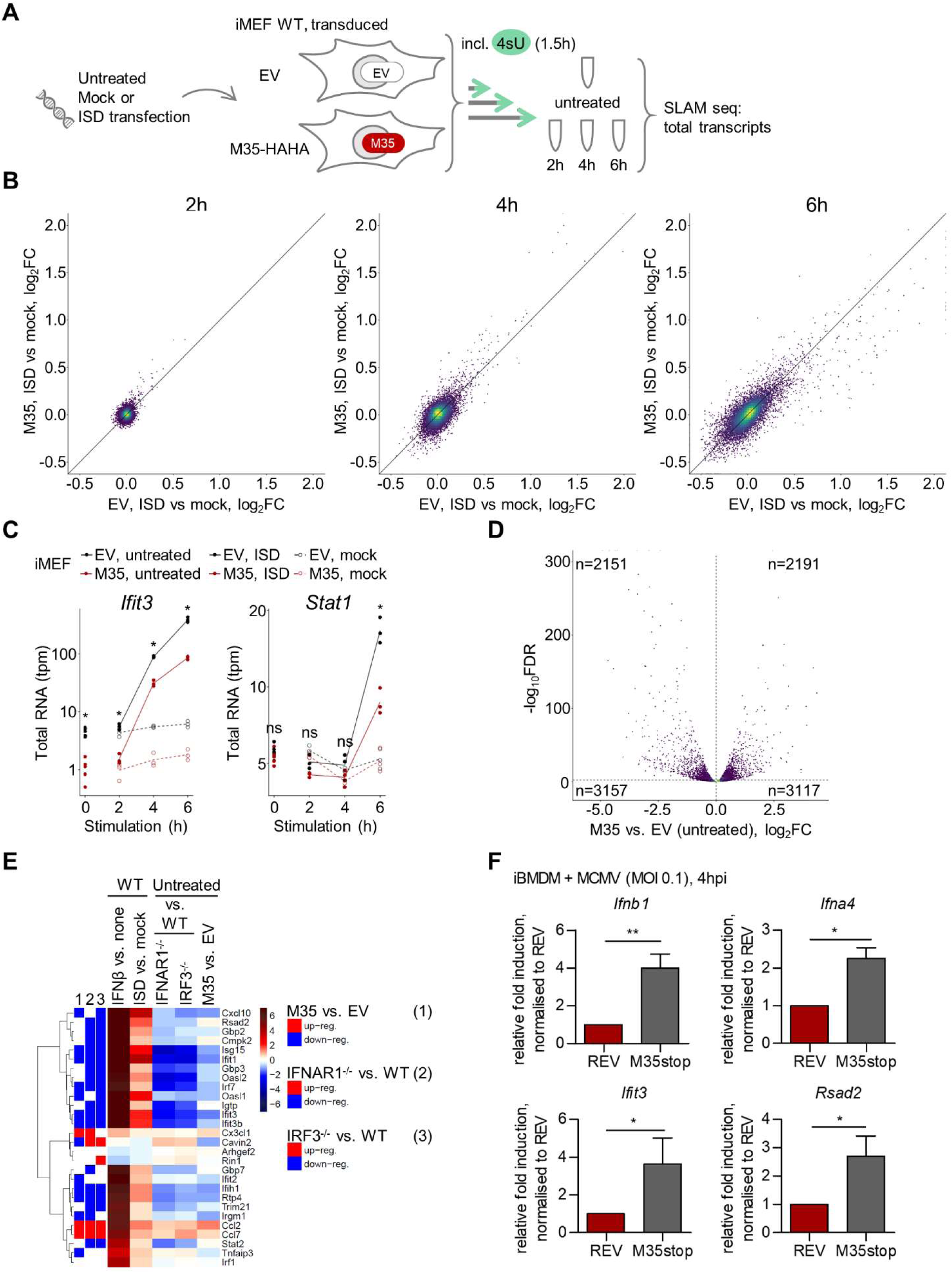
Presence of M35 modulates expression of IRF3-dependent genes. (A) Determination of the global effect of M35’s presence on gene expression. iMEFs stably expressing M35-HAHA or a corresponding EV were stimulated by transfection of 5 µg/mL ISD, mock-transfected or left untreated and incubated for indicated times. Transcripts were labelled in the last 90 min of stimulation by incubation with 200 µM of 4-thiouridine (4sU). Total transcripts were analysed by SLAM-seq. Samples were prepared and analysed in triplicate. (B) Depicted are log_2_FCs of transcripts of EV (x axis) vs. M35-HAHA (y axis) iMEFs after indicated times of ISD stimulation compared to mock-transfection. (C) Expression kinetics of selected transcripts upon PRR stimulation in EV and M35-HAHA iMEFs. Total RNA counts are given in transcripts per million (tpm). Differences between transcript levels in ISD-stimulated EV and M35-HAHA iMEFs with FDR < 0.01 were considered statistically significant, ns non-significant. (D) Volcano plot showing differential expression of total cellular transcripts in EV compared to M35- HAHA iMEFs in untreated conditions as log_2_FC (x axis), plotted against -log_10_ of the FDR (y axis, with significantly (FDR < 0.01) regulated transcripts above the dashed horizontal line). Numbers indicate total up- (log_2_FC > 0) or down-regulated (log_2_FC < 0) transcripts in the respective sections. (E) Heatmaps showing the log_2_FC (blue: down-, red: up-regulation) in the indicated SLAM-seq samples for the 28 IRF3-dependent genes. Genes differentially expressed (FDR ≤ 0.01) in (1) M35-expressing compared to EV iMEFs or in (2) IFNAR1^-/-^ or (3) IRF3^-/-^ compared to WT MEFs are marked at the left (blue: down-, red: up-regulation). (F) Response of IRF3-dependent genes upon infection with MCMV with or without M35. Immortalised BMDMs (iBMDMs) pre-treated with 1 µM ruxolitinib (IFNAR signalling inhibitor) were infected with MCMV M35stopRevertant (REV) or MCMV M35stop (M35stop) at MOI of 0.1 or mock infected. Cells were harvested 4 h post infection for RT-qPCR analysis. Relative fold induction of *Ifnb1*, *Ifna4*, *Ifit3*, and *Rsad2* transcripts was calculated based on the housekeeping gene *Rpl8*, and values were normalised to REV-infected samples. Data is shown as mean ±SD and combined from two (*Ifna4*) or three (*Ifnb1*, *Ifit3*, *Rsad2*) independent experiments. Significance compared to infection with REV was calculated by Student’s *t*-test (unpaired, two-tailed), ns non-significant, * p < 0.05, ** p < 0.01.

We then studied the expression kinetic of individual antiviral genes and found that IRF3-dependent genes such as *Ifit3* were well induced in EV cells after stimulation, as expected, but also in M35-expressing cells (Figure 9C). After 6 h of stimulation, transcription of type I IFN signalling-dependent genes like *Stat1* was upregulated, reflecting activity of IRF3-dependently produced type I IFNs and subsequent IFNAR signalling. Induction of these genes was lower in M35- expressing cells, presumably due to the reduction of type I IFN production in presence of M35. This indicates that on top of the putative direct effect(s) of M35, indirect effects of M35’s antagonism of type I IFN induction contribute to gene regulation after 6 h of stimulation.

Although presence of M35 had no major effect on the fold-change of induction upon stimulation compared to control cells, our analyses revealed that thousands of transcripts already exhibited different basal levels in M35-expressing cells (Figure 9D). 2,151 transcripts were down- and 2,191 transcripts up-regulated in the presence of M35 compared to control cells (FDR < 0.01) even in absence of IRF3 activation. Focusing first on the IRF3-dependent genes due to the proposed antagonism of M35 with IRF3-DNA binding, we found that more than half of the IRF3-dependent genes were differentially regulated in the presence of M35 (16 out of 28 genes, p = 0.06; Figure 9E). Of those, 13 IRF3-dependent genes were significantly down- and three were up-regulated. Remarkably, the pattern of up- or down-regulation of basal expression of IRF3-dependent genes in stably M35-expressing cells was highly similar to the pattern observed in the IRF3^-/-^ or the IFNAR1^-/-^ MEFs (Figure 9E). Accordingly, the transcriptional profile of the IRF3-dependent genes in M35-expressing cells compared to EV cells correlated positively with IRF3^-/-^ and IFNAR1^-/-^ MEFs and negatively with the induction of those genes by ISD transfection (Figure S8A). Regarding the effect of M35’s presence on IFNAR1-responsive genes, we found that M35 caused differential regulation of a significant fraction of these transcripts (overlap of 1,373 genes, p < 0.0001), but the regulation did not correlate with the trends of any of the other tested conditions (Figure S8B, C). Finally, we compared basal levels of IRF3-dependent, IFNAR1-responsive, and all further transcripts in M35-expressing versus EV control cells. This revealed that presence of M35 overall significantly down-regulated IRF3-dependent gene expression, while this tendency was not observed for IFNAR1-responsive or other regulated genes (Figure S8D).

All in all, we observed that the presence of M35 broadly affects basal gene expression in iMEFs, similar to knockouts of IRF3 or IFNAR1, and specifically down-regulates expression of IRF3- dependent genes. This supports the hypothesis that M35 specifically modulates transcription of IRF3-targeted genes aside from *Ifnb1*. To address a direct modulation of IRF3-dependent gene induction by M35 in the infection context, we compared the response elicited by MCMV M35stop, a recombinant that lacks M35 due to introduction of a Stop cassette within the ORF in the viral genome, to the revertant virus (MCMV REV) in which expression of M35 was restored. We infected macrophages, because the effect of M35 on viral replication was best observable in these cells (60), and assessed induction of individual transcripts after 4 h by RT-qPCR. To rule out that type I IFN production and subsequent IFNAR signalling influenced the results, cells were additionally treated with the IFNAR signalling inhibitor ruxolitinib ((82), Figure S9A). As expected, absence of M35 resulted in a higher increase of *Ifnb1* and *Ifna4* transcription early after infection (Figure 9F). Moreover, transcription of the IRF3-dependent genes *Ifit3* and *Rsad2* was significantly increased upon infection with MCMV M35stop compared to MCMV REV. Comparing the fold inductions of the analysed transcripts reflects that the trans-activation of *Ifnb1* is among the strongest responses at this early time point after infection (Figure S9B). Expression of *Stat1* was not detectable after infection, as expected after inhibition of type I IFN signalling. In contrast to IRF3- driven expression, transcription of the NF-κB-dependent genes *Nfkbia* and *Tgfb1* was not affected by the presence or absence of M35 during infection (Figure S9C). This demonstrates that the tegument protein M35 directly antagonised IRF3-mediated gene induction early during MCMV infection, and that this is independent of M35’s inhibition of type I IFN expression and subsequent IFNAR signalling (Figure 10).

**Figure 10:**
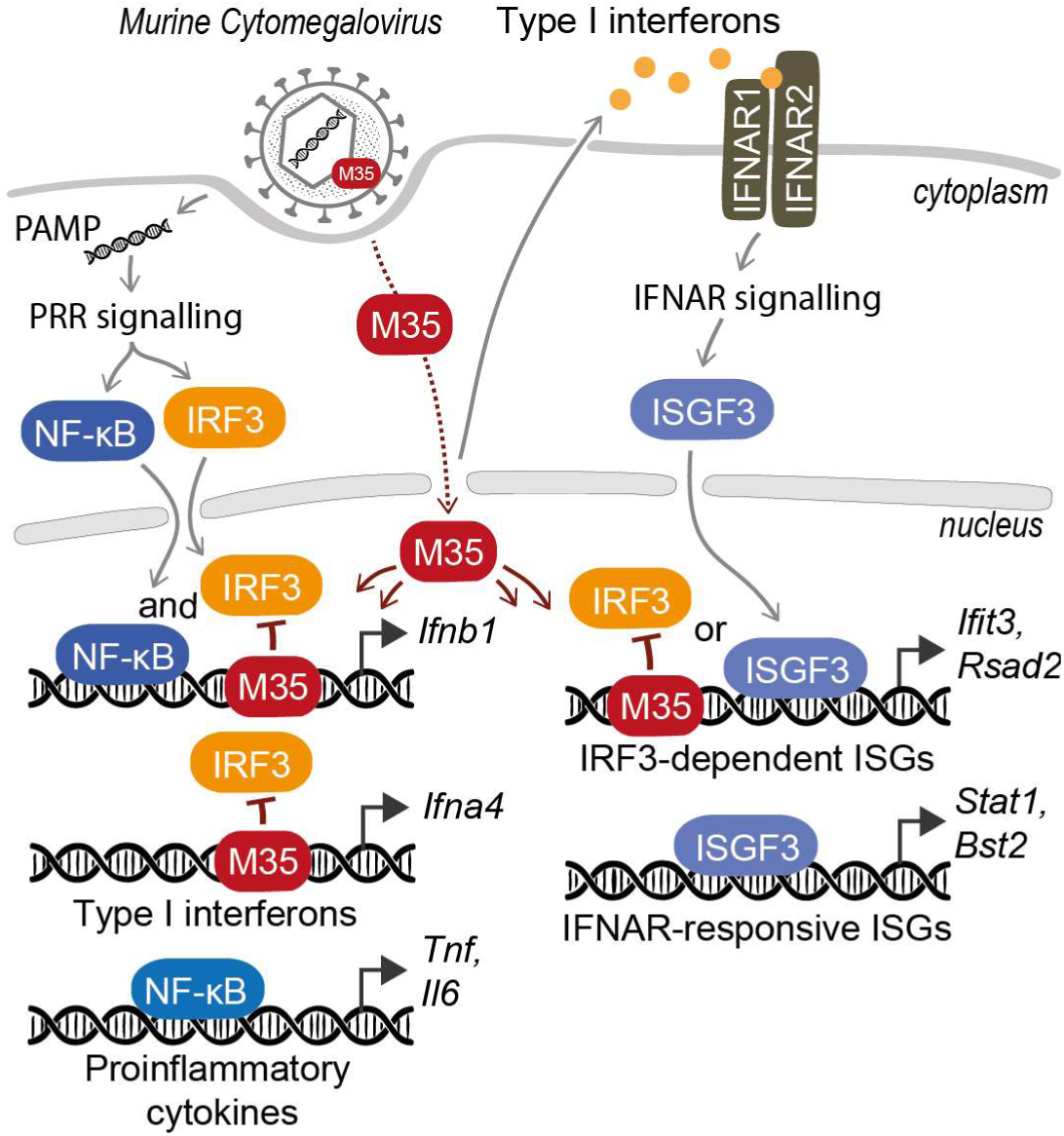
M35 binds to specific host promoters and interferes with IRF3-dependent gene expression. Upon infection of a host cell with MCMV, pathogen-associated molecular patterns (PAMPs) are sensed by pattern recognition receptors (PRRs) and activate the transcription factors NF-κB and IRF3. NF-κB induces expression of proinflammatory cytokines, NF-κB and IRF3 together induce expression of *Ifnb1*, and IRF3 regulates expression of further type I interferons and induces a subset of the interferon-stimulated genes (ISGs). Released type I interferons activate the type I interferon receptor (IFNAR). IFNAR signalling induces assembly of different transcription factors complexes, mainly interferon-stimulated gene factor 3 (ISGF3) which further drives expression of various ISGs. During MCMV infection, the viral tegument protein M35 is released and rapidly shuttles to the nucleus. M35 binds to IRF3-targeted recognition elements in host promoters and thus antagonizes recruitment of IRF3, resulting in inhibition of IRF3-driven gene expression.

## Discussion

To successfully establish persistent infections, members of all herpesvirus subfamilies dedicate a substantial number of gene products to target PRR signalling and induction of type I IFNs (83). We previously identified MCMV M35 as the first CMV antagonist of type I IFN expression and showed its crucial role during infection of the host. In this study, we characterise the structure of the M35 protein and describe the mechanism that this potent immune modulator applies to pave the way for a successful infection.

A previous study of M35’s homologue HHV6B U14 had indicated instability of the full-length proteins due to intrinsically disordered C-termini (66), and in line with this, M35 full-length protein was unstable after purification. Yet, the major N-terminal part of M35 (aa 2-458) could be purified and allowed crystallization. Importantly, this structured domain retained the immunomodulatory activity of M35, demonstrating its contribution to the protein’s function. A loop structure near the β-hairpins was not resolved in the crystal structure of M35, but reverse genetic analysis showed that this loop is not essential for M35’s function. The corresponding segment was resolved in the structure of HHV6B U14 (66), suggesting that the part generally allows crystallisation. Instead, we propose that in the crystal form of M35 obtained here, this region lacked the necessary contacts to keep the loops in an ordered conformation. Combining the data of the crystal structure, SEC-MALS, co-immunoprecipitation, and native PAGE, our findings demonstrate that M35 forms homodimers, and does so independently of experimental conditions. This is in contrast to the prediction by Wang and colleagues, which had suggested that MCMV M35 and HCMV UL35 would not dimerise because they featured markedly different residues at the interface than the *Roseolovirus* U14 proteins (66).

The MCMV M35 crystal structure is highly similar to that of HHV6B U14, but our investigations suggest that the immunomodulatory function is not generally conserved in the members of the pp85 protein superfamily. In line with our observations, a recent study shows that HHV6A U14 does not significantly modulate the activity of the 125 bp human *Ifnb1* promoter in the absence of PRR stimulation, although it interacts with p65 and increases expression of NF-κB dependent promoters (84). Still, down-modulation of *Ifnb1* promoter induction by HHV7 U14 indicates that at least one further homologue might share M35’s immunomodulatory activity. Comparably low protein levels were obtained for all U14 proteins though they were expressed from the same vector as M35. Since the phenotype of HHV7 U14 was observable with both epitope-tagged or untagged variants and despite the low expression level, we conclude that the protein levels very likely do not explain the differences in the putative U14 activities. We recently reported that also the HCMV homologue of M35, UL35, impairs IFNβ production, albeit modulation occurs at a different level than M35 (63). This highlights that while the two CMV homologues assumed related roles to support viral replication, they adapted different strategies to inhibit type I IFN induction. It remains to be determined whether HHV7 U14 influences *Ifnb1* expression using a similar mechanism to M35, or UL35, or even uses a yet unknown way, but this beyond the scope of this study.

Using reverse genetics, we found that the pair of β-hairpins, which seems to interlock the M35 moieties like two thumbs interlock a handshake, is essential for M35’s function. Interestingly, another critical residue, R69, is localised at the opposite side from the β-hairpins, but also directly at the dimer interface. Conspicuously, protein levels of both loss-of-function derivatives (Δβ and R69A) were reduced compared to WT M35 in cell lysates, but since very low amounts of WT M35 protein sufficed to inhibit luciferase induction in our *Ifnb1* reporter assay, this alone would not explain the loss-of-function. The formation of distinct nuclear speckles of M35 Δβ further added to the impression that this loss-of-function mutation corrupted the protein. In contrast, M35 R69A displayed a WT-like nuclear distribution. Remarkably, native PAGE showed that both Δβ and R69A severely impaired dimerisation, revealing that exchange of a single position could critically influence integrity of the M35 dimer. This finding highlighted that dimerisation is an essential feature of M35 and related to the immunomodulatory function.

The purified M35_S protein enabled us to confirm specific binding of M35 to the IFNβ enhancer dsDNA in vitro. At high M35_S concentration, a second band appeared that ran higher than the initial DNA-protein complex. This might indicate (i) the formation of multimeric complexes in which M35-DNA complexes interact with not DNA-bound M35 proteins, or (ii) formation of a higher-order complex formed by binding of several M35 dimers to the DNA, as observed at high concentrations of IRF3 (35). Cooperative binding was also indicated from affinity measurements by the Hill coefficient (h > 1). Assuming that M35 binds to DNA as dimeric entity, and given that no other viral or cellular proteins were present in the reactions, these observations support the possibility of binding of several M35 dimers to one dsDNA strand. The overlapping IREs span about 25 bp in total, and the IRF3/7 dimers are proposed to bind from two sides. Likewise, and in agreement with M35’s binding prerequisites, M35 could bind with two dimers, each from one side of the DNA duplex.

The IFNβ enhancer sequence itself is accessible for transcription factors in steady state (85), supporting a model of direct M35-DNA binding. We determined a K_d_ of 2 µM for binding of M35 to the murine IFNβ enhancer sequence, but since concentrations applied in EMSAs refer to the M35_S monomer, this translates into a K_d_ of about 1 µM for dimeric M35. Based on the EMSA and ChIP, we propose that M35 antagonises binding of IRF3, which displays a similar range of affinity (K_d_ ∼ 1 µM, (86)). However, for successful recruitment to the IFNβ enhancer, IRF3 requires PRR- signalling induced phosphorylation, dimerisation, and interaction with the co-activator CBP/p300 to overcome its intrinsically low DNA affinity (87, 88). Accordingly, with a K_d_ of 6 nM, the phosphomimetic IRF3-5D dimer features a distinctively higher affinity than WT IRF3 (86).

Nevertheless, all of these affinity measurements examined isolated proteins and synthetic dsDNA probes, depriving their interactions of the natural environment. Application of the purified M35 protein gave rise to a specific protein-DNA complex, demonstrating that in general, no host factors or further viral factors are required for M35-DNA binding. Still, host factors, DNA modifications, and the environment of the promoter in the cell could greatly influence the ability and affinity of M35 to bind to DNA and contribute to its efficient recruitment. In this way, the relatively weak intrinsic affinity of M35 in the low µM range could suffice to antagonise binding of endogenous IRF3 after PRR activation. SLAM-seq showed that stable expression of M35 inhibited but did not abolish induction of IRF3-dependent genes, potentially reflecting displacement of promoter-bound M35 by IRF3. Nevertheless, exogenous expression enabled to study the immunomodulatory activity of M35, and ChIP demonstrated that recruitment of IRF3 to the *Ifnb1* promoter was severely impaired in M35’s presence. In the context of infection, viral factors such as the reported interaction partners of M35 may further regulate its immunomodulatory activity (89), and characterisation of their interplay will greatly add to our understanding of CMV immunomodulation.

To formally validate binding of M35 to the host DNA in its native environment, we performed ChIP experiments with the stably M35-expressing cell line or infection of host cells with MCMV REV and MCMV M35stop. However, while M35 protein itself could be precipitated with an epitope tag- or an M35-specific antibody, we could not detect specific enrichment of any host DNA sequences after M35-specific immunoprecipitation by qPCR or unbiased high-throughput sequencing. Detection of transient and low-abundant interactions is a common pitfall in ChIP assays, especially when sample material is limited, and inaccessibility of chromatin-bound M35 for antibodies or changes of the targeted epitope due to formaldehyde cross-linking may prevent this approach from succeeding.

Nonetheless, the DNA-binding ability of M35 opened the possibility that M35 might modulate expression of further genes aside from *Ifnb1*. As an alternative approach, we studied M35’s effect on global gene expression with SLAM-seq to allow the unbiased measurement of transcript levels at a wide dynamic range and with additional temporal resolution by detection of newly synthesized transcripts (78, 90, 91). The herein determined group of ISD-stimulated IRF3- dependent genes is in agreement with those detected upon stimulation of RIG-I signalling in murine cells using mRNA microarrays (23, 92). Likewise, the type I IFN-regulated genes detected in fibroblasts of other species by mRNA sequencing agree with the IFNAR1-responsive group (81). Induction of type I IFN genes could only be quantified by probe-based RT-qPCR, demonstrating its unmatched sensitivity and persisting value for detection of low abundant transcripts. While in total more than one quarter of the detected gene products responded to IFNβ treatment, IRF3 activation regulated only a small group. Moreover, almost all IRF3-dependent genes were also IFNAR1-responsive, underlining the role of IRF3 in priming the induction of a subset of ISGs before type I IFN signalling elicits the full potential of the antiviral response. We also detected IFNAR1-independent IFNβ-stimulated regulation of transcripts commonly attributed to the NF- κB-driven response. This may indicate an impurity in the applied IFNβ, or IFNAR1-independent activation of signalling cascades via IFNAR2 homodimerisation as suggested before (93, 94). Moreover, knockout of IRF3 or IFNAR1 severely affects the signalling circuits of the cell-intrinsic antiviral response, including basal transcription of critical signalling components that are themselves ISGs, such as *Stat1*, *Stat2*, and *Irf9*, consistent with previous reports (92). Accordingly, we based our definitions on the responses of WT cells and compared these to the respective knockouts to exclude unspecific responses.

Expressing M35 in iMEFs revealed that its presence alone greatly changed the cellular transcriptome and systematically modulated expression of IRF3-dependent genes. While this correlated with the changes observed in IRF3^-/-^ or IFNAR1^-/-^ fibroblasts, overexpression of M35 did not simply phenocopy de-regulation of the type I IFN system but has discrete effects. We assume that secondary effects contributed to the transcriptional changes of stably M35-expressing cells, such as down-modulation of signalling components, and at least partly obscured M35’s direct effect on gene expression. Therefore, we concluded our analysis by confirming that independently of the known antagonism of type I IFN signalling, virus-delivered M35 inhibited the induction of IRF3-dependent genes early in MCMV infection.

Taken together, we propose that tegument-delivered M35 directly binds to selected promoters and in this way antagonises binding of IRF3 to overlapping recognition sites. The previous characterisation of M35 had indicated an antagonism of M35 with NF-κB based on (i) M35-mediated down-modulation of induction of an artificial NF-κB luciferase reporter, and (ii) correlation of M35’s phenotype with reduced levels of pro-inflammatory TNFα in macrophages at 16 h post infection. Our latest data demonstrate that the NF-κB motif was not required for M35- DNA binding in EMSA, nor did M35 antagonise *Ifnb1* promoter induction by GFP-p65. Instead, the antagonism of M35 with IRF3 now suggests that the observed modulation of the NF-κB mediated response is a secondary effect of M35-mediated modulation of the type I IFN response. With this and the speed of type I IFN signalling in mind, we ensured monitoring of direct effects for the present study by assaying early time points after PRR stimulation or after infection and additional inhibition of IFNAR signalling during infection.

Interestingly, a similar antagonism of IRF3-promoter binding is also employed by three unrelated proteins from the *Beta-* and *Gammaherpesvirinae*: namely the DNA polymerase subunit UL44 of HCMV (69), the latency-associated nuclear antigen (LANA-1) of KSHV (70), and the transcriptional repressor K-bZIP of KSHV (68). In line with the different evolutionary origins of the proteins, the available information, though limited, does not indicate any structural similarities. Like M35, these three proteins localise to the nucleus, bind to the sequence of the IFNβ enhancer, reduce binding of IRF3 to the *Ifnb1* promoter sequence, and inhibit expression of *Ifnb1* and at least one further IRF3-dependent gene. However, the studies of the other antagonists of IRF3-DNA binding had to rely on ectopically expressed proteins, limiting their characterisation regarding timing of activity and impact on viral fitness. We generated a recombinant MCMV deficient in M35 expression and using this virus enabled more detailed characterisation of M35’s influence during MCMV infection (60). We demonstrated that M35 is necessary for the virus to successfully replicate in cell culture as well as in the host organism (60), and that upon infection, tegument-delivered M35 immediately and directly counters induction of IRF3-dependent transcripts. In addition, by examining the global effect of M35 on cellular gene expression, we uncovered that M35 not only influences the antiviral response by downregulating type I IFNs, but directly affects expression of several IRF3-dependent genes.

Overall, our data illustrate M35 as a specific inhibitor of IRF3-mediated regulation of antiviral genes. We found that by deploying M35, MCMV targets an essential step of the host response and influences the type I IFN response more broadly than anticipated. MCMV also deploys the protein m152 to modulate induction of type I IFNs (61). Remarkably, m152 delays the activation of IRF3 downstream of the DNA sensor cGAS and thus impairs the type I IFN response, but allows activation and signalling of NF-κB to harness its pro-viral benefits. A study by Kropp and colleagues indicated a pro-viral role of activated IRF3 for viral gene expression (95), letting us speculate that by modifying the activation of IRF3 with m152, and IRF3 binding to antiviral host promoters with M35, MCMV could exploit this host transcription factor for its own gene expression. Additional studies will be required to address the interplay and effect of the herpesviral immune modulators in the course of type I IFN signalling during viral infection as well as the potential involvement and regulation of IRF3 and NF-κB in herpesviral gene expression.

## Material and Methods

### Mice for generation of primary cells

Mice (C57BL/6J) were bred at the animal facility of the Helmholtz Centre for Infection Research in Braunschweig and maintained under specific-pathogen-free conditions in accordance with institutional, state, and federal guidelines. IRF3 and IFNAR1 knockout mice have been described (96, 97). Primary mouse embryonic fibroblasts (MEF) from C57BL/6J mice were generated by standard protocol (98).

### Plasmids

pRL-TK, expressing *Renilla* luciferase under control of the thymidine kinase (TK) promoter, is commercially available (#E2241, Promega, Walldorf, Germany). pGL3basic-IFNβ-Luc (*mIfnb1*- FLuc) consists of the 812 bp murine *Ifnb1* promoter region cloned into pGL3basic (Promega) upstream of the firefly luciferase gene (99). The firefly luciferase reporter plasmid p-125 (*hIfnb1*- FLuc), consisting of the human *Ifnb1* promoter region (−125 to +19), was kindly provided by Takashi Fujita (Kyoto University, Japan) (16). pFLAG-CMV-huIPS1 expressing Flag-MAVS was kindly provided by Friedemann Weber (Institute of Virology, Justus Liebig University Giessen, Germany) (100). pCMVBL IRF3-5D encoding constitutively active human IRF3 by containing five amino acid substitutions (S396D, S398D, S402D, S404D, S405D) was kindly provided by John Hiscott (Institut Pasteur Cenci Bolognetti Foundation, Rome, Italy). pEGFP-C1-RelA (GFP-p65, #23255) is available from Addgene (Watertown, MA, USA). pIRES2-GFP (#6029-1), pQCXIH, and pQCXIP (#631516) vectors were purchased from Clontech Laboratories (Mountain View, CA, USA). pcDNA3.1+ empty vector (EV; #V790-20) and pcDNA3.1 TOPO EV (#K480001) are from Invitrogen (Thermo Fisher Scientific).

pEGFP-C1-hIRF3 encoding human IRF3 N-terminally fused with eGFP (eGFP-IRF3) was kindly provided by Friedemann Weber (Institute of Virology, Justus Liebig University Giessen, Germany) (101).

Expression constructs for M35-V5/His, M34-V5/His and M27-V5/His (all in pcDNA3.1 TOPO- V5/His) have been described previously (102). The expression construct pcDNA3.1 TOPO M35_S (short: aa1-452) was generated by PCR amplifying the M35 aa1-452 sequence with a primer pair that introduces overhangs for restriction enzymes, digesting the product with BamHI and EcoRV and ligating it into pcDNA3.1 TOPO EV linearised with BamHI and PmeI.

Constructs for protein production were generated by PCR amplification of the coding sequences of full-length M35 protein (aa 1-519, nucleotides 45,915–47,471 of GenBank accession #GU305914) and of C-terminally truncated M35 (aa 1-452, nucleotides 45,915–47,267 of GenBank accession #GU305914) followed by sequence- and ligation-independent cloning (SLIC, (103)) between the BamHI and AvrII sites of pCAD04, a modified pOpIE2 vector (104) with a Kozak consensus sequence (GGATCACCATGG) in place of pOpIE2’s original BamHI site and an N-terminal Twin-Strep-tag and TEV cleavage site (MASAWSHPQF EKGGGSGGGS GGSAWSHPQF EKSGENLYFQ GS). pOpiE2 contains the promoter of the second immediate early gene of the baculovirus OpMNPV. The resulting plasmids were named pHER08_M35_S_452_NStr (NStr- M35_S) and pHER09_M35_FL_NStr (NStr-M35_FL).

Packaging plasmids VSV-G encoding for the envelop protein of Vesicular Stomatitis Virus and gag-pol encoding for the retroviral polyprotein group-specific antigen (gag) processed to structural proteins and reverse transcriptase (pol) were a kind gift from Boaz Tirosh (The Hebrew University of Jerusalem, Jerusalem, Israel). The pQCXIH M35-myc/His has been described previously (60). For generation of M35 constructs with a C-terminal double HA epitope tag, the HAHA sequence was fused in frame to the full-length M35 ORF by PCR amplification and subcloned into pcDNA3.1(+). M35-HAHA was then cloned into pcDNA3.1 TOPO using HindIII and SacII restriction sites to generate pcDNA3.1 TOPO M35-HAHA with the same upstream backbone as M35-V5/His. To generate the transduction vector pQCXIP M35-HAHA, the M35-HAHA ORF was PCR amplified with a primer pair introducing overhangs for restriction enzymes and the digested product was ligated into linearised pQCXIP vector using AgeI and BamHI sites. The expression construct pcDNA3.1 TOPO M35 (full-length) was generated by PCR amplifying M35 aa1-519 from pcDNA3.1 TOPO M35-V5/His (WT) with a T7fwd standard primer and a reverse primer that introduces a stop codon directly after the M35 ORF followed by a PmeI restriction site, and ligating the digested product into the linearised pcDNA3.1 TOPO EV using BamHI and PmeI restriction sites.

For the generation of U14 expression constructs based on pcDNA3.1 TOPO, the U14 ORFs of HHV6A (strain U1102), HHV6B (strain HST), and HHV7 (strain JI) were ordered as gBlock gene fragments (Integrated DNA Technologies, Leuven, Belgium) with the same upstream and downstream sequences as the M35 ORF in pcDNA3.1 TOPO M35-V5/His including the sequence of the V5/His epitope tag. gBlocks were subcloned into pcDNA3.1 TOPO EV using BamHI and NotI restriction sites to generate pcDNA3.1 TOPO U14-V5/His with the ORF of HHV6A or HHV6B or HHV7. From there, U14 ORFs with N-terminal triple V5 epitope tags (3xV5-U14) were generated by introducing the upstream sequence encoding the epitope tag and a triple GGS linker and a downstream stop codon alongside restriction sites with PCR primers, followed by ligation of the digested product into pcDNA3.1 TOPO using BamHI and PmeI restriction sites. Untagged U14 ORFs (U14) were generated accordingly by amplifying the ORF from the U14-V5/His subclones with a T7fwd standard primer and the reverse primer used for generation of 3xV5-U14. pcDNA3.1+ expression constructs for UL35-HA and untagged UL35 have been described previously (63).

The following M35 derivative constructs were derived from pcDNA3.1 TOPO M35-V5/His using the Q5 site-directed mutagenesis kit (#E0554, New England Biolabs) according to the manufacturer’s protocol: (1) mutations of the β-hairpin: deletion of aa 406-424 (Δβ), deletion/insertion variants replacing aa 406-424 with a single G or P (Δβ+G and Δβ+P, respectively); (2) mutations substituting the residues that form the positive patch: R10A+R20A, R99A+R102A, R257A+R260A, R99A+R102A+R257A+R260A, R10A+R20A+R99A+R102A+R217A+R257A+R260A+R310A; (3) mutations that substitute the hydrophilic residues along the groove: N42A+R69A, K71A+H72A+R73A, H174A+R177A+D180A, K438A+R439A, N42A, R69A; (4) mutations of the loop that is unresolved in M35: deletion/insertion variants replacing aa T343-R375 with GSG or GPG (GSG or GPG, respectively), or replacing L349-K373 with GSGS (GSGS). Indicated positions refer to the protein sequence. Q5 mutagenesis was carried out sequentially to combine several point mutations.

All generated constructs were verified by sequencing. Sequences of primers and constructs are available upon request.

### Cell lines

Mammalian cells were cultured at 37°C in a humidified incubator with 5% or 7.5% CO_2_. M2-10B4 (ATCC #CRL-1972) and human embryonic kidney 293T (HEK293T; ATCC #CRL-3216) cells were obtained from American Type Culture Collection, Manassas, VA, USA (ATCC) and maintained in Dulbecco’s modified Eagle’s medium (DMEM; high glucose) supplemented with 10% foetal calf serum (FCS), 2 mM Glutamine (Gln) and 1% Penicillin/Streptomycin (P/S).

The immortalised WT murine bone marrow-derived macrophage (iBMDM) cell line was obtained through BEI Resources, NIAID NIH (NR-9456) and cultured in DMEM (high glucose) supplemented with 10% FCS, 2 mM Gln, 1% P/S and 50 μM β-mercaptoethanol.

Primary mouse embryonic fibroblasts (MEF) derived from C57BL/6J mice were maintained in MEM supplemented with 10% FCS and 1% P/S. For generation of constitutively M35-expressing stable cell lines, primary MEFs were immortalised with SV40 Large T antigen to generate immortalised MEFs (iMEFs) and maintained in DMEM (high glucose) supplemented with 10% FCS, 1% P/S, 1x non-essential amino acids and 50 µM β-mercaptoethanol. Retroviral particles were generated by co-transfecting a confluent well of a 6-well plate of HEK293T cells with each 1.2 µg of the packaging constructs encoding gag-pol and VSV-G, and with 1.6 µg of the retroviral transduction construct pQCXIH or pQCXIP for stable expression of the respective M35 derivative or with the corresponding empty vector (EV) using Lipofectamine 2000. After 48 h, the culture supernatant was filtered through 0.45 µm syringe-driven filter unit, mixed with polybrene (Santa Cruz Biotechnology, Dallas, US) to reach a final concentration of 8 µg/mL, and added to WT iMEF viral harvest medium (DMEM, 20% FCS, P/S, 10 mM HEPES). Cells were centrifuged for 90 min at 800 g at room temperature, transferred to 37°C for 3 h and then the supernatant was changed to fresh medium. After two days, 250 μg/mL hygromycin or 10 µg/mL puromycin was added to the culture media to select for cells successfully transduced with the pQCXIH or pQCXIP vectors, respectively. This yielded M35-myc/His iMEF and M35-HAHA iMEF constitutively expressing M35 and their respective EV iMEF control cell lines.

The High-Five insect cell line (BTI-Tn-5B1-4, High Five™, Thermo Fisher Scientific) was a kind gift by the Boyce Thompson Institute for Plant Research, Ithaca, USA. High-Five cells adapted to EX- CELL® 405 medium (Sigma-Aldrich, Darmstadt, Germany) were maintained in suspension culture at 27°C (130 rpm) in exponential growth and diluted by passaging to 0.4–0.6×10^6^ cells/mL every 2 or 3 days (104).

### Viruses

Generation of MCMV M35stop and MCMV M35stopREV and preparation of MCMV stocks was reported previously (60). In brief, the MCK-2 repaired genome of MCMV strain Smith carried on a bacterial artificial chromosome (BAC) was manipulated by introduction of a stop cassette (**GGCTAGTTAACTAGCC**) at nucleotide position 46,134 (accession #GU305914) within the M35 ORF to yield the genome of MCMV M35stop. A revertant of MCMV M35stop (REV) was generated by restoring the WT sequence and thus expression of the M35 ORF. M2-10B4 cells were transfected with the BAC DNA using JetPEI for reconstitution of MCMV M35stop (M35stop) and MCMV M35stopREV (REV), respectively. A single clone of each recombinant virus was expanded on M2-10B4 cells and virus from supernatants was concentrated and purified on a 10% Nycodenz cushion. Titres of virus stocks were determined by standard plaque assay on M2-10B4 cells.

### Antibodies and reagents

Generation of the M35-specific monoclonal antibody M35C.01 (α-M35) was described previously (60). Murine anti-V5-tag (clone 7/4, #680602) and rabbit anti-V5-tag (Polyclonal, #903801) antibodies were purchased from BioLegend (San Diego, CA, USA). Anti-V5-tag mAb-magnetic beads (#M167-11) were purchased from MBL International (Woburn, MA, USA). Anti-myc-tag (clone 9E10, #05-419) was purchased from Merck Millipore (Darmstadt, Germany). Rabbit anti- IRF3 antibody (polyclonal, #A303-383A) and rabbit IgG isotype control (#120-101) for ChIP experiments were purchased from Bethyl Laboratories (Montgomery, TX, USA). Murine anti-myc-tag (clone 9B11, #2276), rabbit anti-phospho-IRF3 (clone 4D4G, Serine 396, #4947), rabbit anti-fibrillarin (clone C13C3, #2639), anti-GAPDH (clone 14C10, #2118), rabbit anti-HA-tag (clone C29F4, #3724) antibodies for immunoblots were purchased from Cell Signaling Technology (Frankfurt am Main, Germany). Mouse anti-β-actin (clone AC-15, #A5441) antibody was purchased from Sigma-Aldrich. HRP-coupled GFP-antibody (clone B-2, #sc-9996 HRP) was purchased from Santa Cruz Biotechnology (Heidelberg, Germany). Anti-mouse and anti-rabbit HRP-conjugated or Alexa Fluor488-conjugated secondary antibodies were purchased from Dianova (Hamburg, Germany) and Invitrogen (Thermo Fisher Scientific), respectively.

High molecular weight poly(I:C) (#tlrl-pic) was purchased from Invivogen (San Diego, CA, USA). Interferon-stimulatory DNA (ISD) was generated by mixing the complementary forward (ISD45 bp-for: 5’-TACAGATCTACTAGTGATCTATGACTGATCTGTACATGATCTACA) and reverse (ISD45 bp-rev: 5’-TGTAGATCATGTACAGATCAGTCATAGATCACTAGTAGATCTGTA) 45 bp oligonucleotides, heating to 70°C for 10 min followed by annealing at room temperature. For preparation of Alexa488-labelled ISD, the forward oligonucleotide was ordered with a 5’- Alexa488 conjugate and processed in the same way.

The transfection reagents Lipofectamine 2000 (#11668019, Invitrogen, Thermo Fisher Scientific), FuGENE HD (#E2312, Promega, Walldorf, Germany), and linear polyethylenimine (PEI, 25K, #23966-100, Polysciences, Warrington, PA, USA) were purchased from Life-Technologies, Promega, and Polysciences, respectively. JetPEI was obtained from Polyplus (Illkirch, France). Gibco™ Opti-MEM, DMEM and further additives for cell culture media were obtained from Thermo Fisher Scientific. Protease inhibitors (PI, cOmplete, #4693116001) and phosphatase inhibitors (PhI, PhosSTOP, #4906837001) were from Roche (Mannheim, Germany). Recombinant murine IFNβ (#12405-1) was ordered from PBL Assay Science (Piscataway, NJ, USA) and Ruxolitinib (IFNAR inhibitor, dissolved in DMSO, #S1378) from Selleck Chemicals GmbH (Absource Diagnostics, Munich, Germany).

### Production and purification of recombinant proteins

Full-length M35 and the M35_S were produced by transient transfection of High-Five insect cells with the respective pHER plasmids, followed by purification of two steps of affinity chromatography. 1 L High-Five insect cell culture was transfected using PEI as described (105), resulting in about 25 g cell pellet (wet weight). The cells were resuspended in 50 mL lysis buffer (50 mM Tris pH 7.4, 0.5 M NaCl, 10% glycerol, 1 mM TCEP, 0.5% (v/v) IGEPAL CA-630) after addition of 1 µL Benzonase (25U/µL) and 1 tablet of PI and lysed by vortexing and repeated shearing by pressing the extract with a syringe through a needle of 0.9 mm diameter. Subsequently, the extract was cleared by two runs of centrifugation for 20 min at 16,000 g in a Sorvall F18-12x50, rotor (Thermo Fisher Scientific). The soluble protein fraction was filtered through a 0.45 µm filter. First, the tagged NStr-M35_FL or NStr-M35_S protein were purified by StrepTactin Superflow high capacity (IBA Lifesciences, Göttingen, Germany) chromatography with a 1 mL self-made column (Mobicol, MoBiTec GmbH, Göttingen, Germany) after preincubation in batch for 2 h at 4°C with the column material (primary purification). The column was rinsed with a wash buffer (50 mM Tris pH 8.0, 0.5 M NaCl, 10% glycerol, 5 mM β-mercaptoethanol). For elution, 10 mM desthiobiotin was added to the wash buffer and eluates were collected at a flow rate of 1 mL per minute in 0.5 mL fractions. The eluted fractions were analysed by SDS-PAGE and stained with InstantBlue® Coomassie protein stain. The eluted protein samples were pooled and digested overnight at 4°C using TEV-protease (2 mg/mL) at a ratio of 1:10 (TEV-protease:M35 protein). For the second and final purification, the untagged M35_FL or M35_S protein were purified on a Superdex 200 (26/60) column (Cytiva, Freiburg, Germany) using storage buffer (50 mM Tris pH 7.4, 0.25 M NaCl, 10% glycerol, 1 mM DTT).

### Luciferase-based reporter assays

To study induction of the *Ifnb1* promoter in the luciferase reporter assay, specific components of the signalling cascade were ectopically expressed to mimic pathway activation from a known level. For all reporter assays, 25,000 HEK293T cells were seeded in 96-well plates in 100 µL of culture medium per well and transfected on the following day. All samples were transfected and measured in technical duplicates.

MAVS-stimulated assay, murine *Ifnb1-*reporter: Cells were transiently transfected with 10 ng Flag-MAVS (stimulated) or pcDNA3.1(+) (unstimulated) together with 100 ng *mIfnb1*-FLuc, 10 ng pRL-TK, and 100 ng expression plasmid for the protein of interest complexed with 0.75 µL FuGENE HD in 10 µL Opti-MEM per well.

MAVS-stimulated assay, human *Ifnb1-*reporter: Cells were transiently transfected with 10 ng Flag-MAVS (stimulated) or pcDNA3.1(+) (unstimulated) together with 50 ng *hIfnb1*-FLuc, 10 ng pRL-TK, and 100 ng expression plasmid for the protein of interest complexed with 0.75 µL FuGENE HD in 10 µL Opti-MEM per well.

IRF3-5D-stimulated assay: Cells were transiently transfected with 60 ng pIRF3-5D (stimulated) or pIRES2-GFP (unstimulated) together with 100 ng *mIfnb1*-FLuc, 10 ng pRL-TK, and 120 ng expression plasmid for the protein of interest complexed with 1.0 µL FuGENE HD in 10 µL Opti-MEM per well.

p65-stimulated assay: Cells were transiently transfected with 20 ng GFP-p65 (stimulated) or pcDNA3.1(+) (unstimulated) together with 100 ng *mIfnb1*-FLuc, 10 ng pRL-TK, and 200 ng expression plasmid for the protein of interest complexed with 1.1 µL FuGENE HD in 10 µL Opti-MEM per well.

For all luciferase assays, cells were lysed in 50 µL of 1x passive lysis buffer (Promega) per 96-well at 20 h post transfection. Luciferase production was measured with the Dual-Luciferase® reporter assay system (Promega, #E1980) at a Tecan Infinite® 200 Pro microplate luminometer (Tecan, Männedorf, Switzerland) with signal integration over 2000 ms. Fold induction of Firefly luciferase was calculated by dividing Firefly luciferase values through *Renilla* luciferase values for normalisation and then dividing obtained values from stimulated samples by the corresponding values from unstimulated samples.

### Electrophoretic mobility shift assay (EMSA)

Complementary 5’-biotinylated oligonucleotides pairs harbouring the human or murine IFNβ enhancer sequence and corresponding mutated sequences (see Table 1) were purchased from Integrated DNA Technologies (Leuven, Belgium). Sense and antisense oligonucleotides were annealed together at a 1:1 molar ratio in water at 95°C for 5 minutes, with the temperature decreasing 1°C per minute until the corresponding melting temperature (T_M_) of the oligonucleotide pair (73°C) was reached, held for 30 minutes, followed by another 1°C/minute decrease cycle until 4°C. Reactions were carried out according to the manufacturer’s instructions with the Gelshift™ Chemiluminescent EMSA Kit (#37341, Active Motif, Waterloo, Belgium). Purified M35_S protein was diluted in storage buffer to reach the indicated concentration (0.1 to 10 µM, referring to the 50.1 kDa M35_S monomer) and incubated in 1x kit binding buffer supplemented with 50 ng/μL poly d(l:C), 0.05% (v/v) Nonidet P-40, 5 mM MgCl_2_, 1 mM EDTA, 50 mM KCl, and 3 µg BSA together with 2 fmol of indicated biotinylated oligonucleotides. For competitive EMSA reactions, 200 fmol competitor (non-biotinylated oligonucleotides) was added. Reactions were incubated for 30 minutes on ice. The sample was separated into DNA-protein complexes and free probes by electrophoresis of samples mixed with provided 5x loading dye on a 6% native polyacrylamide gel in 0.5 × TBE containing 2.5% glycerol at 4°C. EMSA gels were pre-run for at least 30 minutes at 4°C prior sample loading. Biotinylated DNA was transferred onto a nylon membrane (Amersham Hybond N+, #RPN203B, Cytiva, Freiburg, Germany) at 380 mA for 40 minutes at 4°C. Membranes were fixed with 120 J/cm^2^ UV-B irradiation using a Bio-Link® BXL Crosslinker (Vilber Lourmat, Eberhardzell, Germany). Blocking, washing, and detection were performed using the Gelshift™ Chemiluminescent EMSA Kit according to the manufacturer’s instructions. Membranes were imaged using a ChemoStar ECL Imager (INTAS, Göttingen, Germany).

**Table 1:**
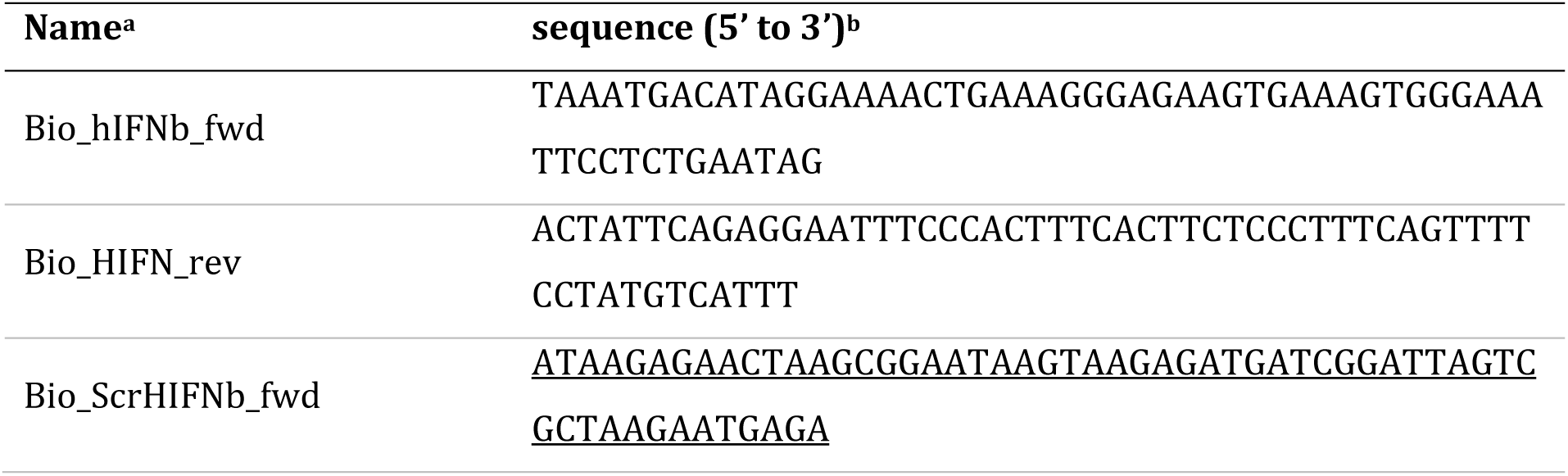
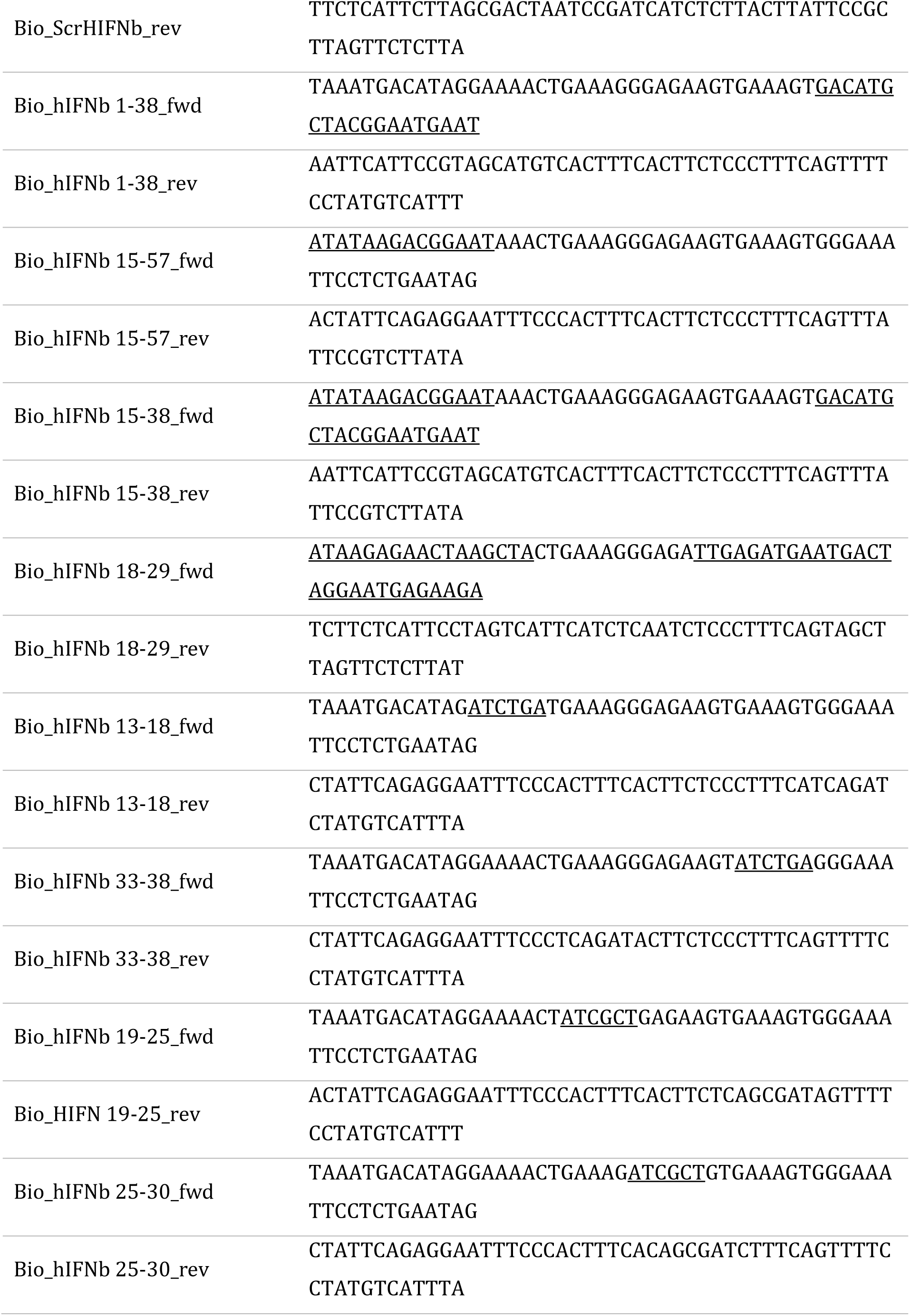
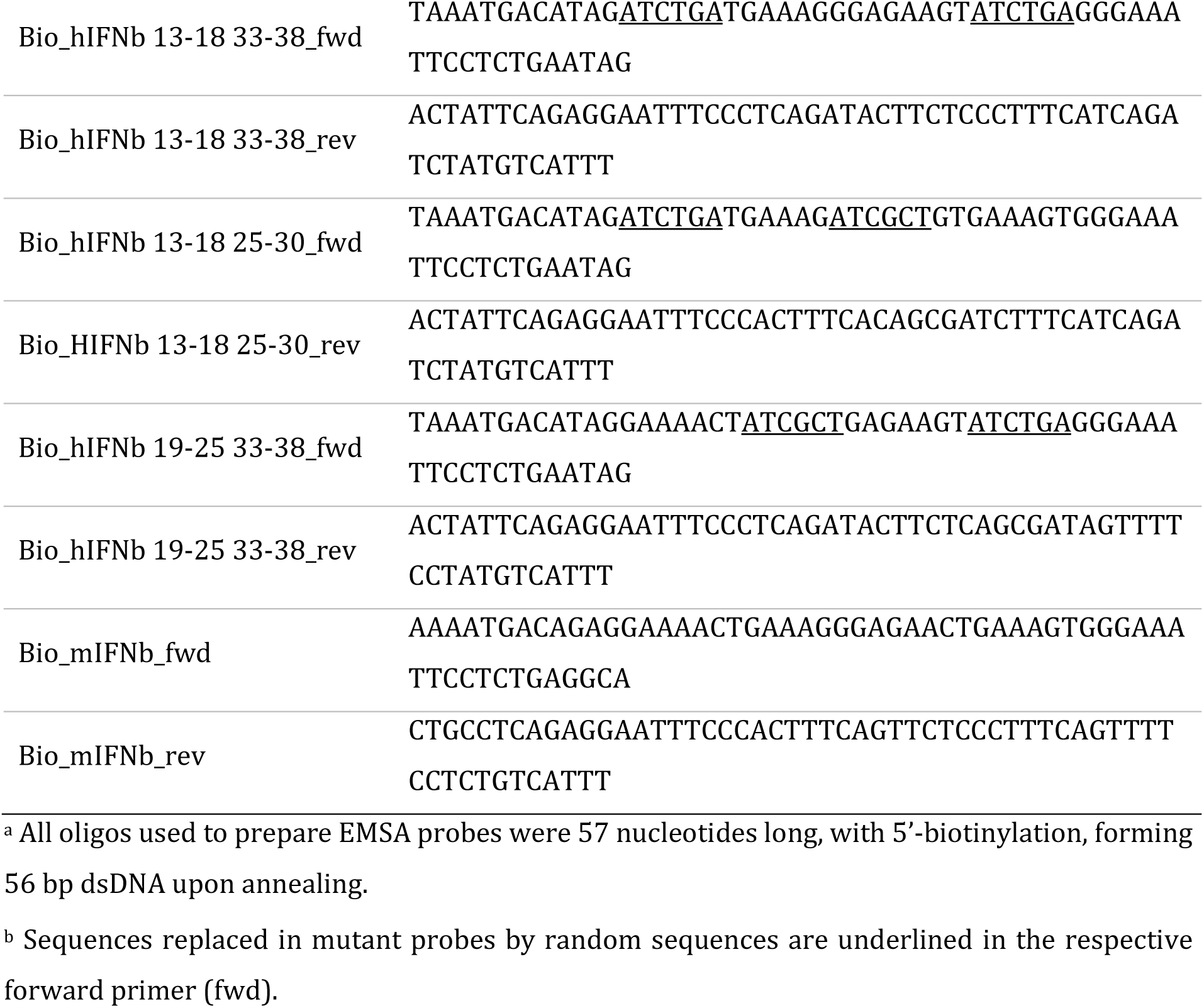
Oligonucleotides used to generate EMSA probes.

For determination of the bound probe fraction, bands of the free probe and the complexed probe were quantified for each replicate using Fiji (version 1.53f51, (106)). The relative band intensities of the complexed probe were divided by the total signal of the free and complexed probes to obtain the bound fraction. The bound fractions determined in three independent experiments were plotted against the protein concentration in GraphPad Prism (version 5; GraphPad Software, San Diego, CA/USA), and fitted using the Binding-Saturation module “Specific binding with Hill slope”.

### Protein crystallography

Structure determination by protein crystallography followed standard protocols. Briefly, initial crystallisation conditions were identified with automated procedures using the sitting drop vapor diffusion method. Crystallisation experiments were performed at room temperature. Conditions for crystallisation of M35_S were determined using a NeXtal JCSG+ matrix screen (#130920, NeXtal, Holland, OH, USA) and M35_S (aa1-452 of 519) readily crystallised in well F10 (1.1 M Na_2_ Malon, 0.1 M HEPES pH 7.0, 0.5% (v/v), Jeffamine ED-2001). Therefore, equal amounts of precipitant solution were mixed with M35_S (3.6 mg/mL in 50 mM Tris pH 7.4, 10% (v/v) glycerol, 250 mM NaCl and 1mM DTT) and incubated at 19°C, yielding a single crystal. Diffraction data of the flash-cooled crystal (cryoprotected with 12% (v/v) 2,3-butanediol) were collected at 100K on beamline P11 of the PETRAIII synchrotron (107) and reduced with autoPROC (108) and STARANISO (109) for scaling. Phasing was achieved by molecular replacement in PHASER (110) using PDB entry 5B1Q as a search model. Refinement involved alternating rounds of manual adjustments in COOT (111) and minimisation with phenix.refine of the PHENIX software suite (112). Data collection and refinement statistics are listed in Table S2. Figures have been prepared with PyMOL (113).

### Size-exclusion chromatography combined with multi-angle light scattering (SEC-MALS)

Experiments were performed on an Agilent 1260 Infinity II HPLC system equipped with a Superdex 200 Increase 10/300 column (Cytiva), a miniDAWN TREOS MALS detector, and an Optilab T-rEX 505 refractometer (Wyatt Technology Corp., Santa Barbara, CA, USA). The column was equilibrated in 50 mM Tris pH 7.4, 10% (v/v) glycerol, 250 mM NaCl and 1mM DTT. 100 µg of protein was injected and SEC-separated on the system. Data were processed with the Astra software package (Wyatt Technology Corp.).

### Immunoblotting

Standard Tris-glycine buffer chemistry (25 mM Tris-base, 192 mM glycine, pH 8.3) was applied for SDS as well as native PAGE and wet transfer. SDS-PAGE used 10% polyacrylamide gels with 0.1% SDS that were run in Tris-glycine running buffer with 0.1% SDS, and gels were blotted in Tris-glycine transfer buffer with 0.05% SDS and 20% methanol for 1 h at 350 mA.

For analysis of protein levels in luciferase assay samples, lysates of technical duplicates were pooled and centrifuged at 11,000 rpm for 10 min to spin out debris. Supernatant was mixed with 4x SDS loading buffer (0.25 M Tris-HCl pH 6.8, 40% glycerol, 8% SDS, 0.04% bromophenol blue, 10% β-mercaptoethanol in H_2_O) and boiled at 95°C for 10 min, then 10 or 15 µL SDS sample was subjected to denaturing SDS-PAGE, followed by blotting on a nitrocellulose membrane (Amersham™ Protean™ 0.45 µm NC, #10600002, Cytiva) as above. Protein transfer for aliquots of the EMSA reactions run on EMSA gels was performed equally. For analysis of chromatin samples prepared for ChIP, aliquots of the prepared chromatin were subjected to SDS-PAGE and blotted on PVDF membrane (Amersham™ Hybond™ P 0.2 µm PVDF, #10600021, Cytiva).

For co-immunoprecipitation of M35-HAHA with M35-V5/His, 800,000 HEK293T cells were seeded and transfected the next day with 4 µg total DNA complexed with 15 µL PEI diluted in a total volume of 300 µL Opti-MEM. 24 h post transfection, cells were lysed in RIPA lysis buffer (20 mM Tris-HCl pH 7.5, 100 mM NaCl, 1 mM EDTA, 1% Triton X-100, 0.1% SDS, 0.5% Na-deoxycholate) freshly supplemented with PI and incubated for 1 h at 4°C on a rotator. 10% of the lysate was used as input control, the remainder was pre-cleared by co-incubation with 40 µL of PureProteome Protein A/G magnetic beads (Merck Millipore, LSKMAGAG10) per sample and rotation for 1 h at 4°C. Supernatant was incubated for 1 h with 50 µL anti-V5-tag mAb-Magnetic beads (#M167-11, MBL) blocked with 1 mg/mL BSA (#B9000S, New England Biolabs). Beads were washed seven times with lysis buffer, and bound protein was eluted by resuspending beads in 1x SDS loading dye in lysis buffer and incubation for 10 min at 95°C. One third of the IP samples and one fourth of the 10% input samples was subjected to SDS-PAGE and immunoblot for analysis.

For validation of M35-HAHA expression in stable cell line, 150,000 cells were washed with PBS, lysed in 100µL RIPA lysis buffer freshly supplemented with PI for 20 min and 15 µL were mixed with 4x SDS loading buffer, incubated as above and subjected to SDS-PAGE and immunoblotting.

For native PAGE (adapted from (114)), 150,000 HEK293T cells were seeded and transfected the next day with 405 ng total DNA complexed with 1.6 µL FuGENE HD in 30µL Opti-MEM per well. Cells were lysed 20 h post transfection in 75 µL RIPA lysis buffer freshly supplemented with PI and incubated for 1 h at 4°C. Debris was pelleted by centrifugation, sample supernatant was mixed with 2x native loading buffer (125 mM Tris-HCl, pH 6.8, 60% glycerol, 0.2% bromophenol blue) and loaded on pre-run native gels with 5% polyacrylamide stacking gel and 7.5% separating gel. Native gels were run at 4°C at 10 mA per gel using Tris-Glycine buffer with 0.2% sodium deoxycholate as cathode buffer and standard Tris-Glycine buffer as anode buffer until the dye ran out. Separated samples were transferred in cold Tris-Glycine buffer for 1 h at 350 mA on a nitrocellulose membrane, then the membrane was incubated for 15 min in fixation solution (40% ethanol, 7% acetic acid, 3% glycerol in H_2_O) at RT and washed 3x in PBS before developing as described below.

After transfer or fixation, membranes were blocked with 5% BSA in TBS with 0.1% Tween-20 (blocking solution), followed by incubation with primary and secondary HRP-coupled antibodies diluted in blocking solution. Membranes were developed with Lumi-Light (#12015200001, Roche) or Pierce™ ECL (#32106, Thermo Fisher Scientific) Western Blotting substrates and imaged on a ChemoStar ECL Imager (INTAS).

### Multiple sequence alignment

Members of the *Betaherpesvirinae* were selected based on the master species list of the International Committee on Taxonomy of Viruses ((115), accessed 24.2.2021 at https://ictv.global/msl, list version 2018b.v2), including the type species of every genus, at least one virus species per family of host organisms, all murine and human members relevant for our comparison, and the bat herpesvirus recently suggested to belong to this subfamily (116). Sequences of the U14 proteins of HHV6A, HHV6B and HHV7 were taken from the same virus strains as in the previous comparison (66). Sequences were aligned online with Clustal Omega (117, 118) (accessed 24.04.2021 at https://www.ebi.ac.uk/Tools/msa/clustalo/). Based on the generated alignment, a phylogenetic tree was created and visualised using the software MEGA-X (119) (Version v.10.2.5). The alignment was illustrated in Jalview (120) (version 2.11.2.5) by highlighting amino acids by characters (colour setting: Clustal X) and by conservation (shade). The percent amino acid (% aa) identities of the virus proteins compared to MCMV M35 were calculated based on the optimal global or local alignment in the online EMBOSS tools Needle and Water (118), respectively (both accessed 15.09.2022).

### Immunofluorescence assay

To characterise M35 derivatives, respective expression constructs were transfected in HEK293T cells in a setup comparable to the luciferase reporter assays. For this, acid-washed glass coverslips (12 mm) were placed in the wells of a 24-well plate and coated by covering with poly-D lysine solution (100 mg/mL in H_2_O) for 15 min. Coverslips were then washed 3x with PBS and 50,000 HEK293T cells were seeded in culture medium. The next day, 405 ng of the plasmid of interest was mixed with 1.6 µL FuGENE HD in 30 µL Opti-MEM per well, incubated for 15 min and added dropwise to conditioned medium. Cells were permeabilised 24 h post transfection by incubation in ice-cold methanol for 5 min at −20°C followed by fixation with 4% PFA in PBS for 20 min at room temperature. Cells were washed three times with PBS and then incubated in IF blocking solution (10% FCS, 1% BSA in PBS) for 1 h at room temperature. Blocked coverslips were incubated with the primary antibody diluted in 1% BSA in PBS overnight at 4°C, followed by three washes with PBS and incubation with secondary antibody and Hoechst (1:500; #33342, Thermo Fisher Scientific) in 1% BSA in PBS for 45 min at room temperature. Coverslips were mounted on glass slides with Prolong Gold (#P36930, Invitrogen, Thermo Fisher Scientific). Imaging was performed on a Nikon ECLIPSE Ti-E-inverted microscope equipped with a spinning disk device (Perkin Elmer Ultraview, Perkin Elmer, Hamburg, Germany), and images were processed using Volocity software (version 6.2.1, Perkin Elmer).

To characterise the stably expressing M35-HAHA compared to EV iMEFs, 150,000 iMEFs were seeded per well of a 12-well plate, allowed to settle for about 6 h, and processed in the wells as described above. Imaging was performed with an EVOS FL cell imaging system (Thermo Fisher Scientific).

### Chromatin immunoprecipitation (ChIP)

To prepare samples for ChIP, 2.5×10^6^ EV or M35-myc/His iMEFs were seeded in 10 cm dishes. After settling for 6 h, poly(I:C) was diluted in Opti-MEM and mixed with diluted Lipofectamine 2000 (1:1), incubated for 20 min at RT, mixed into fresh medium and applied to the cells to obtain a final final concentration of 10 µg/mL. Control cells were mock-treated with Opti-MEM.

After 6 h, formaldehyde (16%, #28908, Thermo Fisher Scientific) was added directly into the culture medium to yield 1% final concentration and incubated for 10 min at RT. To quench, the fixation medium was aspirated, replaced by cold PBS with 0.125 M glycine and incubated for 5 min at RT. Following, samples were processed on ice. Cells were washed 3 times for 10 min with cold PBS, collected by scraping and pelleted at 800 g for 5 min at 4°C. Supernatant was aspirated and pellets were snap-frozen. Pellets were resuspended in 900 µL L1 buffer (50 mM Tris-HCl pH 8.0, 2 mM EDTA, 0.1% Nonidet®P40 substitute (#74385, Fluka), 10% glycerol) freshly supplemented with PI and PhI and incubated for 5 min on ice to lyse the cell membrane. Nuclei were pelleted by centrifugation at 4°C for 5 min at 3000 g and resuspended in 300 µL L2 buffer (50 mM Tris-HCl pH 8.0, 5 mM EDTA, 1% SDS) freshly supplemented with PI and PhI. Chromatin was sonicated in 1.5 mL TPX microtubes (#C30010010, Diagenode, Seraing, Belgium) for 20 cycles (30 sec on/30 sec off, high-intensity) in a Bioruptor NextGen (Diagenode). Chromatin and DNA samples were processed in DNA LoBind tubes from here on (Eppendorf #2023-04-28; #2023-01-28). For control, aliquots of the chromatin (5%) were subjected to SDS-PAGE and immunoblotting as described above.

Per ChIP sample, 10 µg of chromatin were diluted with 9 volumes of ChIP dilution buffer (50 mM Tris-HCl pH 8.0, 5 mM EDTA, 200 mM NaCl, 0.5% Nonidet®P40 substitute), and 1% was removed and purified as input. 50µL Dynabeads™ Protein G Magnetic Beads (#10007D, Invitrogen, Thermo Fisher Scientific) were coupled with 5 µg of the indicated antibody by incubation for 10 min at RT, added to the chromatin and incubated overnight rotating at 4°C. After 16 h, samples were washed in 1 mL of the following buffers freshly supplemented with PI and PhI for each 5 min rotating at 4°C: 1x in ChIP dilution buffer, 3x in high salt washing buffer (20 mM Tris-HCl pH8.0, 0.1% SDS, 2 mM EDTA, 1% Nonidet®P40 substitute, 500 mM NaCl), 1x in LiCl washing buffer (10 mM Tris-HCl pH8.0, 0.25M LiCl, 1 mM EDTA, 1% sodium deoxycholate, 1% Nonidet®P40 substitute), 2x in TE buffer (10 mM Tris-HCl pH8.0, 1 mM EDTA). Lastly, supernatant was discarded, beads resuspended in 100 µL elution buffer (1% SDS, 0.1 M NaHCO3) and incubated for 15 min shaking at 65°C. Supernatant was collected and beads eluted with another 100 µL elution buffer. Eluates were combined and incubated over night at 65°C. Input samples were filled up to 200 µL with elution buffer and processed alongside ChIP samples. The next day, 4 µL of 10 mg/mL RNase A (DNeasy Blood &Tissue kit, #69504, Qiagen, Hilden, Germany) was added to each sample and incubated at 37°C for 2 h. Then 2 µL of 20 mg/mL Proteinase K (#3115879001, Roche) was added and again incubated at 55°C for 2 h. DNA was purified using the NucleoSpin Gel and PCR Clean-up (#74609.250, Macherey-Nagel, Düren, Germany) kit with NTB binding buffer (#740595.150, Macherey-Nagel) according to the manufacturer’s instructions and eluted in 30 µL.

For analysis, 1 µL per input or ChIP sample was used in a qPCR using the GoTaq® Mastermix (#M7133, Promega) and primer pairs for the amplification of the *Ifnb1* promoter region (ChIP_IFNb1_fwd 5’-GCCAGGAGCTTGAATAAAATG, and ChIP_IFNb1_rev 5’-GATGGTCCTTTCTGCCTCAG) or the *Il6* promoter region upstream of the predicted IRF3 binding site (ChIP_Ctrl_IL-6_fwd 5’-CTAGGTACTTCCCTGCAGCC, and ChIP_Ctrl_IL-6_rev 5’-ACCTGCAAACTGGCAAATCG) as control. Enrichment was calculated by the percent input method, where % input = 2^((Cq(input)-Log_2_(dilution factor))-Cq(ChIP sample))x100.

### Stimulation

The timepoint for analysis of upregulated transcripts after IFNβ treatment was determined in a small kinetic experiment. 350,000 WT MEFs cells were seeded per well of a 6-well plate in the evening and stimulated the next morning by diluting IFNβ in Opti-MEM and mixing the pre-dilution into the conditioned medium. A parallel sample was transfected with 5 µg/µl of ISD with Lipofectamine 2000 as described above; an untreated sample served as control. IFNβ-treated cells were harvested after 1, 2, 3 or 4 h, ISD-treated or untreated cells after 4 h, and samples were analysed by RT-qPCR.

To assess induction of *Ifna4* expression in WT, IRF3^-/-^, or IFNAR1^-/-^ MEFs, 100,000 cells were seeded per well of a 12-well plate in the evening and stimulated the next day for 3 h with 100 U/mL of IFNβ or for 4 h by transfection of 5 µg/mL ISD with Lipofectamine 2000 as described above. Untreated and mock-transfected cells served as control, respectively. Samples were analysed by RT-qPCR.

For validation of the M35-mediated phenotype in M35-HAHA compared to EV iMEFs, 80,000 cells were seeded per well of a 12-well plate in the evening and stimulated the next day by transfection of 10 µg/mL of poly(I:C) with Lipofectamine 2000 as described above. Samples were harvested after 4 h and analysed by RT-qPCR.

The timepoints for analysis of transcripts regulated in response to (Alexa488-labelled) ISD transfection were determined in a small kinetic experiment. 450,000 EV iMEFs were seeded per well of a 6-well plate in the evening and stimulated the next morning by transfection with 5 µg/mL Alexa488-labelled ISD. Samples were harvested after 1, 2, 3, 4, 5, 6, or 8 h and analysed by RT-qPCR. Untreated cells and mock-transfected cells harvested after 4 h served as control.

### Infection of iBMDMs

To determine the effect of M35 on IRF3-dependent gene expression during MCMV infection, 800,000 iBMDMs were seeded in wells of a 6-well plate the day prior to the experiment and pre-treated by replacing conditioned medium with fresh medium containing 1 µM ruxolitinib. After 20 min, cells were infected on ice by replacing the cell culture supernatant with diluted MCMV REV or M35stop in fresh medium supplemented with 1 µM ruxolitinib to obtain an MOI of 0.1. Plates with cells were centrifuged at 805 g and 4°C for 30 min to enhance infection before shifting samples to 37°C with 5% CO_2_. The moment when the infected cells were shifted to 37°C incubation was defined as time point 0. After 30 min at 37°C, medium was replaced with fresh medium supplemented with 1 µM ruxolitinib. Samples were harvested after 4 h for analysis by RT-qPCR. To control the activity of ruxolitinib, an additional set of iBMDMs was pre-treated with DMSO or 1 µM ruxolitinib for 20 min and then treated either with 100 U/mL of murine IFNβ and ruxolitinib, or with ruxolitinib alone, or mock-treated with DMSO. Cells were harvested after 3 h and analysed by RT-qPCR.

### Quantitative PCR with reverse transcription (RT-qPCR)

For simultaneous analysis of multiple transcripts in primary MEF or iBMDM, cDNA was generated and then applied to SYBR Green-based qPCR. RNA was extracted using the innuPREP RNA mini Kit 2.0 (#845-KS-2040250, Analytik Jena, Jena, Germany), genomic DNA was removed using the iScript gDNA Clear cDNA Synthesis Kit (#1725035, Bio-Rad Laboratories, Feldkirchen, Germany), and cDNA was synthesised with the iScript cDNA Synthesis kit (#1708891, Bio-Rad Laboratories) according to the manufacturers’ instructions. Quantification of transcripts was performed using the GoTaq® qPCR Master Mix (#A6002, Promega) on a LightCycler 96 instrument (Roche). qPCR primers were as follows: *Rlp8* (Rlp8_for: 5’ CAACAGAGCCGTTGTTGGT-3’, Rlp8_rev: 5’ CAGCCTTTAAGATAGGCTTGTCA-3’); *Ifnb1* (IFNb_for: 5’ CTGGCTTCCATCATGAACAA-3’, IFNb_rev: 5’ AGAGGGCTGTGGTGGAGAA-3’), *Ifna4* (IFNa4_for: 5’-TCAAGCCATCCTTGTGCTAA-3’, IFNa4_rev: 5’-GTCTTTTGATGTGAAGAGGTTCAA-3’), *Isg15* (mISG15_fwd 5’-AGTCGACCCAGTCTCTGACTCT-3’, mISG15_rev 5’-CCCCAGCATCTTCACCTTTA-3’), *Ifit3*(mIfit3_fwd 5’-TGGACTGAGATTTCTGAACTGC-3’, mIfit3_rev 5’-AGAGATTCCCGGTTGACCTC-3’), *Rsad2* (mRsad2_fwd 5’-GGAAGGTTTTCCAGTGCCTCCT-3’, mRsad2_rev 5’-ACAGGACACCTCTTTGTGACGC-3’), *Stat1* (mStat1_for 5’-GCCTCTCATTGTCACCGAAGAAC-3’, mStat1_rev 5’-TGGCTGACGTTGGAGATCACCA-3’), *Nfkbia* (mNfkbia_for 5’-GCCAGGAATTGCTGAGGCACTT-3’, mNfkbia_rev 5’-GTCTGCGTCAAGACTGCTACAC-3’), *Tgfb1* (mTgfb1_for 5’-TGATACGCCTGAGTGGCTGTCT-3’, mTgfb1_rev 5’-CACAAGAGCAGTGAGCGCTGAA-3’).

To determine induction of *Ifna4* expression in primary MEF, RNA was prepared using the innuPREP RNA mini Kit 2.0, followed by removal of genomic DNA using the DNA-free kit (Ambion, Thermo Fisher Scientific). cDNA synthesis and quantification of transcripts was carried out using the EXPRESS One-Step Superscript™ RT-qPCR Kit (#11781200, Invitrogen, Thermo Fisher Scientific) with 100 ng RNA per sample on a LightCycler 96 instrument (Roche). PCR primers for *Rpl8* and *Ifna4* were used as given above together with the universal probe library probes (UPL, Roche) #5 and #3, respectively.

To determine the induction of *Ifnb1* expression in EV and M35-HAHA iMEFs, samples were lysed in RLT buffer supplemented with β-mercaptoethanol and RNA was purified using the RNeasy Mini Kit (#7410, Qiagen) with on-column DNase treatment (#79254, Qiagen) according to the manufacturer’s instructions. cDNA synthesis and quantification of transcripts was carried out using the EXPRESS One-Step Superscript™ RT-qPCR Kit as described above, using the *Rpl8* primer pair with probe #5 and the *Ifnb1* primer pair with probe #18.

Relative fold inductions were calculated using the 2^-ΔΔCt^ method.

### Statistical analysis

For luciferase reporter assays, RT-qPCR, and ChIP-qPCR, differences between two groups were evaluated by Student’s t-test (unpaired, two-tailed) using GraphPad Prism (version 5.0, GraphPad Software, San Diego, CA). p values < 0.05 were considered statistically significant. * p < 0.05, ** p < 0.01, *** p < 0.001, **** p < 0.0001.

Significance (p) of overlaps between two given groups of regulated gene products was determined by one-sided Fisher’s exact test, alternative=greater.

### SLAM-seq

To determine the amount of 4-thiouridine (4sU; #NT06186, Biosynth Carbosynth) for efficient labelling of nascent transcripts, 350,000 primary WT MEFs were seeded per well of a 6-well plate a day prior to the experiment. Cells were incubated with 100, 200, 400 or 800 µM of 4sU diluted into the conditioned medium and harvested after 2 h. Untreated cells served as control. Samples were harvested at indicated timepoints by lysis in 750 µL TRIzol® (#5596026, Invitrogen, Thermo Fisher Scientific) per well for 2 min. RNA of half of the sample volume was purified using the DirectZOL Microprep kit (#2060, Zymo Research) according to the manufacturer’s instructions including the on-column DNase digestion. For SLAM conversion, 90 µL of 20 mM iodoacetamide (Pierce™ IAA, #A39271, Thermo Fisher Scientific) solution in DMSO was mixed with 90 µL of RNA in 1x PBS and incubated at 50°C and 1,000 rpm for 30 min in the dark. The reaction was stopped by mixing with 20 µL of 1 M dithiothreitol (DTT). Converted RNA was purified using the RNA Clean & Concentrator-5 (#R1015, Zymo Research).

Quality and integrity of total RNA was controlled using a 2100 Bioanalyzer instrument with an RNA 6000 nano Chip (#5067-1511, Agilent Technologies, Santa Clara, CA, USA). The RNA sequencing library was generated from 300 ng total RNA using the TruSeq Stranded mRNA Library Prep kit (#20020595, Illumina, San Diego, CA, USA) with oligo-dT beads for capture of poly-A-mRNA according to manufacturer’s protocol. Quality and integrity of the libraries was controlled using a Bioanalyzer DNA 1000 Chip (#5067-1504, Agilent). The libraries were treated with Illumina Free Adapter Blocking Reagent (#20024145) and sequenced on an Illumina NextSeq 500 system using the NextSeq 500/550 Mid Output Kit v2.5 (#20024904, Illumina; 150 cycles, paired-end run 2x 75 bp) with an average of 1×10^7^ reads per RNA sample.

For characterisation of IRF3- and type I IFN-dependent genes, 300,000 primary WT, IRF3^-/-^ or IFNAR1^-/-^ MEFs were seeded per well of a 6-well plate a day prior to harvest to reach 80% confluency. To stimulate PRR signalling, ISD was mixed with an equal volume of Lipofectamine 2000 in 100µL Opti-MEM, incubated for 20 min and added to the conditioned medium to yield 5 µg/mL final concentration of ISD. For control, cells were mock-treated with the same amount of Lipofectamine 2000 diluted in Opti-MEM. Type I IFN signalling was stimulated in a parallel set of samples by pre-diluting murine IFNβ in Opti-MEM and adding the mix into the conditioned medium to reach a final concentration of 100 U/mL. Untreated samples served as control. All samples were prepared in quadruplicate. 2 h before lysis, 200 µM 4sU was added to the culture medium to label nascent RNAs. A set of untreated cells without 4sU treatment was prepared to control the incorporation rate. Samples were harvested at indicated timepoints by lysis in 750 µL TRIzol® and processed as described above.

Quality and integrity of total RNA was controlled on 5200 Fragment Analyzer System (Agilent Technologies, Santa Clara, CA, USA). The RNA sequencing library was generated from 200 ng total RNA using Dynabeads® mRNA DIRECT™ Micro Purification Kit (Thermo Fisher Scientific) for mRNA purification followed by NEBNext® Ultra™ II Directional RNA Library Prep Kit (New England BioLabs) according to manufacturer’s protocols. The libraries were treated with Illumina Free Adapter Blocking Reagent (Illumina, San Diego, CA, USA) and were sequenced on Illumina NovaSeq 6000 using NovaSeq 6000 S1 Reagent Kit (150 cycles, paired end run 2x 150 bp) with an average of 3×10^7^ reads per RNA sample.

Samples subjected to total transcriptome analysis were generated and processed similar to SLAM-seq samples with minor changes: 450,000 EV or M35-HAHA iMEFs were seeded per sample, stimulation was conducted by transfection of 5 µg/mL of Alexa488-coupled ISD, and 200 µM 4sU was added for 90 min prior to lysis. RNA was purified using the DirectZOL Miniprep kit (#R2050, Zymo Research, Freiburg, Germany), and 2 µg RNA per sample was used for SLAM conversion. Converted RNA was purified using the RNeasy Micro Kit (#74004, Qiagen) and measured using a Qubit™ Fluorometer (Thermo Fisher Scientific) with the Qubit RNA HS Assay Kit (#Q32852, Thermo Fisher Scientific). The RNA sequencing library was generated from 100 ng RNA using the NEBNext® Ultra II Directional RNA Library Prep Kit for Illumina® (#57760S and #57765S, New England Biolabs, Frankfurt am Main, Germany) with the NEBNext® Poly(A) mRNA Magnetic Isolation Module (#E7490, New England Biolabs) and SPRIselect beads (#B23319, Beckman Coulter) according to the manufacturer’s protocols. Quality and integrity of the libraries was controlled on a 2100 Bioanalyzer Instrument (Agilent) using a DNA chip (#5067, Agilent). The libraries were treated with Illumina Free Adapter Blocking Reagent (#20024145) and sequenced on an Illumina NovaSeq 6000 system using the NovaSeq 6000 S4 Individual Lane Loading Reagent kit (#20028313, Illumina; 150 cycles, paired end run 1x 111 bp) with an average of 2×10^7^ reads per sample. However, the conversion efficiency was too low to quantify newly synthesized transcripts, while overall integrity of transcripts was not influenced, therefore total transcripts were processed to evaluate this experiment.

### Data evaluation of SLAM-seq experiments

Reads from all three data sets (4sU titration, WT vs. IRF3^-/-^ vs. IFNAR1^-/-^ MEFs, and M35 vs. EV iMEFs) were processed by the same pipeline with the same parameters. First, reads were mapped against murine rRNA (less than 3% of reads in all cases) and common mycoplasma contaminations (less than 0.1% of reads in all cases) using bowtie2 version 2.3.0 (121) with standard parameters. All remaining reads were mapped to the murine genome (Ensembl version 90) using STAR version 2.5.3a (122) using parameters --outFilterMismatchNmax 20 -- outFilterScoreMinOverLread 0.4 --outFilterMatchNminOverLread 0.4 --alignEndsType Extend5pOfReads12 --outSAMattributes nM MD NH (uniquely mappable reads > 85% in all cases). Mapped reads from each of the three experiments were then further processed separately using GRAND-SLAM 2.0.7 (123) with parameters -trim5p 15 -modelall. The output tables of GRAND-SLAM were then further analysed using our grandR package 0.2.1 (124). Toxicity plots for the 4sU titration experiments were generated using the PlotToxicityTest function. Genes for the WT vs. IRF3^-/-^ vs. IFNAR1^-/-^ MEFs (E1) and M35 vs. EV iMEFs (E2) experiments were filtered such that at least 100 reads were present on average across replicates for at least one condition of E1 and one condition of E2. Differential gene expression was computed using the Wald test implemented in DESeq2 (125) with Benjamini-Hochberg multiple testing correction and the lfc package (126). Statistical tests and Spearman correlation were calculated with R. Venn diagrams were created using the R VennDiagram package. Heatmaps were created using the R pheatmap package and clustering was performed according to Euclidean distances with Ward’s clustering criterion.

### Functional enrichment analysis

The analysis for enriched biological processes based on Gene Ontology (GO) terms or for regulatory DNA motifs based on the transcription factor database TRANSFAC (127) was performed using the online tool g:GOSt of the g:Profiler web server (https://biit.cs.ut.ee/gprofiler/gost, version e107_eg54_p17_bf42210, accessed 25.01.2023 for GO biological processes; version e107_eg54_p17_bf42210, accessed 20.02.2023 for transcription factors; (128)) using own background data. P-values were corrected for multiple testing using the method by Benjamini and Hochberg for controlling the FDR. terms with FDR < 0.001 were considered statistically significant.

### Data availability

The SLAM-seq datasets have been deposited in the NCBI GEO database (Classification of IRF3- dependent vs IFNAR1-responsive gene induction in murine fibroblasts; Effect of MCMV M35 on global gene expression during PRR signalling in murine fibroblasts).

## Supplemental Material

Figures S1 – S9, Tables S1 – S5, and legends for all supplementary materials - Supplements.docx File 1 – File S1.pdf

## Acknowledgements

We thank Markus Stempel, Friedemann Weber, and Baca Chan for fruitful discussions and Sara Becker, Niels Lemmermann, and Kai Kropp for insightful comments. We also thank Daniela Gebauer, Maria Ebel, Christine Standfuß-Gabisch, Lennart Krause, and Susanne Zock-Emmenthal for excellent technical assistance. Special thanks go to the staff of the beamline P11 at the PETRAIII synchrotron (DESY campus Hamburg, Germany).

## Funding Information

This project was funded by the Deutsche Forschungsgemeinschaft (DFG, German Research Foundation) in the framework of the Research Unit FOR5200 DEEP-DV (443644894) projects BR 3432/7-1 (H.S., M.M.B), FR 2938/11-1 (C.C.F.), ER 927/4-1 (F.E.), DO 1275/12-1 (T.H., L.D.), GR 3318/5-1 (T.G., A.G.), LA 2941/18-1 (E.W., M.L.) and the Helmholtz Association (W2/W3-090) (V.M., M.M.B.). and supported by the Helmholtz Protein Sample Production Facility (PSPF).

The funders had no role in the design of the study; in the collection, analyses, or interpretation of data; in the writing of the manuscript, or in the decision to publish the results.

## Autor Contributions

Conceptualisation: H.S., V.M., S.S., E.W., F.E., W.B., M.M.B.

Data Curation: H.S., V.M., S.S., E.W., M.v.H., J.v.d.H., W.B., C.C.F., F.E., M.M.B.

Formal analysis: H.S., S.S., E.W., M.v.H., T.G., C.C.F., F.E.

Funding acquisition: L.J., W.B., M.L., A.G., L.D., C.C.F., F.E., M.M.B.

Investigation: H.S., V.M., S.S., E.W., T.H., T.G., K.B., M.v.H., J.v.d.H.

Methodology: H.S., V.M., S.S., E.W., T.G., K.B., J.v.d.H., L.J., W.B., F.E., M.M.B.

Project administration: M.M.B.

Resources: L.J., J.v.d.H., M.L., L.D., A.G., W.B., C.C.F., F.E., M.M.B.

Software: S.S., F.E., C.C.F., T.G.

Supervision: H.S., M.M.B.

Validation: H.S., V.M., F.E.

Visualisation: H.S., S.S., C.C.F., F.E.

Writing – original draft: H.S., V.M., S.S., J.v.d.H., C.C.F., F.E.

Writing – reviewing and editing: H.S., V.M., S.S., E.W., T.H., T.G., A.G., L.D., M.L., M.v.H., L.J., K.B., W.B., C.C.F., F.E., M.M.B.

## Notes

### Competing Interest Statement

The authors have declared no competing interest.

