## Supplementary Figures and Tables for "The Cytomegalovirus M35 Protein Modulates Transcription of *Ifnb1* and Other IRF3-Driven Genes by Direct Promoter Binding"

### Supplementary Material

#### Supplementary Figures

Figure S1

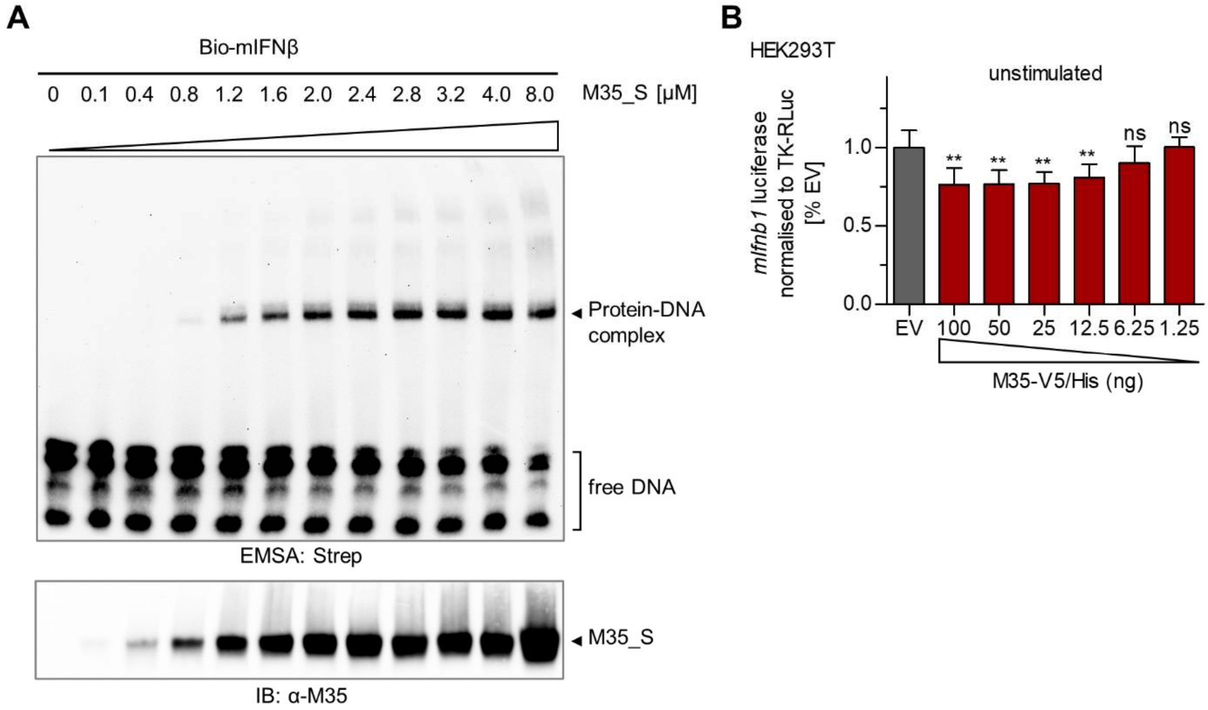

**Figure S1:** Titration of purified M35_S in EMSA and of ectopically expressed M35-V5/His in the luciferase reporter assay without stimulation.

(A) Representative experiment of the determination of the M35_S-DNA binding affinity by EMSA. EMSA was performed as before (Figure 1E) with increasing amounts (0.1 to 8 µM) of purified M35_S protein and a 5'-biotinylated double-stranded 56 bp oligonucleotide probe containing the sequence of the murine IFNβ enhancer (Bio-mIFNβ). One representative of three independent experiments is shown.

(B) Luciferase reporter assay for M35-mediated modulation of *Ifnb1* expression in the absence of stimulation. Luciferase reporter assays were performed as described before (Figure 1D) by transfection of HEK293T cells with *mIfnb1*-FLuc, and EV or a titration series of M35-V5/His, filled up with EV to 100 ng. Firefly luciferase values were normalized to *Renilla* luciferase values and results normalized to EV control. Data are represented as mean ±SD combined from three independent experiments. Significance compared to EV was calculated by Student's *t*-test (unpaired, two-tailed) comparing M35 derivatives to EV, ns not significant, ** $p < 0.01$.

Figure S2

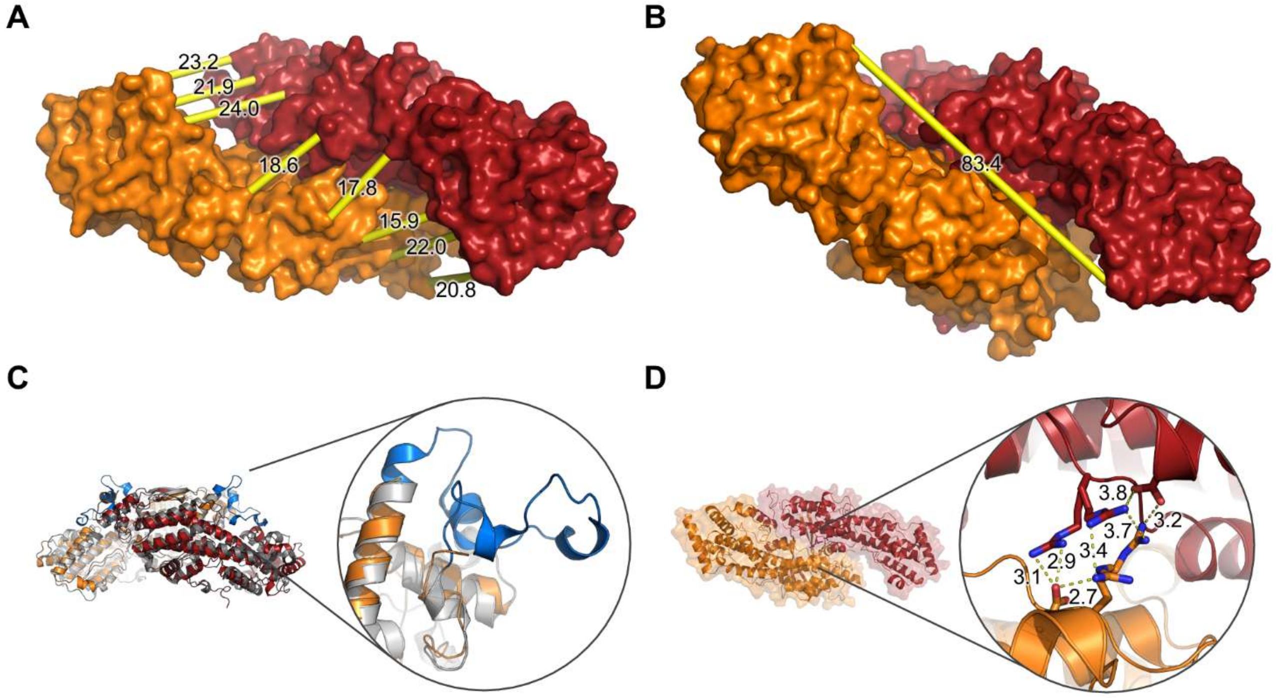

**Figure S2:** Dimensions of the groove at the dimer interface, inspection of the unresolved loop, and orientation of the residue R69 in the M35 crystal structure.

(A, B) Measurements approximating (A) the diameter and (B) the length of the groove at the dimer interface of M35 (surface representation, monomers indicated in orange and red). Yellow tubes indicate the measured distances, marked in Å. (C) Superposition of each one chain of MCMV M35 (orange) with HHV6B U14 (grey) depicted as ribbons with close-up of the region that is unresolved in M35 and forms a loop in U14 (blue). (D) Ribbon representation of the M35 crystal structure with close-up on the residues R69 that adapts two conformations (represented as sticks, coloured by element). Potential interactions distances of R69 with the R69 or the D44 residues of the other M35 moiety at the dimer interface are indicated by dashed yellow lines with measurements given in Å.

Figure S3

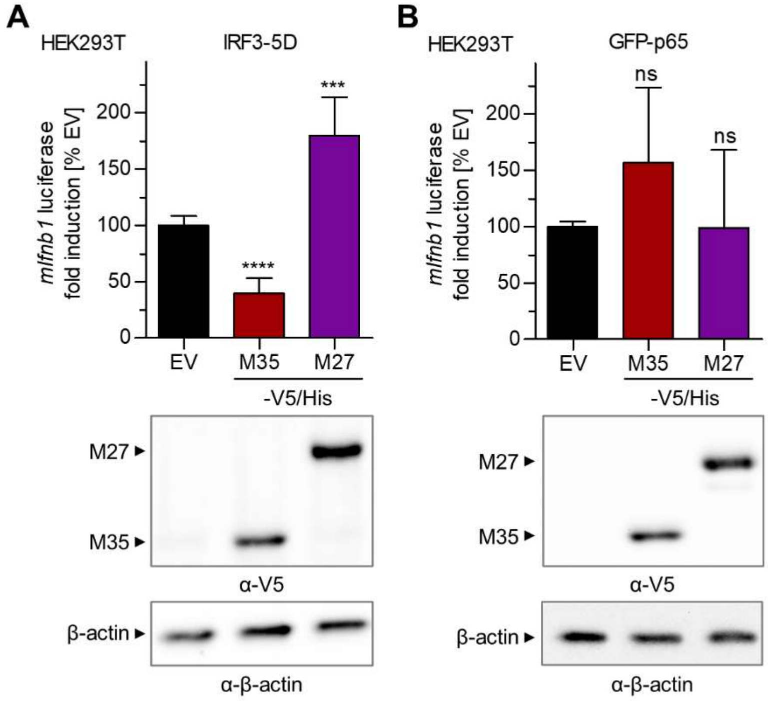

**Figure S3:** M35 inhibits induction of the murine *Ifnb1* promoter mediated by IRF3-5D, but not by NF-kB subunit p65.

Analysis of M35-mediated inhibition of *Ifnb1* transcription in the luciferase reporter assay stimulated by overexpression of the transcription factor (A) IRF3-5D or (B) GFP-p65. (A-B) Luciferase reporter assays were performed as described before (Figure 1D) by transfection of HEK293T cells with expression plasmids for (A) constitutively active IRF3-5D or (B) GFP-p56 (stimulated conditions) or the respective EV (unstimulated conditions), *mIfnb1*-FLuc, and indicated expression plasmids for M35-V5/His, M27-V5/His or corresponding EV. Values were normalized to EV control. Data was combined from three independent experiments, represented as mean ±SD, and significance compared to EV was calculated by Student's *t*-test (unpaired, two-tailed), ns not significant, *** $p < 0.001$, **** $p < 0.0001$. Lysates were analysed by immunoblotting with a V5-specific antibody. Detection of β-actin served as loading control.

Figure S4

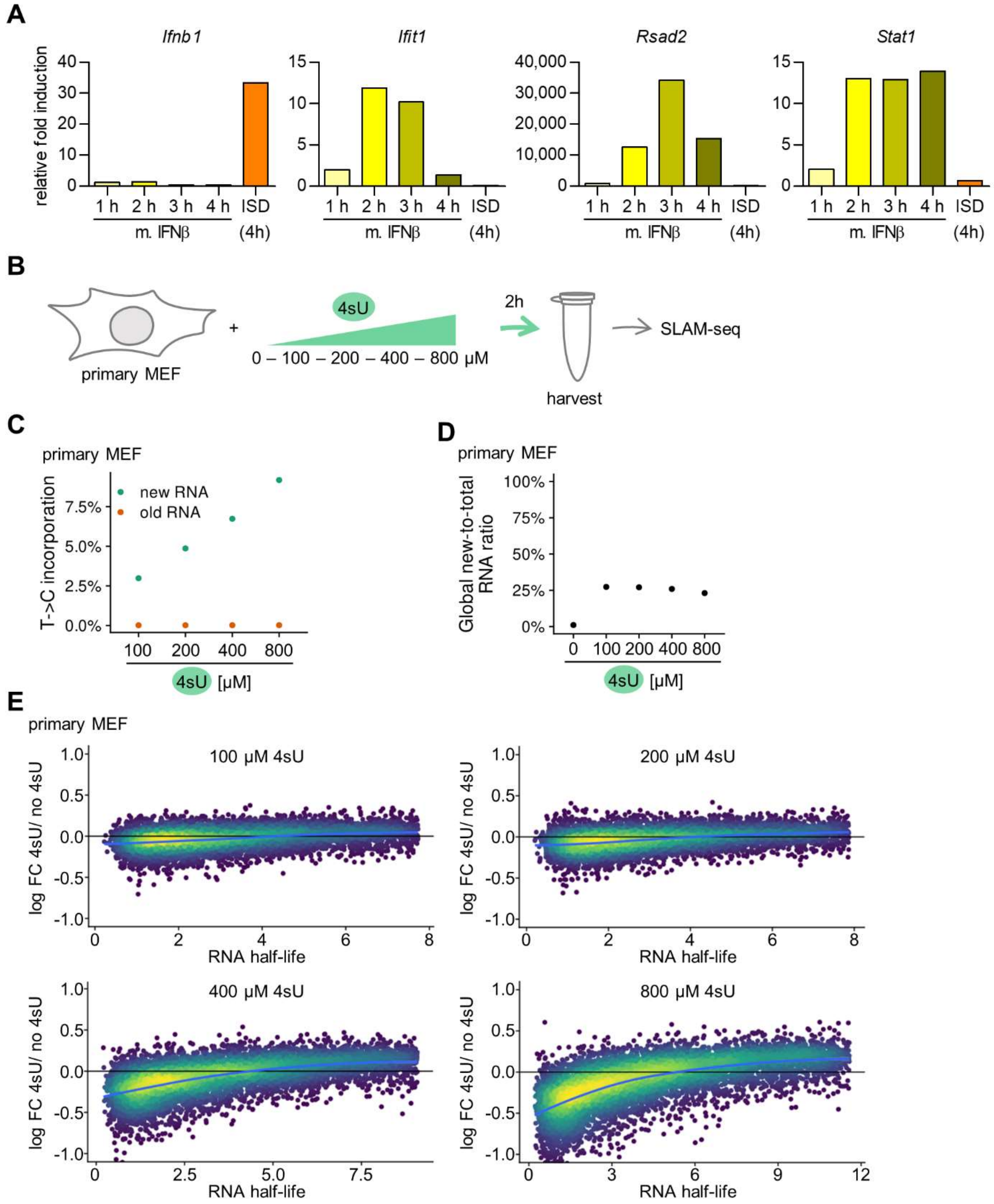

**Figure S4:** Kinetics of ISG induction and titration of 4-thiouridine for metabolic labelling of RNA in primary MEFs.

(A) Primary MEFs were left stimulated with 100 U/mL of murine IFNβ for indicated time points or with 5 µg/mL of ISD for 4 h or mock treated. Relative expression of *Ifnb1*, *Ifit1*, *Rsad2,* or *Stat1* normalised to *Rpl8* of treated compared to mock-treated samples was quantified by RT-qPCR. One of two independent experiments is presented. (B) Titration of 4-thiouridine (4sU) for labelling of newly synthesized RNA. Primary MEFs were incubated with 100 to 800 µM of 4-thiouridine (4sU) for 2 h or left untreated and the incorporation rate was determined by SLAM-seq. (C-D) The amount of 4sU is plotted against (C) the T->C conversion rate (incorporation frequency) in transcripts and against (D) the ratio of new RNA in indicated samples. (E) 4sU dropout plot (1). RNA half-life (h) is plotted against $log_2$FC of 4sU-treated versus untreated samples of indicated 4sU amounts applied to primary MEFs. A local polynomial regression curve is shown in blue. Excessive 4sU labelling (400uM and 800uM) resulted in apparent downregulation of genes with short RNA half-lives which could be indicative of a partial shutdown of transcription or technical issues during library preparation, sequencing or read mapping of 4sU-containing RNA.

Figure S5

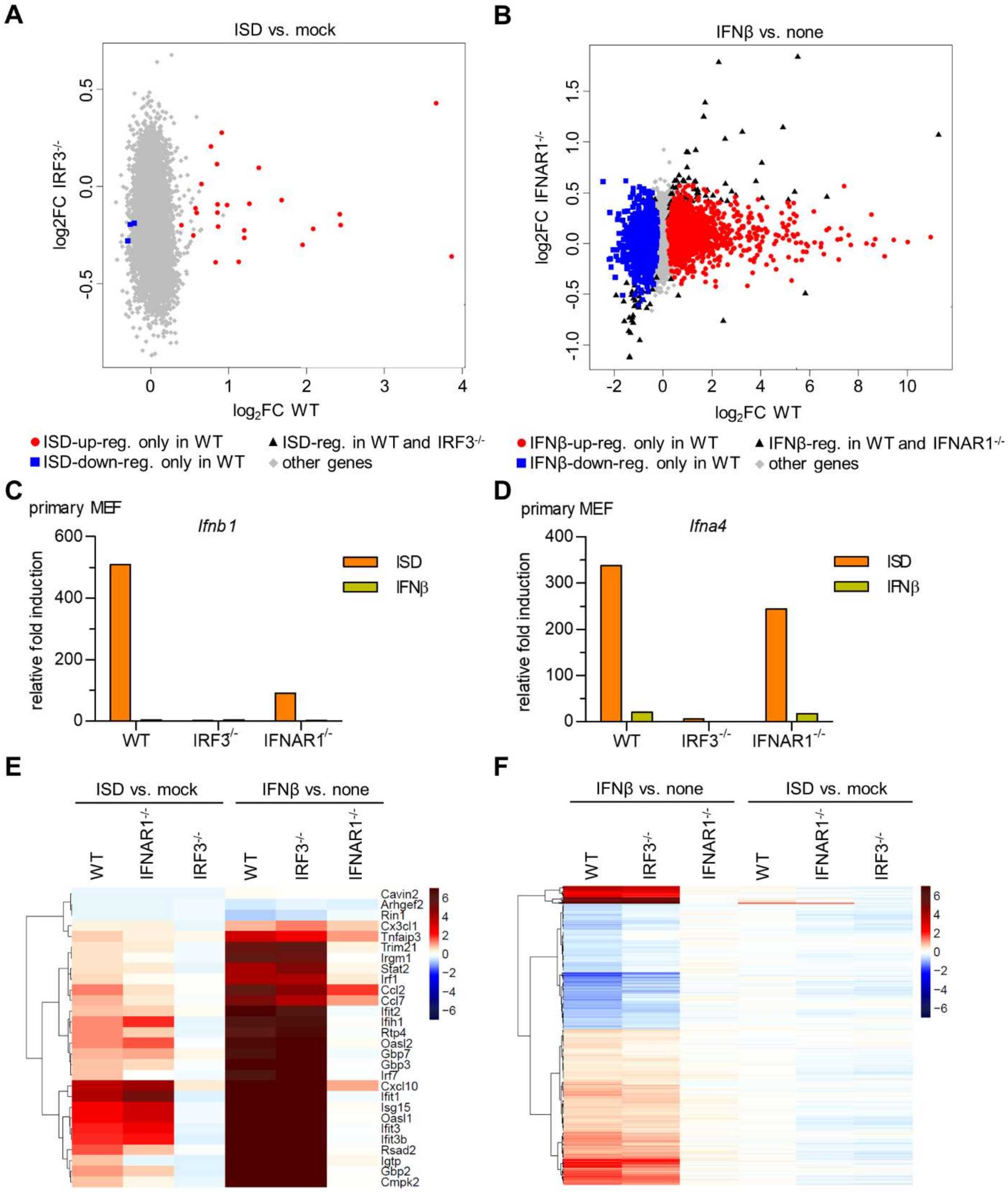

**Figure S5:** Determination and characterisation of IRF3-dependent and IFNAR1-responsive genes in primary MEFs.

(A-B) Scatter plots for $\log_2$FC of newly synthesized transcripts (A) in response to ISD treatment vs. mock-transfection in WT (x axis) compared to IRF3$^{-/-}$ MEFs (y axis), or (B) in response to IFNβ treatment vs. without treatment (none) in WT (x axis) compared to IFNAR1$^{-/-}$ MEFs (y axis). Transcripts significantly (FDR ≤ 0.01) up-regulated only in WT are highlighted red ($\log_2$FC > 0), transcripts significantly down-regulated only in WT are highlighted in blue ($\log_2$FC < 0). Transcripts responding to

(A) ISD transfection independently of IRF3 (FDR > 0.01 in IRF3$^{-/-}$) or (B) to IFNβ stimulation independently of IFNAR1 (FDR > 0.01 in IFNAR1$^{-/-}$) are marked as black triangles. (C, D) Primary WT, IRF3$^{-/-}$ or IFNAR1$^{-/-}$ MEFs were stimulated as in (A-B) and analysed by RT-qPCR for induction of (C) *Ifnb1* or (D) *Ifna4*, normalised to *Rpl8*. One of two independent experiments is presented. (E-F) Heatmaps showing $log_2$FC (blue: down-, red: up-regulation) for indicated conditions and cells for (E) IRF3-dependent genes, or for (F) IFNAR1-responsive genes in WT, IRF3$^{-/-}$ or IFNAR1$^{-/-}$ MEFs.

Figure S6

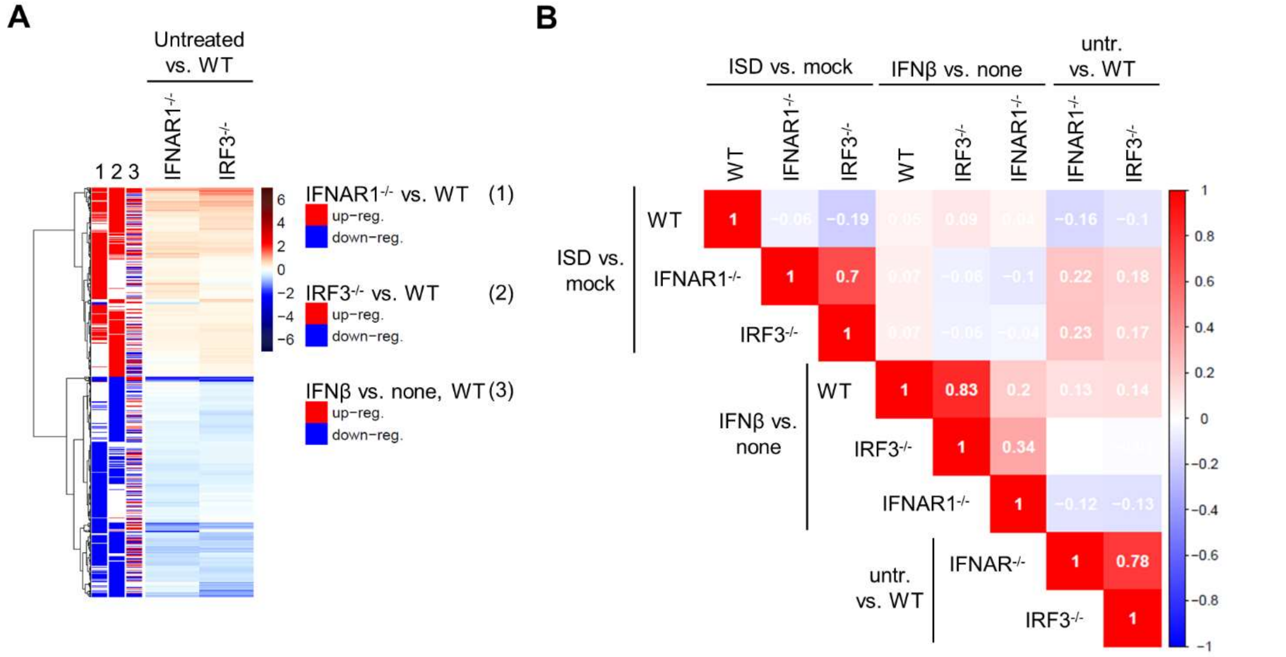

**Figure S6:** Knockout of key components of the type I IFN signalling pathway affects basal gene expression.

(A) Heatmap showing $log_2$FC in IRF3-/- or IFNAR1-/- compared to WT MEFs for all genes that are significantly regulated (FDR ≤ 0.01) in at least one of the knockouts. Genes significantly up- (red) or down-regulated (blue) in untreated (1) IFNAR1-/- or (2) IRF3-/- compared to WT MEFs or (3) by IFNβ stimulation in WT MEFs are marked at the left. (B) Correlation plot showing spearman correlations for pairwise comparisons of $log_2$FC over the whole transcriptome between WT, IRF3-/- or IFNAR1-/- MEFs with ISD, IFNβ or no stimulation (untr.).

Figure S7

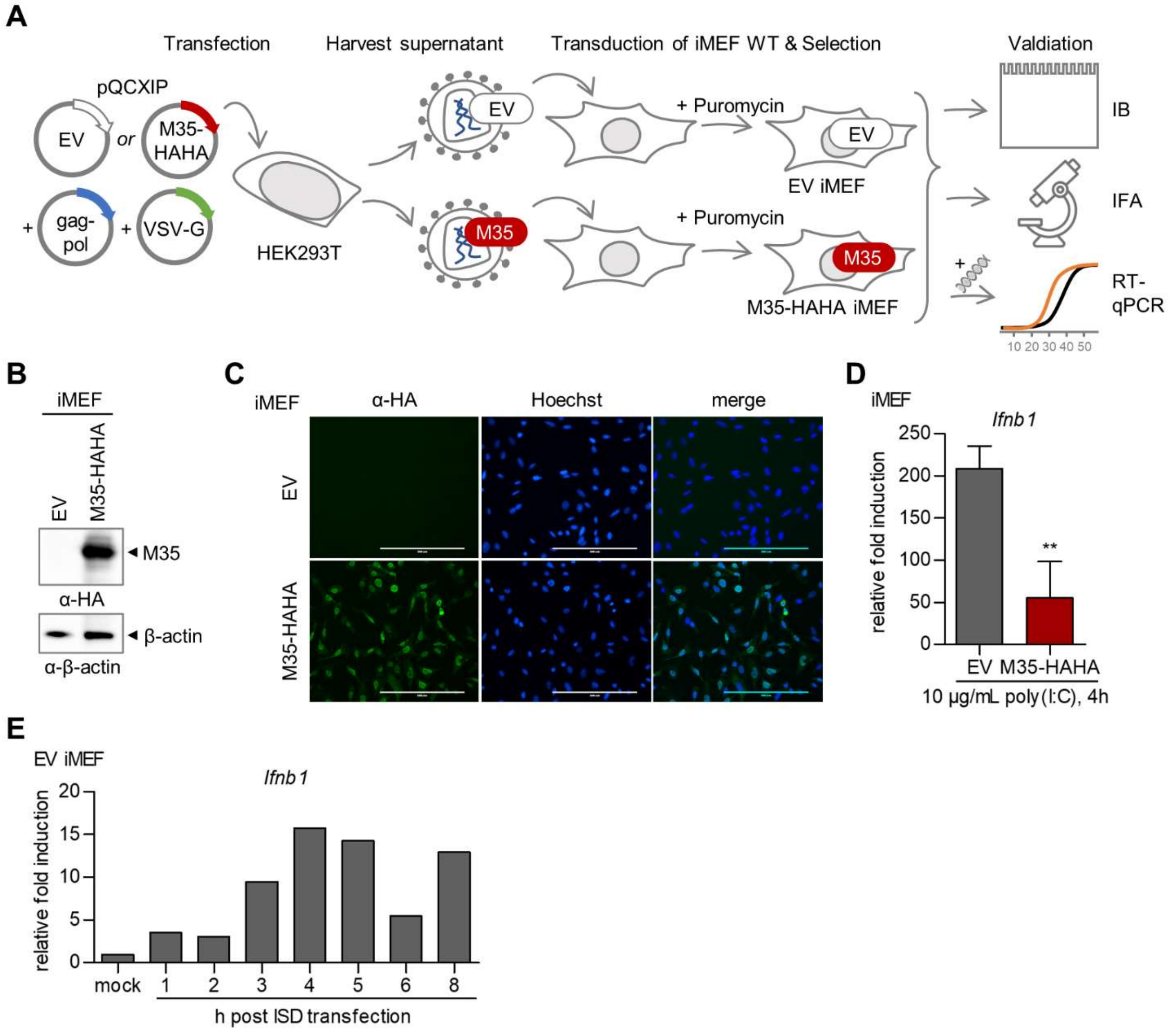

**Figure S7:** Generation and characterisation of stably M35-HAHA expressing immortalised MEFs.
(A) Generation and characterisation of stable cell lines of immortalised MEFs (iMEFs) with constitutive expression of M35-HAHA. HEK293T cells were transfected with retroviral transduction and packaging plasmids, and generated retroviral particles were used to infect WT iMEFs. Cell pools transduced with pQCXIP plasmids were selected by the encoded antibiotic resistance for puromycin, yielding EV and M35-HAHA iMEFs, respectively. Expression of M35-HAHA was validated by immunoblot (IB) of cell lysates, immunofluorescence assay (IFA) on cell pools, and analysis of *Ifnb1* expression upon PRR stimulation by RT-qPCR. (B) Immunoblot of EV and M35-HAHA iMEF lysates with an HA-specific antibody. Detection of β-actin served as loading control. (C) Immunofluorescence assay of EV and M35-HAHA iMEFs with an HA-specific antibody. Nuclei were stained with Hoechst. The scale bar represents 200 µm. (D) EV and M35-HAHA iMEFs were transfected with 10 µg/mL of poly(I:C) or mock-transfected, harvested after 4 h and analysed by RT-qPCR for induction of *Ifnb1*, normalised to *Rpl8*. Shown are combined results of three independent experiments. Significance compared to induction in EV iMEFs was calculated by Student's *t*-test (unpaired, two-tailed), ** $p < 0.01$. (E) EV iMEFs transfected with 5 µg/mL of Alexa488-labelled ISD or mock-transfected (4 h) were lysed at indicated time points and analysed by RT-qPCR for *Ifnb1* expression, normalised to *Rpl8*. Shown is one of two independent experiments.

Figure S8

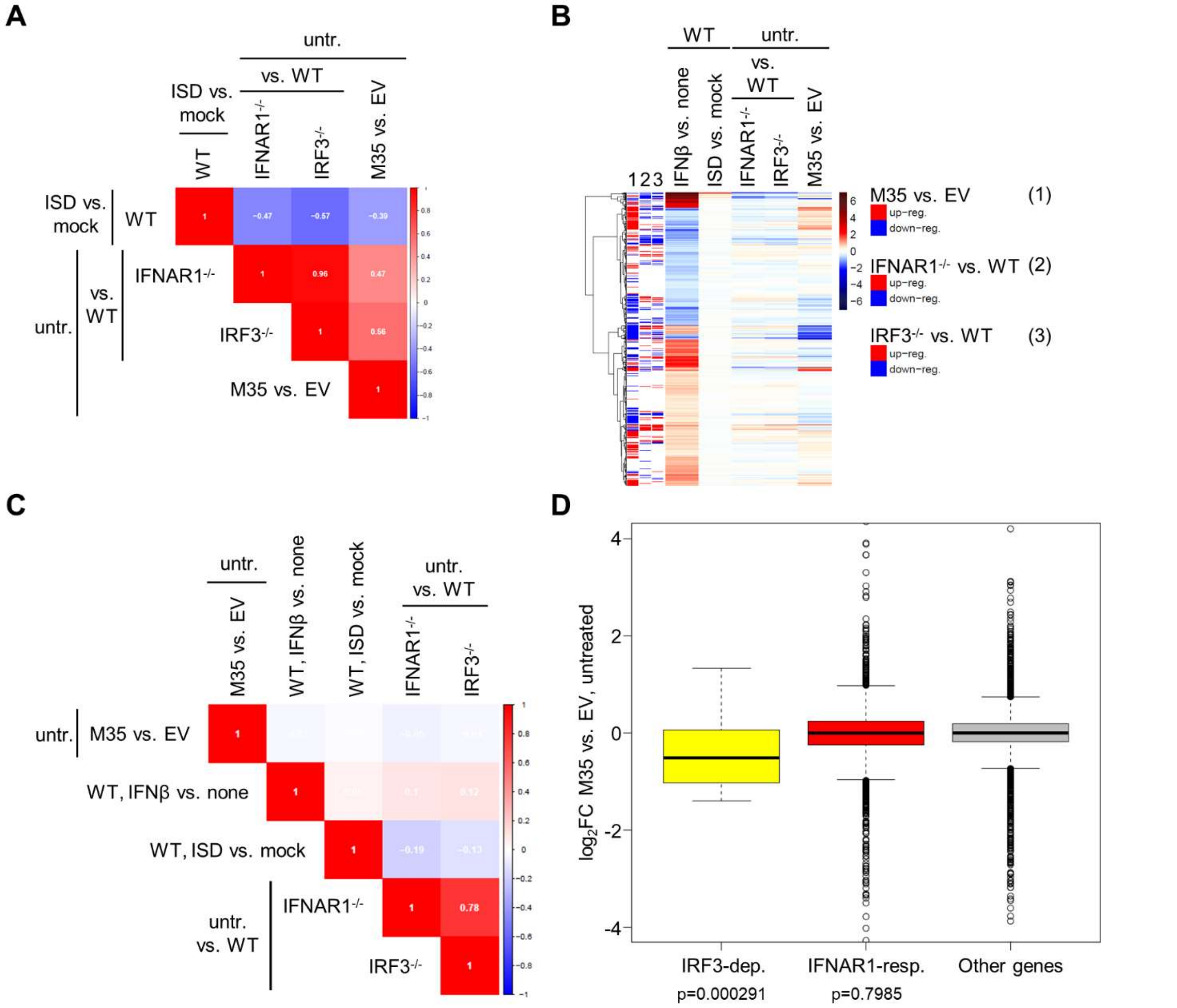

**Figure S8:** Effect of M35 on IRF3- or type I IFN signalling-dependent transcripts.

(A) Correlation plot showing spearman correlations (blue: negative, red: positive correlation) for $log_2FC$ of IRF3-dependent genes determined by SLAM-seq between ISD-stimulated vs. mock-treated WT MEFs, untreated (untr.) IFNAR1$^{-/-}$ or IRF3$^{-/-}$ vs. WT MEFs, and untreated M35-HAHA vs. EV iMEFs. (B) Heatmap showing the $log_2FC$ (blue: down-, red: up-regulation) in indicated SLAM-seq samples for the 2,888 genes IFNAR1-responsive genes. Genes significantly (FDR ≤ 0.01) up- (red) or down-regulated (blue) in (1) M35-HAHA compared to EV iMEFs or in (2) IFNAR1$^{-/-}$ or (3) IRF3$^{-/-}$ compared to WT MEFs are marked at the left. (C) Correlation plot showing spearman correlation (blue: negative, red: positive correlation) comparing the $log_2FC$ of IFNAR1-responsive genes between untreated M35-HAHA vs. EV iMEFs, IFNβ-stimulated vs. untreated WT MEFs, ISD-stimulated vs. mock-treated WT MEFs, and untreated IFNAR1$^{-/-}$ or IRF3$^{-/-}$ vs. WT MEFs. (D) Boxplots showing the distribution of $log_2FC$ in M35-HAHA compared to EV iMEFs for IRF3-dependent ISD-responsive genes (IRF3-dep.), IFNAR1-dependent IFNβ-responsive genes (IFNAR1-resp.) and other differentially expressed genes (other genes). P-values comparing median $log_2FC$ for IRF3-dependent and IFNAR1-responsive genes, respectively, to other genes were calculated by Wilcoxon rank sum test and are indicated below.

Figure S9

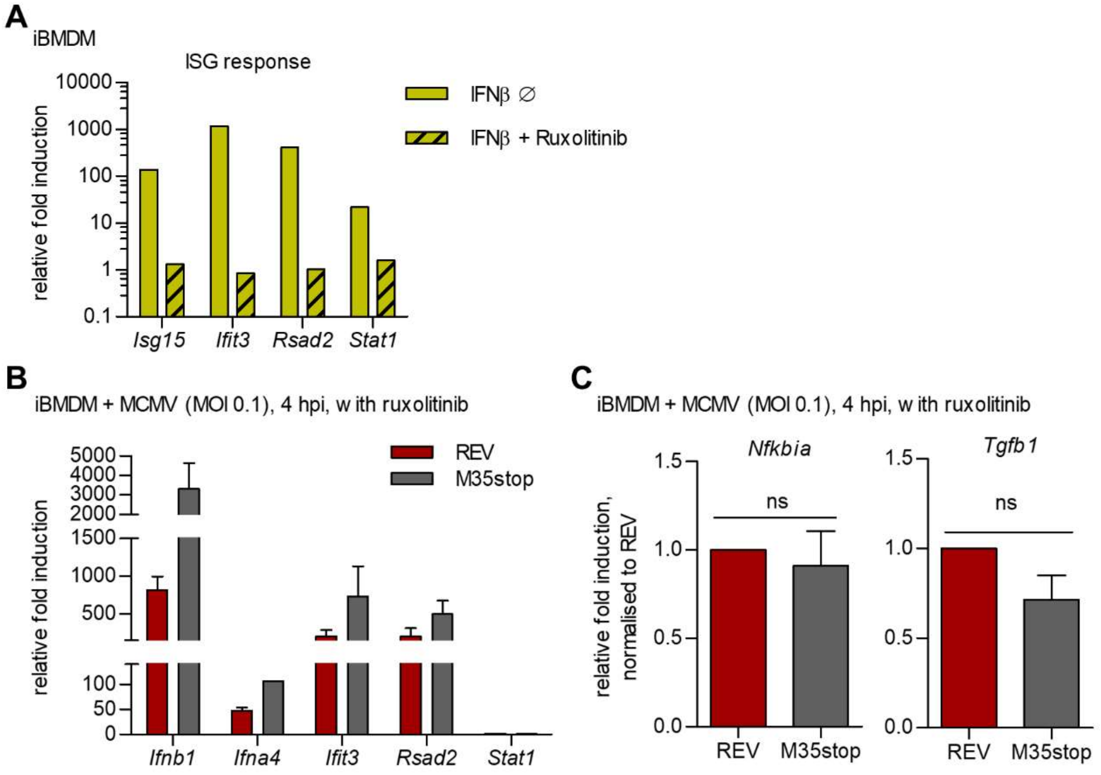

**Figure S9:** Characterisation of the response of IRF3-dependent genes to MCMV stimulation under inhibition of type I IFN signalling.

(A) WT iBMDMs were treated with 1 µM of ruxolitinib (IFNAR signalling inhibitor) for 20 min prior and during the experiment, or treated with DMSO vehicle. Cells were left unstimulated or stimulated with 100 U/mL of murine IFNβ for 3 h. Samples were harvested and analysed by RT-qPCR for induction of the ISGs *Isg15*, *Ifit3*, *Rsad2*, and *Stat2*, normalised to *Rpl8*. Shown is one experiment that was performed in parallel with the following iBMDM infection experiment. (B-C) Response of (B) IRF3-dependent or (C) proinflammatory genes upon infection with MCMV with or without M35. iBMDMs pre-treated with 1 µM of ruxolitinib were infected with MCMV M35stop-Revertant (REV) or MCMV M35stop (M35stop) at MOI of 0.1 or mock infected. Cells were harvested 4 h post infection for analysis by RT-qPCR. (B) Presented are the results shown in Figure 9F for *Ifnb1*, *Ifna4*, *Ifit3*, and *Rsad2* as relative fold induction normalised to *Rpl8* without normalisation to REV infection. Data is shown as mean ±SD and combined from two (*Ifna4*) or three (*Ifnb1*, *Ifit3*, *Rsad2*) independent experiments. (C) Relative fold induction of *Nfkbia* and *Tgfb1* transcripts was calculated based on the housekeeping gene *Rpl8* and normalised to REV infection. Data is shown as mean ±SD and combined from two independent experiments.

#### Supplementary Tables

**Table S1:** Determination of the DNA-binding affinity based on signals of EMSA probes. The measurements correspond to the blot in Figure S1A.

| Purified M35_S protein [µM] | Relative signal intensity[a] of protein-DNA complex | Relative signal intensity[a] of free DNA | fraction bound[b] |
|---|---|---|---|
| 8 | 13243.31 | 9812.321 | 0.574 |
| 4 | 18517.67 | 19079.77 | 0.493 |
| 3.2 | 15982.38 | 17487.89 | 0.478 |
| 2.8 | 16560.14 | 20767.08 | 0.444 |
| 2.4 | 15250.72 | 22218.45 | 0.407 |
| 2 | 12915.6 | 25079.05 | 0.340 |
| 1.6 | 8978.995 | 28992.37 | 0.236 |
| 1.2 | 5861.681 | 34421.99 | 0.146 |
| 0.8 | 468.849 | 39387.04 | 0.012 |
| 0.4 | 282.121 | 37288.22 | 0.008 |
| 0.1 | 0 | 36988.32 | 0.000 |
| 0 | 0 | 34260.61 | 0.000 |

[a] Intensities of probe signals on EMSA blots were quantified using Fiji.

[b] The bound fraction was calculated by dividing the signal of the bound probes by the total signal per lane.

**Table S2:** Data collection and refinement statistics for the crystal structure of M35_S.

| | M35 |
|---|---|
| **Data collection** | |
| Space group | P $2_1$ |
| Cell dimensions | |
| *a*, *b*, *c* (Å) | 59.1, 130.7, 67.8 |
| a, b, g (°) | 90.0, 101.3, 90.0 |
| Resolution (Å) | 66.52-1.94 (2.15-1.94) |
| Ellipsoidal[a] resolution (Å) (direction*) [b] | 2.46 (a*)<br>1.92 (b*)<br>2.12 (c*) |
| $R_{\mathrm{merge}}$ | 0.148 (2.151) |
| $R_{\mathrm{pim}}$ | 0.047 (0.488) |
| CC(1/2) | 0.999 (0.581) |
| *I* / s*I* | 11.6 (1.6) |
| Completeness | |
| spherical (%) | 67.9 (13.1) |
| ellipsoidal (%) | 92.0 (58.9) |
| Redundancy | 10.6 (8.4) |
| **Refinement** | |
| Resolution (Å) | 57.95-1.94 (2.15-1.94) |
| No. reflections | 50226 (ellipsoidal) |
| $R_{\mathrm{work}}$ / $R_{\mathrm{free}}$ | 0.174/0.226 |
| No. atoms | |
| Protein | 6493 |
| Ligand/ion | 67 |
| Water | 299 |
| *B*-factors | |
| Protein | 36.7 |
| Ligand/ion | 57.6 |
| Water | 38.1 |
| R.m.s. deviations | |
| Bond lengths (Å) | 0.012 |
| Bond angles (°) | 1.24 |
| PDB ID | 8BTJ |

*Values in parentheses are for highest-resolution shell.

[a] Data were processed anisotropically via STARANISO included in the Autoproc suite.

[b] The resolution limits for three directions in reciprocal space (a*, b*, c*) are indicated here.

**Table S3:** Selection of proteins of the *Betaherpesvirinea* pp85 protein superfamily for multiple sequence alignment based on the ICTV master species list (version 2018b.v2).

| **Genus** | **Abbreviation** | **Species**[a] | **Strain** | **Genome accession** | **Host** | **Protein name** | **Protein ID** |
|---|---|---|---|---|---|---|---|
| *Cytomegalovirus* | CeHV-5 | Cercopithecine betaherpesvirus 5 | 2715 | NC_012783.2 | Meercat | tegument protein UL35 | YP_004936011.1 |
| *Cytomegalovirus* | HCMV | **Human betaherpesvirus 5** | TB40/E | EF999921.1 | Human | UL35 | ABV71565.1 |
| *Cytomegalovirus* | CCMV | Panine betaherpesvirus 2 (Chimpanzee cytomegalovirus) | Heberling | NC_003521.1 | Chimp | tegument protein UL35 | NP_612679.1 |
| *Cytomegalovirus* | SaHV-4 | Saimiriine betaherpesvirus 4 | SqSHV | NC_016448.1 | Squirrel monkey | tegument protein UL35 | YP_004940207.1 |
| *Muromegalovirus* | MCMV (MuHV-1) | **Murid betaherpesvirus 1** | Smith | GU305914.1 | Mouse | M35 | ADD10418.1 |
| *Muromegalovirus* | RCMV (MuHV-2) | Murid betaherpesvirus 2 | Maastrich | NC_002512.2 | Rat | pR35 | NP_064140.1 |
| *Muromegalovirus* | MuHV-8 (RCMV-E) | Murid betaherpesvirus 8 | English isolate | NC_019559.2 | Rat | E35 | YP_007016439.1 |
| *Proboscivirus* | EIHV-1 (EEHV) | **Elephantid betaherpesvirus 1** (Elephant endotheliotropic herpesvirus) | Raman | NC_020474.2 | Elephant | tegument protein UL35 | YP_007969827.1 |
| *Roseolovirus* | HHV6A | **Human betaherpesvirus 6A** | U1102 | NC_001664.4 | Human | U14 | NP_042906.1 |
| *Roseolovirus* | HHV6B | Human betaherpesvirus 6B | HST | AB021506.1 | Human | U14 | BAA78235.1 |
| *Roseolovirus* | HHV7 | Human betaherpesvirus 7 | JI | U43400.1 | Human | U14 | AAC54675.1 |
| *Quwivirus*[b] | CaHV-2 | Caviid betaherpesvirus 2 | 21222 | NC_020231.1 | Guinea pig | GP35 | YP_007417809.1 |

| Genus | Abbreviation | Species[a] | Strain | Genome accession | Host | Protein name | Protein ID |
|---|---|---|---|---|---|---|---|
| *Quwivirus*[b] | TuHV-1 | Tupaiid betaherpesvirus 1 | 2 | NC_002794.1 | Treeshrew | T35 | NP_116385.1 |
| | BHV | Bat Herpesvirus | B7D8 | JQ805139.1 | Bat | B35 | AFK83854.1 |

[a] Type species are highlighted in bold.

[b] The genus *Quwivirus* was assigned in 2021 (2).

**Table S4:** Transcription factors corresponding to the regulatory DNA motifs with significant enrichment in the promoters of genes significantly (FDR ≤ 0.01) up- or down-regulated in both WT and IFNAR1$^{-/-}$ MEFs after IFNβ treatment (IFNAR1-independent).

| **Term ID**[a] | **Term Name** | $\mathbf{p_{adjust}}$ |
|---|---|---|
| TF:M00052 | Factor: NF-kappaB; motif: GGGRATTTCC | $2.864 \times 10^{-4}$ |
| TF:M00774 | Factor: NF-kappaB; motif: NGGGANTTYCCMNNNN | $2.864 \times 10^{-4}$ |
| TF:M00053 | Factor: c-Rel; motif: SGGRNTTTCC | $2.864 \times 10^{-4}$ |
| TF:M10311 | Factor: NF-kappaB1; motif: GGGAAANYCCC | $2.904 \times 10^{-4}$ |
| TF:M10312 | Factor: NF-kappaB2; motif: GGGRAANYCCC | $3.395 \times 10^{-4}$ |
| TF:M03545 | Factor: c-Rel; motif: NGGGAATYTCCN | $7.049 \times 10^{-4}$ |
| TF:M03563_1 | Factor: RelA-p65; motif: GGGANTTTCCNN; match class: 1 | $7.049 \times 10^{-4}$ |
| TF:M00052_1 | Factor: NF-kappaB; motif: GGGRATTTCC; match class: 1 | $8.716 \times 10^{-4}$ |

[a] The functional enrichment analysis for regulatory DNA motifs based on the transcription factor database TRANSFAC was performed using the g:GOSt function of g:Profiler (version e107_eg54_p17_bf42210) applying FDR ($p_{adjust}$) < 0.001 as significance threshold (3).

**Table S5:** 20 most significantly enriched biological pathways among significantly (FDR ≤ 0.01) regulated genes in $IRF3^{-/-}$ and/or $IFNAR1^{-/-}$ compared to WT MEFs.

| **GO Term ID[a]** | **GO Term Name** | $\mathbf{p_{adjust}}$ |
|---|---|---|
| GO:0009605 | response to external stimulus | $1.168\times10^{-23}$ |
| GO:0051239 | regulation of multicellular organismal process | $5.817\times10^{-23}$ |
| GO:0032501 | multicellular organismal process | $5.817\times10^{-23}$ |
| GO:0050896 | response to stimulus | $1.116\times10^{-22}$ |
| GO:0007155 | cell adhesion | $3.046\times10^{-22}$ |
| GO:0007154 | cell communication | $1.224\times10^{-21}$ |
| GO:0035295 | tube development | $1.470\times10^{-21}$ |
| GO:0023052 | signaling | $3.069\times10^{-21}$ |
| GO:0035239 | tube morphogenesis | $6.663\times10^{-21}$ |
| GO:0009653 | anatomical structure morphogenesis | $7.068\times10^{-21}$ |
| GO:0042221 | response to chemical | $1.354\times10^{-19}$ |
| GO:0048731 | system development | $2.176\times10^{-19}$ |
| GO:0002376 | immune system process | $2.866\times10^{-19}$ |
| GO:0007166 | cell surface receptor signaling pathway | $2.919\times10^{-19}$ |
| GO:0009888 | tissue development | $2.919\times10^{-19}$ |
| GO:0048856 | anatomical structure development | $4.259\times10^{-19}$ |
| GO:0040011 | locomotion | $7.641\times10^{-19}$ |
| GO:0048513 | animal organ development | $1.139\times10^{-18}$ |
| GO:0007165 | signal transduction | $2.458\times10^{-18}$ |
| GO:0001944 | vasculature development | $5.607\times10^{-18}$ |

[a] The functional GO term enrichment analysis for biological processes was performed using the g:GOSt function of g:Profiler (version e107_eg54_p17_bf42210) applying FDR ($p_{adjust}$) < 0.001 as significance threshold (3).

#### Supplementary Files

**File S1:** Multiple sequence alignment of the pp85 protein superfamily of *Betaherpesvirinea.*
Protein sequences were selected as described in Figure 3A, aligned using Clustal Omega (4) and illustrated using Jalview (5). Amino acid characters are indicated by colour, and conservation of positions by shading (darker for higher conservation, threshold value 15). The grey occupancy histogram below reflects the number of aligned residues per position, and the consensus sequence is annotated with the percentage of different residues per position shown as a black histogram.
