## Supplementary File S1 for "The Cytomegalovirus M35 Protein Modulates Transcription of *Ifnb1* and Other IRF3-Driven Genes by Direct Promoter Binding"

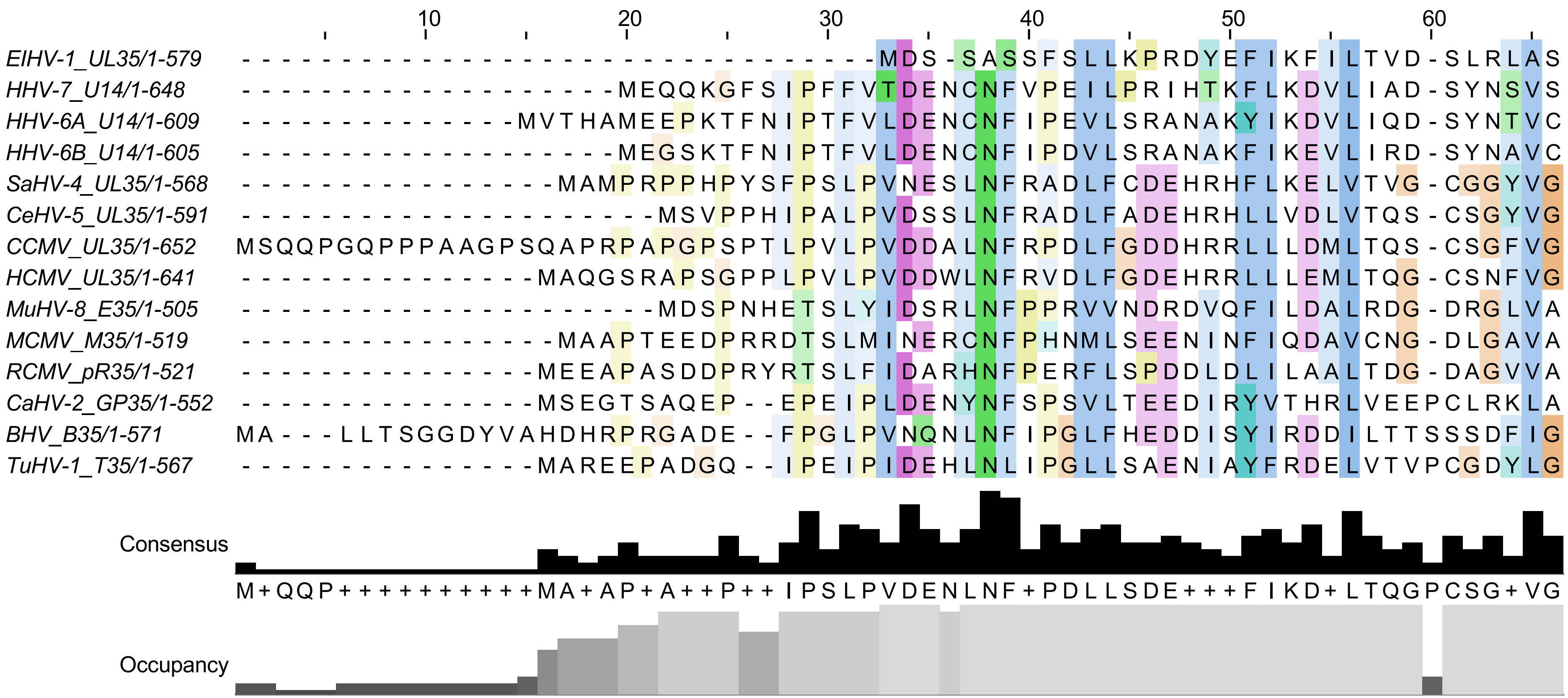
10
20
30
40
50
60
EIHV-1_UL35/1-579 --------------------------------MDS-SASSFSLLKPRDYEFIKFILTVD-SLRLAS
HHV-7_U14/1-648 -------------------MEQQKGFSIPFFVTDENCNFVPEILPRIHTKFLKDVLIAD-SYNSVS
HHV-6A_U14/1-609 --------------MVTHAMEEPKTFNIPTFVLDENCNFIPEVLSRANAKYIKDVLIQD-SYNTVC
HHV-6B_U14/1-605 -------------------MEGSKTFNIPTFVLDENCNFIPDVLSRANAKFIKEVLIRD-SYNAVC
SaHV-4_UL35/1-568 ----------------MAMPRPPHPYSFPSLPVNESLNFRADLFCDEHRHFLKELVTVG-CGGYVG
CeHV-5_UL35/1-591 ---------------------MSVPPHIPALPVDSSLNFRADLFADEHRHLLVDLVTQS-CSGYVG
CCMV_UL35/1-652 MSQQPGQPPPAAGPSQAPRPAPGPSPTLPVLPVDDALNFRPDLFGDDHRRLLLDMLTQS-CSGFVG
HCMV_UL35/1-641 ---------------MAQGSRAPSGPPLPVLPVDDWLNFRVDLFGDEHRRLLLEMLTQG-CSNFVG
MuHV-8_E35/1-505 ---------------------MDSPNHETSLYIDSRLNFPPRVVNDRDVQFILDALRDG-DRGLVA
MCMV_M35/1-519 ----------------MAAPTEEDPRRDTSLMINERCNFPHNMLSEENINFIQDAVCNG-DLGAVA
RCMV_pR35/1-521 ---------------MEEAPASDDPRYRTSLFIDARHNFPERFLSPDDLDLILAALTDG-DAGVVA
CaHV-2_GP35/1-552 ---------------MSEGTSAQEP--EPEIPLDENYNFSPSVLTEEDIRYVTHRLVEEPCLRKLA
BHV_B35/1-571 MA---LLTSGGDYVAHDHRPRGADE--FPGLPVNQNLNFIPGLFHEDDISYIRDDILTTSSSDFIG
TuHV-1_T35/1-567 ---------------MAREEPADGQ--IPEIPIDEHLNLIPGLLSAENIAYFRDELVTVPCGDYLG
Consensus
M+QQP++++++++++MA+AP+A++P++IPSLPVDENLNF+PDLLSDE+++FIKD+LTQGPCSG+VG
Occupancy

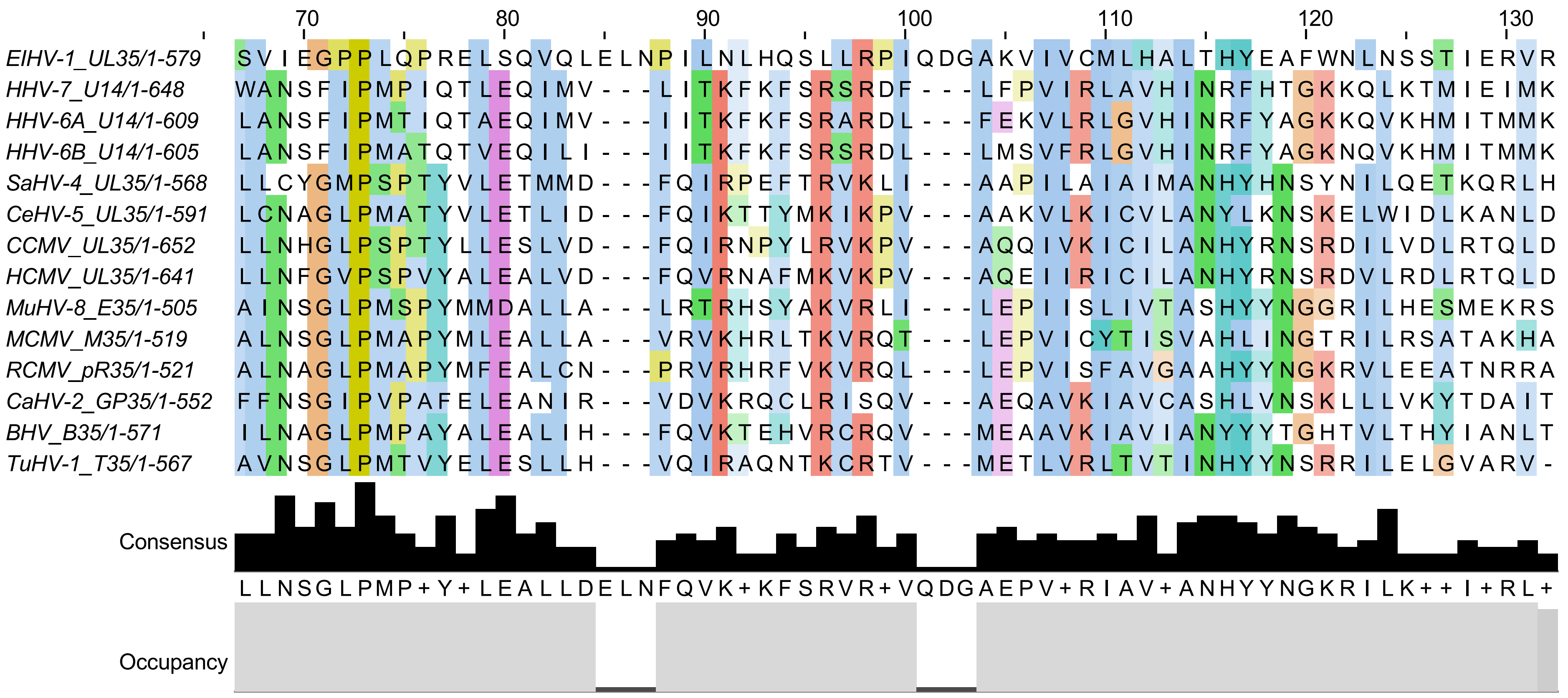
EIHV-1_UL35/1-579
HHV-7_U14/1-648
HHV-6A_U14/1-609
HHV-6B_U14/1-605
SaHV-4_UL35/1-568
CeHV-5_UL35/1-591
CCMV_UL35/1-652
HCMV_UL35/1-641
MuHV-8_E35/1-505
MCMV_M35/1-519
RCMV_pR35/1-521
CaHV-2_GP35/1-552
BHV_B35/1-571
TuHV-1_T35/1-567
Consensus
Occupancy
70
80
90
100
110
120
130

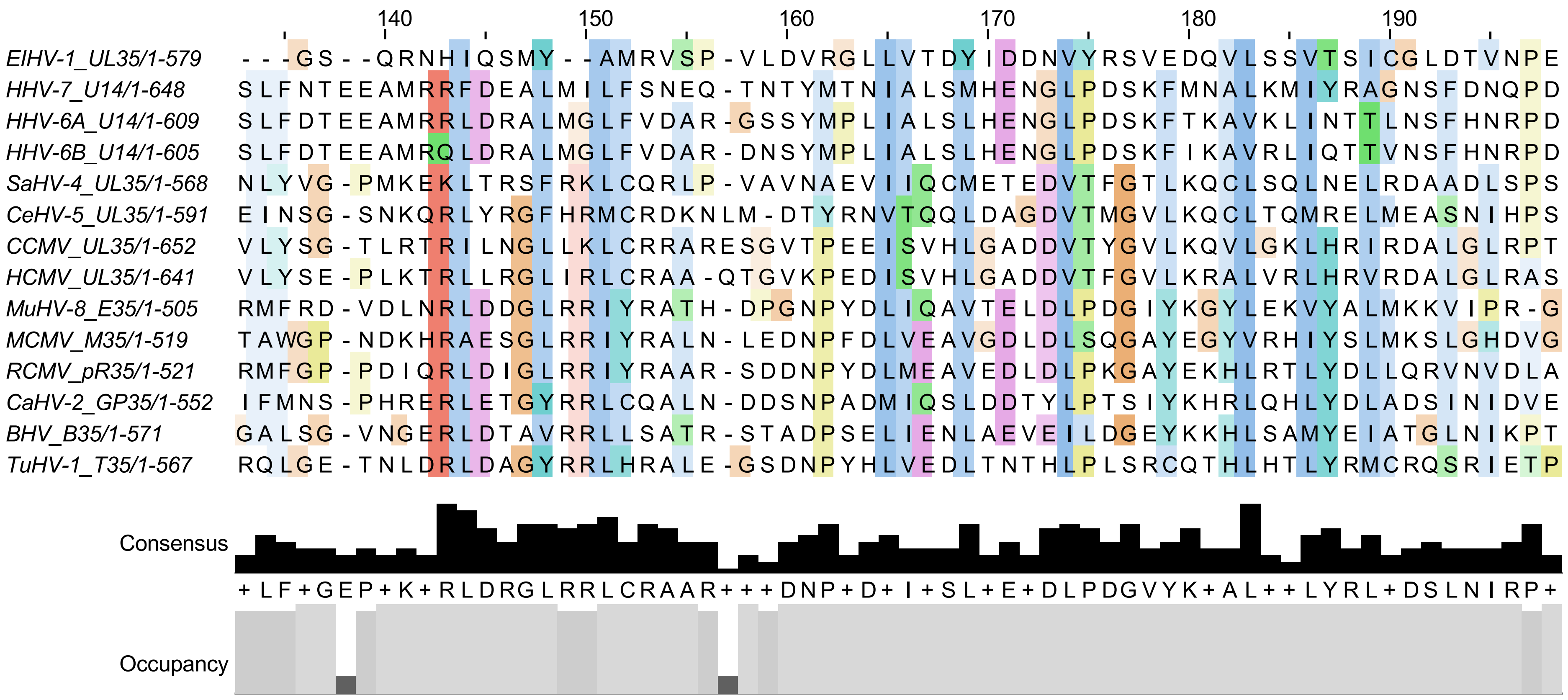

140
150
160
170
180
190
EIHV-1_UL35/1-579 ---GS--QRNHIQSMY--AMRVSP-VLDVRGLLVTDYIDDNVYRSVEDQVLSSVTSICGLDTVNPE
HHV-7_U14/1-648 SLFNTEEAMRRFDEALMILFSNEQ-TNTYMTNIALSMHENGLPDSKFMNALKMIYRAGNSFDNQPD
HHV-6A_U14/1-609 SLFDTEEAMRRLDRALMGLFVDAR-GSSYMPLIALSLHENGLPDSKFTKAVKLINTTLNSFHNRPD
HHV-6B_U14/1-605 SLFDTEEAMRQLDRALMGLFVDAR-DNSYMPLIALSLHENGLPDSKFIKAVRLIQTTVNSFHNRPD
SaHV-4_UL35/1-568 NLYVG-PMKEKLTRSFRKLCQRLP-VAVNAEVIIQCMETEDVTFGTLKQCLSQLNELRDAADLSPS
CeHV-5_UL35/1-591 EINSG-SNKQRLYRGFHRMCRDKNLM-DTYRNVTQQLDAGDVTMGVLKQCLTQMRELMEASNIHPS
CCMV_UL35/1-652 VLYSG-TLRTRILNGLLKLCRRARESGVTPEEISVHLGADDVTYGVLKQVLGKLHRIRDALGLRPT
HCMV_UL35/1-641 VLYSE-PLKTRLLRGLIRLCRAA-QTGVKPEDISVHLGADDVTFGVLKRALVRLHRVRDALGLRAS
MuHV-8_E35/1-505 RMFRD-VDLNRLDDGLRRIYRATH-DPGNPYDLIQAVTELDLPDGIYKGYLEKVYALMKKVIPR-G
MCMV_M35/1-519 TAWGP-NDKHRAESGLRRIYRALN-LEDNPFDLVEAVGDLDLSQGAYEGYVRHIYSLMKSLGHDVG
RCMV_pR35/1-521 RMFGP-PDIQRLDIGLRRIYRAAR-SDDNPYDLMEAVEDLDLPKGAYEKHLRTLYDLLQRVNVDLA
CaHV-2_GP35/1-552 IFMNS-PHRERLETGYRRLCQALN-DDSNPADMIQSLDDTYLPTSIYKHRLQHLYDLADSINIDVE
BHV_B35/1-571 GALSG-VNGERLDTAVRRLLSATR-STADPSELIENLAEVEILDGEYKKHLSAMYEIATGLNIKPT
TuHV-1_T35/1-567 RQLGE-TNLDRLDAGYRRLHRALE-GSDNPYHLVEDLTNTHLPLSRCQTHLHTLYRMCRQSRIETP
Consensus
+LF+GEP+K+RLDRGLRRLCRAAR+++DNP+D+I+SL+E+DLPDGVYK+AL++LYRL+DSLNIRP+
Occupancy

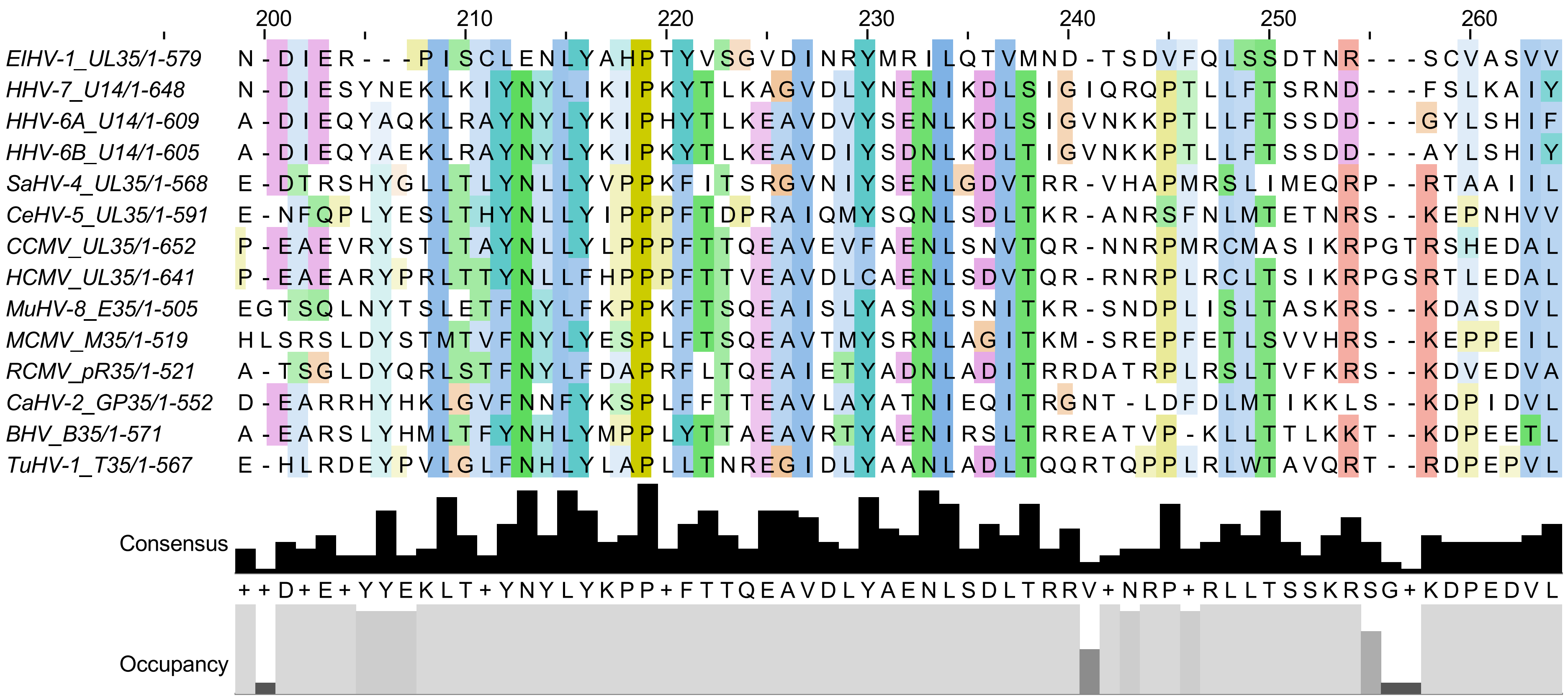

200
210
220
230
240
250
260
EIHV-1_UL35/1-579 N-DIER---PISCLENLYAHPTYVSGVDINRYMRILQTVMND-TSDVFQLSSDTNR---SCVASVV
HHV-7_U14/1-648 N-DIESYNEKLKIYNYLIKIPKYTLKAGVDLYNENIKDLSIGIQRQPTLLFTSRND---FSLKAIY
HHV-6A_U14/1-609 A-DIEQYAQKLRAYNYLYKIPHYTLKEAVDVYSENLKDLSIGVNKKPTLLFTSSDD---GYLSHIF
HHV-6B_U14/1-605 A-DIEQYAEKLRAYNYLYKIPKYTLKEAVDIYSDNLKDLTIGVNKKPTLLFTSSDD---AYLSHIY
SaHV-4_UL35/1-568 E-DTRSHYGLLTLYNLLYVPPKFITSRGVNIYSENLGDVTRR-VHAPMRSLIMEQRP--RTAAIIL
CeHV-5_UL35/1-591 E-NFQPLYESLTHYNLLYIPPPFTDPRAIQMYSQNLSDLTKR-ANRSFNLMTETNRS--KEPNHVV
CCMV_UL35/1-652 P-EAEVRYSTLTAYNLLYLPPPFTTQEAVEVFAENLSNVTQR-NNRPMRCMASIKRPGTRSHEDAL
HCMV_UL35/1-641 P-EAEARYPRLTTYNLLFHPPPFTTVEAVDLCAENLSDVTQR-RNRPLRCLTSIKRPGSRTLEDAL
MuHV-8_E35/1-505 EGTSQLNYTSLETFNYLFKPPKFTSQEAISLYASNLSNITKR-SNDPLISLTASKRS--KDASDVL
MCMV_M35/1-519 HLSRSLDYSTMTVFNYLYESPLFTSQEAVTMYSRNLAGITKM-SREPFETLSVVHRS--KEPPEIL
RCMV_pR35/1-521 A-TSGLDYQRLSTFNYLFDAPRFLTQEAIETYADNLADITRRDATRPLRSLTVFKRS--KDVEDVA
CaHV-2_GP35/1-552 D-EARRHYHKLGVFNNFYKSPLFFTTEAVLAYATNIEQITRGNT-LDFDLMTIKKLS--KDPIDVL
BHV_B35/1-571 A-EARSLYHMLTFYNHLYMPPLYTTAEAVRTYAENIRSLTRREATVP-KLLTTLKKT--KDPEETL
TuHV-1_T35/1-567 E-HLRDEYPVLGLFNHLYLAPLLTNREGIDLYAANLADLTQQRTQPPLRLWTAVQRT--RDPEPVL
Consensus
++D+E+YYEKLT+YNYLYKPP+FTTQEAVDLYAENLSDLTRRV+NRP+RLLTSSKRSG+KDPEDVL
Occupancy

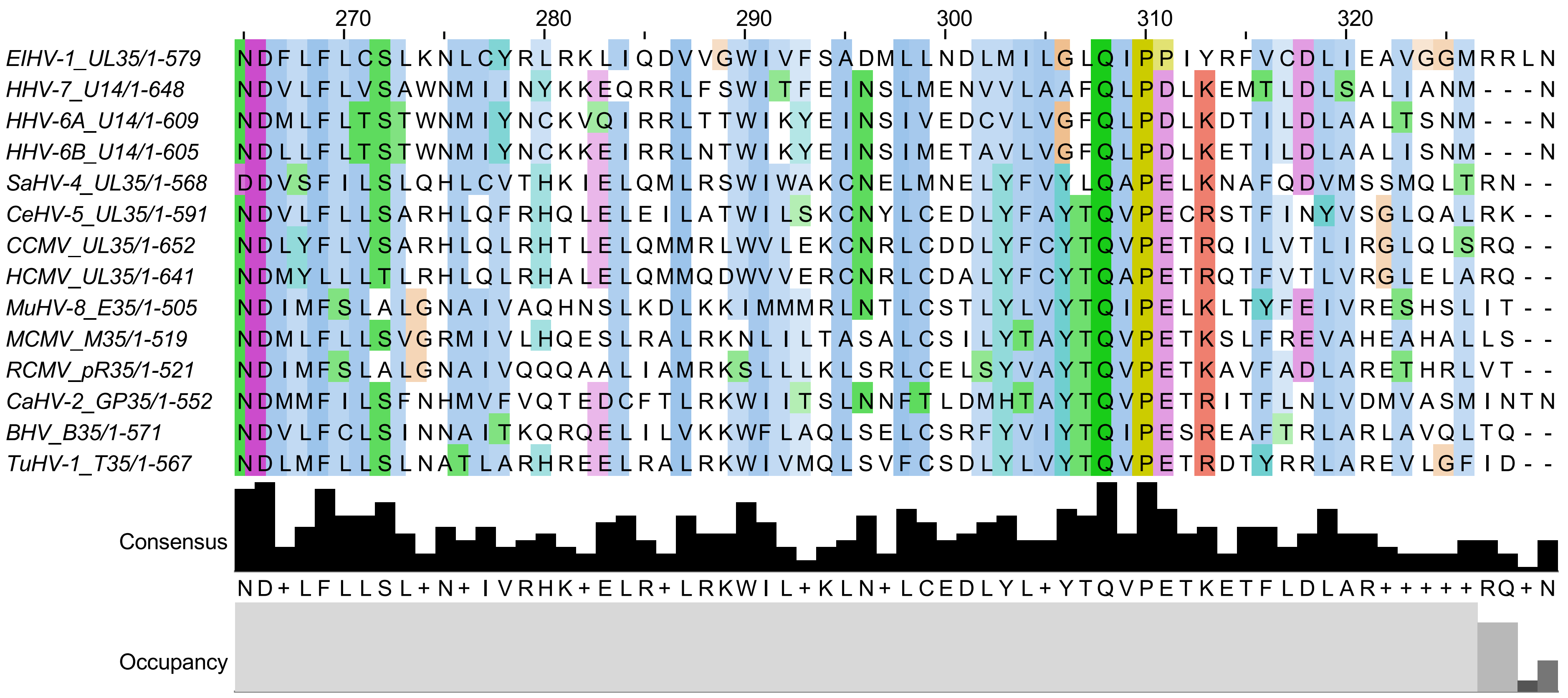

ElHV-1_UL35/1-579
HHV-7_U14/1-648
HHV-6A_U14/1-609
HHV-6B_U14/1-605
SaHV-4_UL35/1-568
CeHV-5_UL35/1-591
CCMV_UL35/1-652
HCMV_UL35/1-641
MuHV-8_E35/1-505
MCMV_M35/1-519
RCMV_pR35/1-521
CaHV-2_GP35/1-552
BHV_B35/1-571
TuHV-1_T35/1-567
Consensus
Occupancy
270
280
290
300
310
320

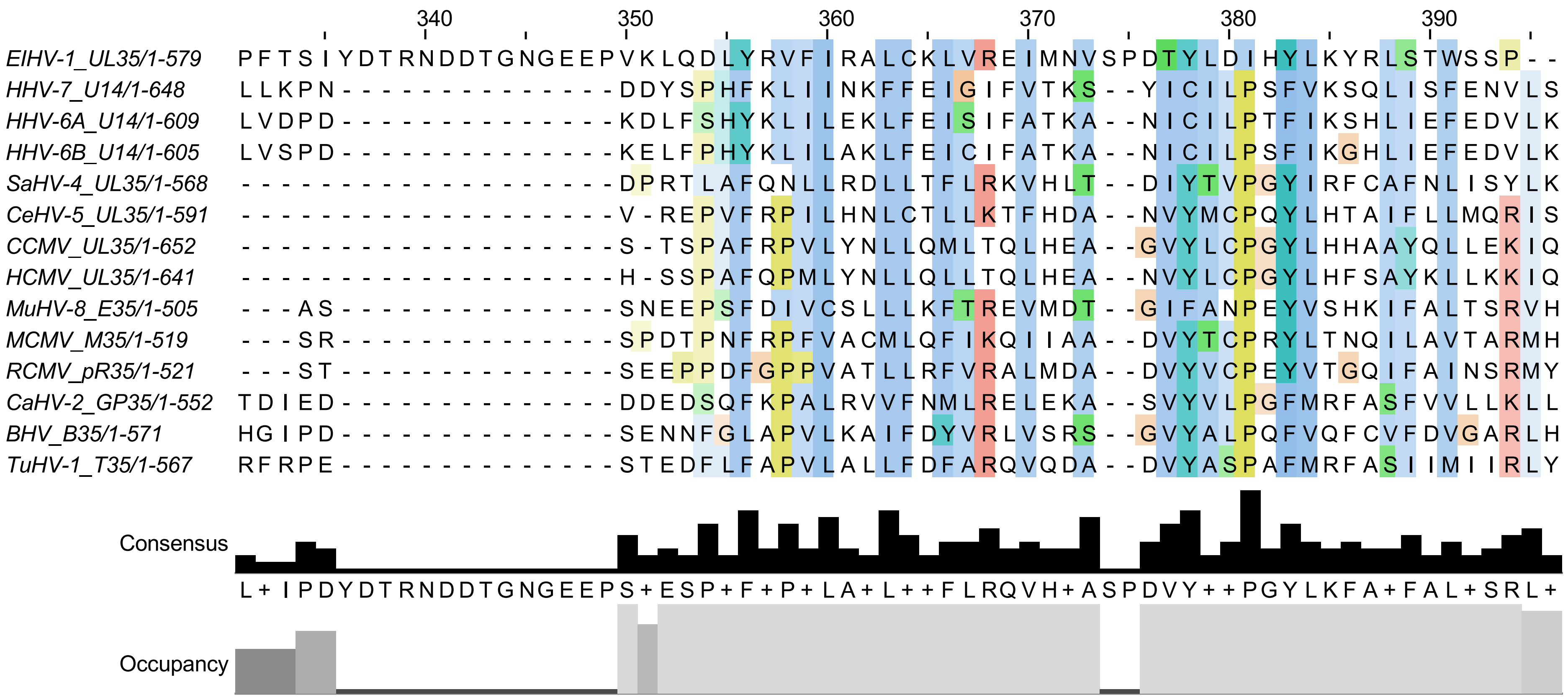
340 350 360 370 380 390
EIHV-1_UL35/1-579 PFTSIYDTRNDDTGNGEEPVKLQDLYRVFIRALCKLVREIMNVSPDTYLDIHYLKYRLSTWSSP--
HHV-7_U14/1-648 LLKPN---------------DDYSPHFKLIINKFFEIGIFVTKS--YICILPSFVKSQLISFENVLS
HHV-6A_U14/1-609 LVDPD---------------KDLFSHYKLILEKLFEISIFATKA--NICILPTFIKSHLIEFEDVLK
HHV-6B_U14/1-605 LVSPD---------------KELFPHYKLILAKLFEICIFATKA--NICILPSFIKGHLIEFEDVLK
SaHV-4_UL35/1-568 --------------------DPRTLAFQNLLRDLLTFLRKVHLT--DIYTVPGYIRFCAFNLISYLK
CeHV-5_UL35/1-591 --------------------V-REPVFRPILHNLCTLLKTFHDA--NVYMCPQYLHTAIFLLMQRIS
CCMV_UL35/1-652 --------------------S-TSPAFRPVLYNLLQMLTQLHEA--GVYLCPGYLHHAAYQLLEKIQ
HCMV_UL35/1-641 --------------------H-SSPAFQPMLYNLLQLLTQLHEA--NVYLCPGYLHFSAYKLLKKIQ
MuHV-8_E35/1-505 ---AS---------------SNEEPSFDIVCSLLLKFTREVMDT--GIFANPEYVSHKIFALTSRVH
MCMV_M35/1-519 ---SR---------------SPDTPNFRPFVACMLQFIKQIIAA--DVYTCPRYLTNQILAVTARMH
RCMV_pR35/1-521 ---ST---------------SEEPPDFGPPVATLLRFVRALMDA--DVYVCPEYVTGQIFAINSRMY
CaHV-2_GP35/1-552 TDIED---------------DDEDSQFKPALRVVFNMLRELEKA--SVYVLPGFMRFASFVVLLKLL
BHV_B35/1-571 HGIPD---------------SENNFGLAPVLKAIFDYVRLVSRS--GVYALPQFVQFCVFDVGARLH
TuHV-1_T35/1-567 RFRPE---------------STEDFLFAPVLALLFDFARQVQDA--DVYASPAFMRFASIIMIIRLY
Consensus
L+IPDYDTRNDDTGNGEEPS+ESP+F+P+LA+L++FLRQVH+ASPDVY++PGYLKFA+FAL+SRL+
Occupancy

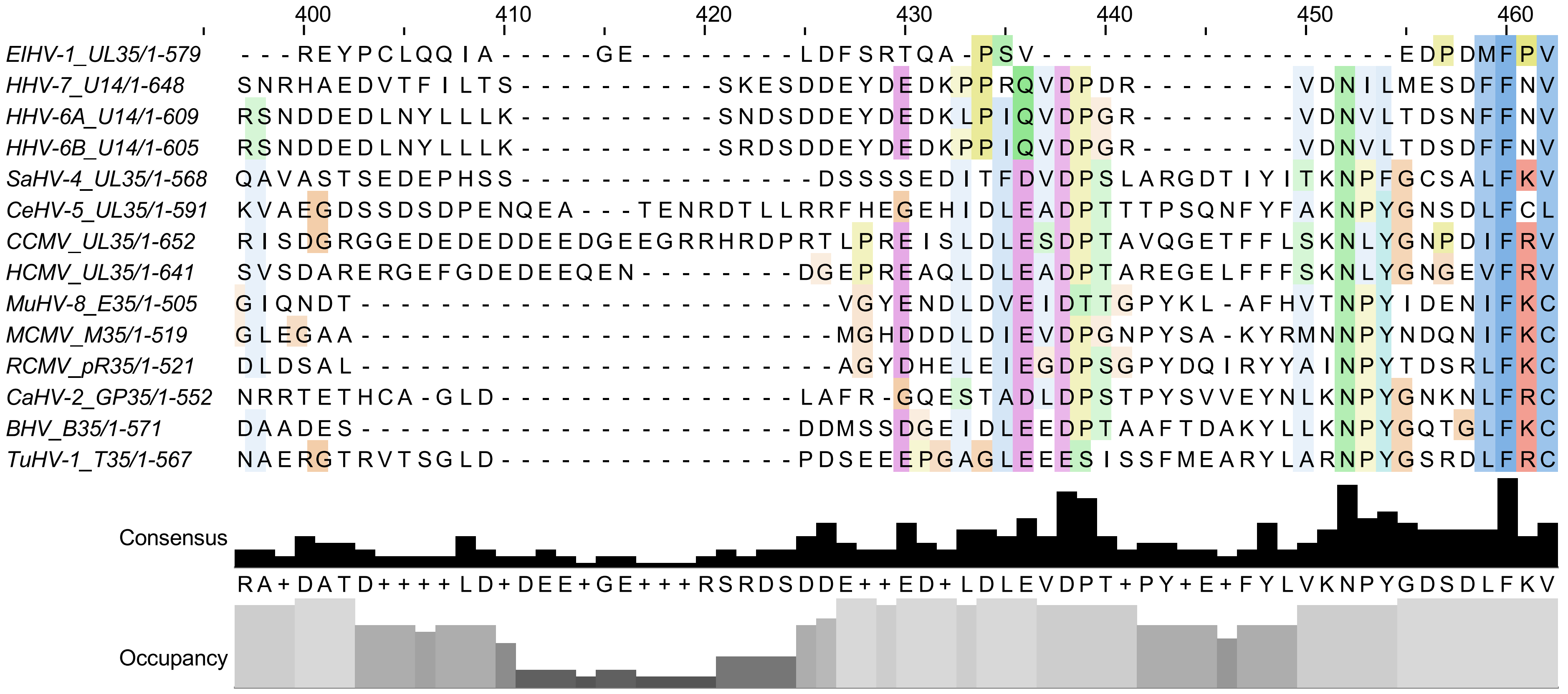
400
410
420
430
440
450
460
EIHV-1_UL35/1-579 ---REYPCLQQIA-----GE--------LDFSRTQA-PSV------------------EDPDMFPV
HHV-7_U14/1-648 SNRHAEDVTFILTS-----------SKESDDEYDEDKPPRQVDPDR--------VDNILMESDFFNV
HHV-6A_U14/1-609 RSNDDEDLNYLLLK-----------SNDSDDEYDEDKLPIQVDPGR--------VDNVLTDSNFFNV
HHV-6B_U14/1-605 RSNDDEDLNYLLLK-----------SRDSDDEYDEDKPPIQVDPGR--------VDNVLTDSDFFNV
SaHV-4_UL35/1-568 QAVASTSEDEPHSS----------------DSSSSEDITFDVDPSLARGDTIYITKNPFGCSALFKV
CeHV-5_UL35/1-591 KVAEGDSSDSDPENQEA---TENRDTLLRRFHEGEHIDLEADPTTTPSQNFYFAKNPYGNSDLFCL
CCMV_UL35/1-652 RISDGRGGEDEDEDDEEDGEEGRRHRDPRTLPREISLDLESDPTAVQGETFFLSKNLYGNPDIFRV
HCMV_UL35/1-641 SVSDARERGEFGDEDEEQEN--------DGEPREAQLDLEADPTAREGELFFFSKNLYGNGEVFRV
MuHV-8_E35/1-505 GIQNDT------------------------VGYENDLDVEIDTTGPYKL-AFHVTNPYIDENIFKC
MCMV_M35/1-519 GLEGAA------------------------MGHDDDLDIEVDPGNPYSA-KYRMNNPYNDQNIFKC
RCMV_pR35/1-521 DLDSAL------------------------AGYDHELEIEGDPSGPYDQIRYYAINPYTDSRLFKC
CaHV-2_GP35/1-552 NRRTETHCA-GLD----------------LAFR-GQESTADLDPSTPYSVVEYNLKNPYGNKNLFRC
BHV_B35/1-571 DAADES------------------------DDMSSDGEIDLEEDPTAAFTDAKYLLKNPYGQTGLFKC
TuHV-1_T35/1-567 NAERGTRVTSGLD----------------PDSEEEPGAGLEEEESISSFMEARYLARNPYGSRDLFRC
Consensus
RA+DATD++++LD+DEE+GE+++RSRDSDDE++ED+LDLEVDPT+PY+E+FYLVKNPYGDSDLFKV
Occupancy

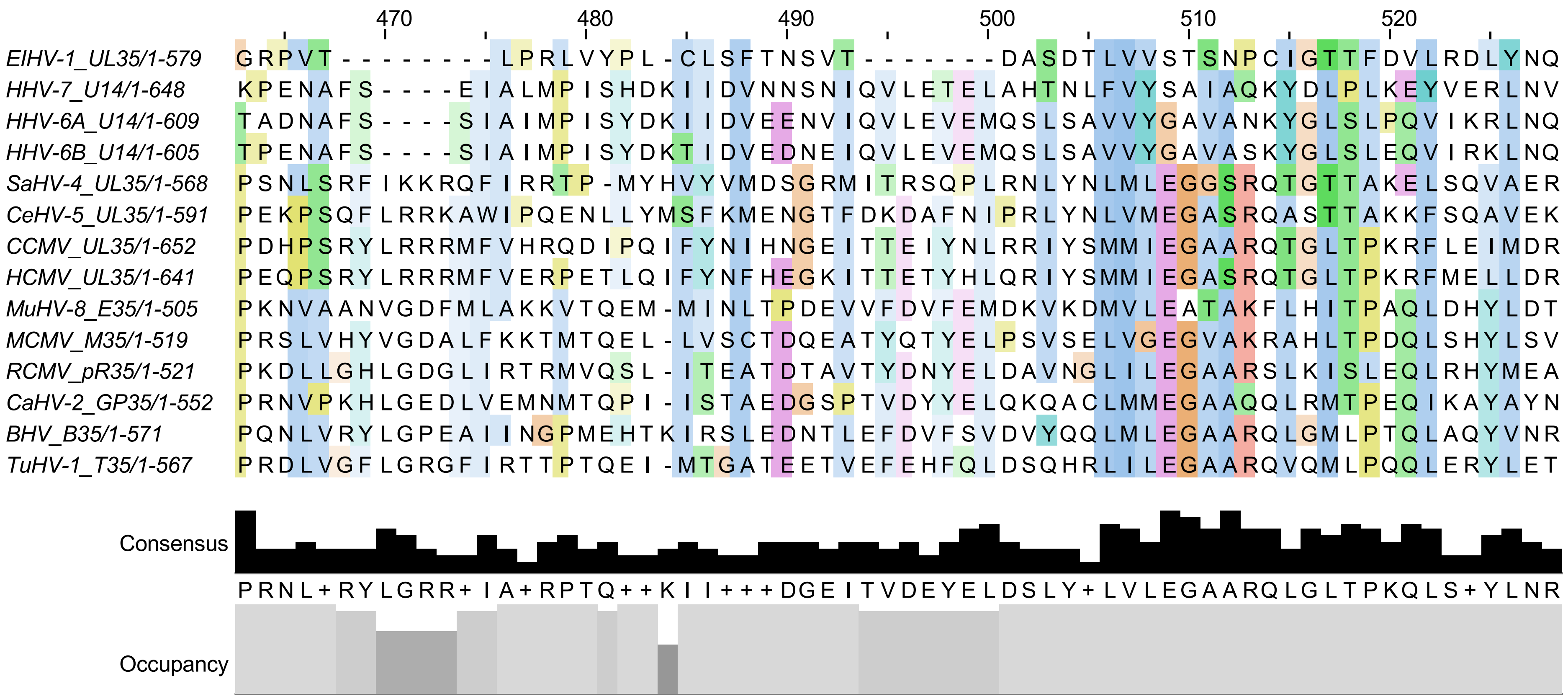
470
480
490
500
510
520
EIHV-1_UL35/1-579 GRPVT--------LPRLVYPL-CLSFTNSVT-------DASDTLVVSTSNPCIGTTFDVLRDLYNQ
HHV-7_U14/1-648 KPENAFS----EIALMPISHDKIIDVNNSNIQVLETELAHTNLFVYSAIAQKYDLPLKEYVERLNV
HHV-6A_U14/1-609 TADNAFS----SIAIMPISYDKIIDVEENVIQVLEVEMQSLSAVVYGAVANKYGLSLPQVIKRLNQ
HHV-6B_U14/1-605 TPENAFS----SIAIMPISYDKTIDVEDNEIQVLEVEMQSLSAVVYGAVASKYGLSLEQVIRKLNQ
SaHV-4_UL35/1-568 PSNLSRFIKKRQFIRRTP-MYHVYVMDSGRMITRSQPLRNLYNLMLEGGSRQTGTTAKELSQVAER
CeHV-5_UL35/1-591 PEKPSQFLRRKAWIPQENLLYMSFKMENGTFDKDAFNIPRLYNLVMEGASRQASTTAKKFSQAVEK
CCMV_UL35/1-652 PDHPSRYLRRRMFVHRQDIPQIFYNIHNGEITTEIYNLRRIYSMMIEGAARQTGLTPKRFLEIMDR
HCMV_UL35/1-641 PEQPSRYLRRRMFVERPETLQIFYNFHEGKITTETYHLQRIYSMMIEGASRQTGLTPKRFMELLDR
MuHV-8_E35/1-505 PKNVAANVGDFMLAKKVTQEM-MINLTPDEVVFDVFEMDKVKDMVLEATAKFLHITPAQLDHYLDT
MCMV_M35/1-519 PRSLVHYVGDALFKKTMTQEL-LVSCTDQEATYQTYELPSVSELVGEGVAKRAHLTPDQLSHYLSV
RCMV_pR35/1-521 PKDLLGHLGDGLIRTRMVQSL-ITEATDTAVTYDNYELDAVNGLILEGAARSLKISLEQLRHYMEA
CaHV-2_GP35/1-552 PRNVPKHLGEDLVEMNMTQPI-ISTAEDGSPTVDYYELQKQACLMMEGAAQQLRMTPEQIKAYAYN
BHV_B35/1-571 PQNLVRYLGPEAIINGPMEHTKIRSLEDNTLEFDVFSVDVYQQLMLEGAARQLGMLPTQLAQYVNR
TuHV-1_T35/1-567 PRDLVGFLGRGFIRTTPTQEI-MTGATEETVEFEHFQLDSQHRLILEGAARQVQMLPQQLERYLET
Consensus
PRNL+RYLGRR+IA+RPTQ++KII+++DGEITVDEYELDSLY+LVLEGAARQLGLTPKQLS+YLNR
Occupancy

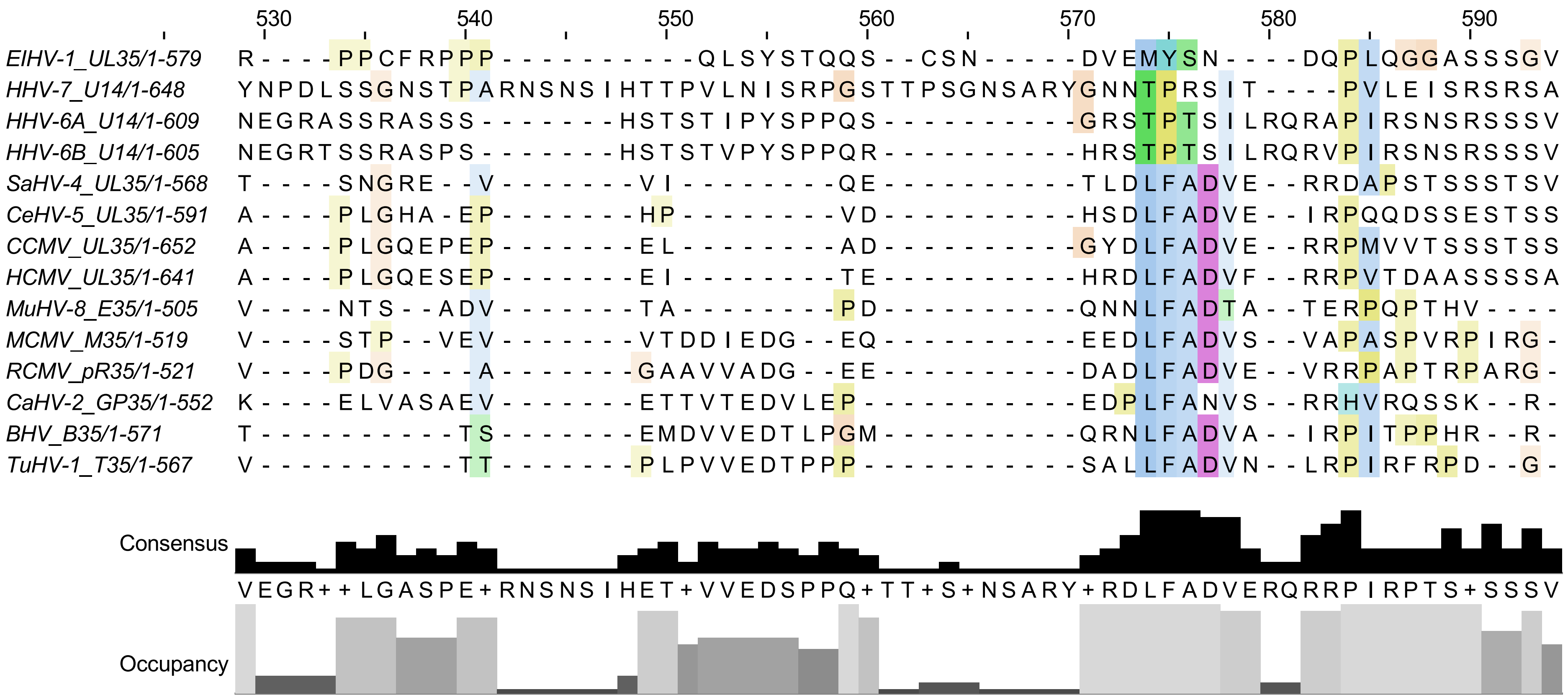

530
540
550
560
570
580
590
EIHV-1_UL35/1-579 R - - - - P P C F R P P P - - - - - - - - - - - Q L S Y S T Q Q S - - C S N - - - - - D V E M Y S N - - - - D Q P L Q G G A S S S G V
HHV-7_U14/1-648 Y N P D L S S G N S T P A R N S N S I H T T P V L N I S R P G S T T P S G N S A R Y G N N T P R S I T - - - - P V L E I S R S R S A
HHV-6A_U14/1-609 N E G R A S S R A S S S - - - - - - - H S T S T I P Y S P P Q S - - - - - - - - - - G R S T P T S I L R Q R A P I R S N S R S S S V
HHV-6B_U14/1-605 N E G R T S S R A S P S - - - - - - - H S T S T V P Y S P P Q R - - - - - - - - - - H R S T P T S I L R Q R V P I R S N S R S S S V
SaHV-4_UL35/1-568 T - - - - S N G R E - - V - - - - - - - V I - - - - - - - - Q E - - - - - - - - - - T L D L F A D V E - - R R D A P S T S S S T S V
CeHV-5_UL35/1-591 A - - - - P L G H A - E P - - - - - - - H P - - - - - - - - V D - - - - - - - - - - H S D L F A D V E - - I R P Q Q D S S E S T S S
CCMV_UL35/1-652 A - - - - P L G Q E P E P - - - - - - - E L - - - - - - - - A D - - - - - - - - - - G Y D L F A D V E - - R R P M V V T S S S T S S
HCMV_UL35/1-641 A - - - - P L G Q E S E P - - - - - - - E I - - - - - - - - T E - - - - - - - - - - H R D L F A D V F - - R R P V T D A A S S S S A
MuHV-8_E35/1-505 V - - - - N T S - - A D V - - - - - - - T A - - - - - - - - P D - - - - - - - - - - Q N N L F A D T A - - T E R P Q P T H V - - - -
MCMV_M35/1-519 V - - - - S T P - - V E V - - - - - - - V T D D I E D G - - E Q - - - - - - - - - - E E D L F A D V S - - V A P A S P V R P I R G -
RCMV_pR35/1-521 V - - - - P D G - - - - A - - - - - - - G A A V V A D G - - E E - - - - - - - - - - D A D L F A D V E - - V R R P A P T R P A R G -
CaHV-2_GP35/1-552 K - - - - E L V A S A E V - - - - - - - E T T V T E D V L E P - - - - - - - - - - - E D P L F A N V S - - R R H V R Q S S K - - R -
BHV_B35/1-571 T - - - - - - - - - - T S - - - - - - - E M D V V E D T L P G M - - - - - - - - - - Q R N L F A D V A - - I R P I T P P H R - - R -
TuHV-1_T35/1-567 V - - - - - - - - - - T T - - - - - - - P L P V V E D T P P P - - - - - - - - - - - S A L L F A D V N - - L R P I R F R P D - - G -
Consensus
V E G R + + L G A S P E + R N S N S I H E T + V V E D S P P Q + T T + S + N S A R Y + R D L F A D V E R Q R R P I R P T S + S S S V
Occupancy

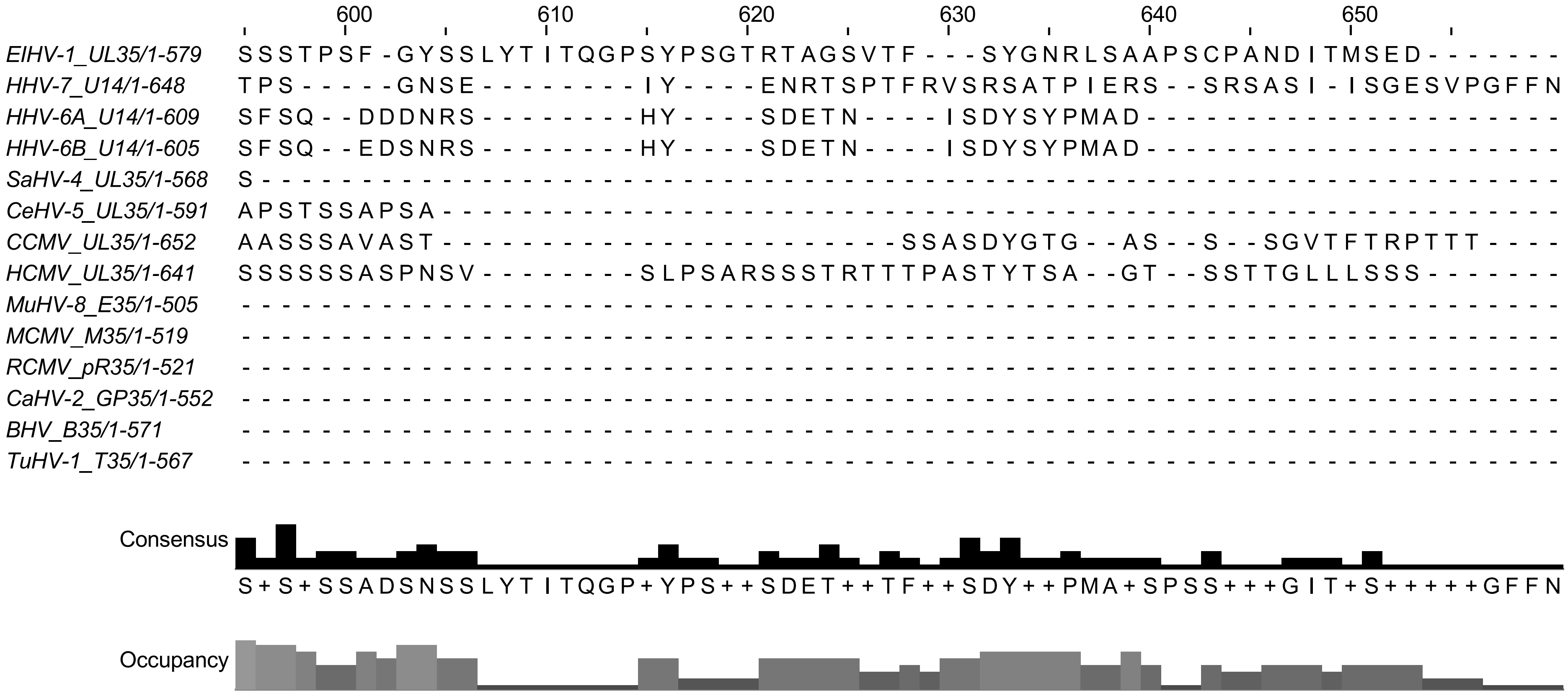

600 610 620 630 640 650
ElHV-1_UL35/1-579 SSSTPSF-GYSSLYTITQGPSYPSGTRTAGSVTF---SYGNRLSAAPSCPANDITMSED-------
HHV-7_U14/1-648 TPS-----GNSE---------IY----ENRTSPTFRVSRSATPIERS--SRSASI-ISGESVPGFFN
HHV-6A_U14/1-609 SFSQ--DDDNRS--------HY----SDETN----ISDYSYPMAD---------------------
HHV-6B_U14/1-605 SFSQ--EDSNRS--------HY----SDETN----ISDYSYPMAD---------------------
SaHV-4_UL35/1-568 S------------------------------------------------------------------
CeHV-5_UL35/1-591 APSTSSAPSA---------------------------------------------------------
CCMV_UL35/1-652 AASSSAVAST--------------------SSASDYGTG--AS--S--SGVTFTRPTTT----
HCMV_UL35/1-641 SSSSSSASPNSV--------SLPSARSSSTRTTTPASTYTSA--GT--SSTTGLLLSSS-------
MuHV-8_E35/1-505
MCMV_M35/1-519
RCMV_pR35/1-521
CaHV-2_GP35/1-552
BHV_B35/1-571
TuHV-1_T35/1-567
Consensus
S+S+SSADSNSSLYTITQGP+YPS++SDET++TF++SDY++PMA+SPSS+++GIT+S+++++GFFN
Occupancy

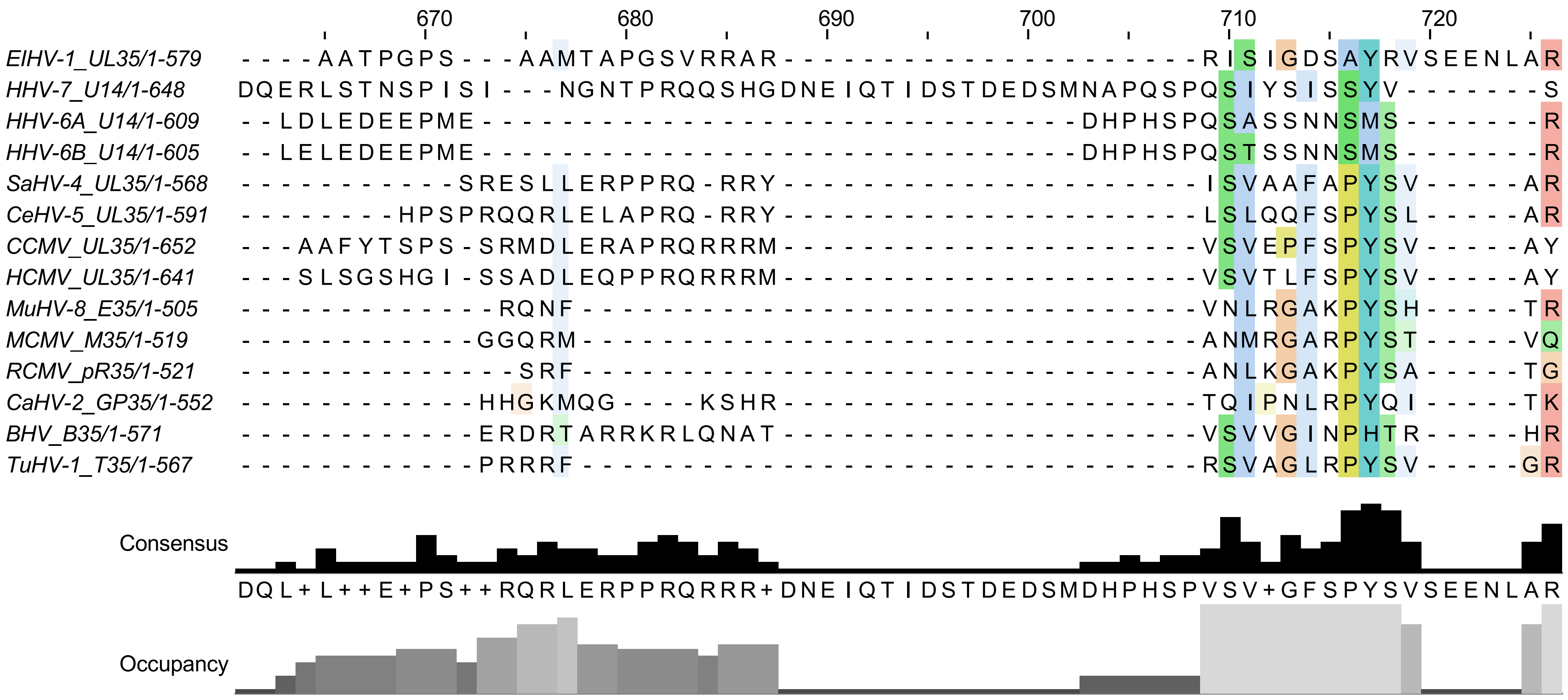
670
680
690
700
710
720
EIHV-1_UL35/1-579
HHV-7_U14/1-648
HHV-6A_U14/1-609
HHV-6B_U14/1-605
SaHV-4_UL35/1-568
CeHV-5_UL35/1-591
CCMV_UL35/1-652
HCMV_UL35/1-641
MuHV-8_E35/1-505
MCMV_M35/1-519
RCMV_pR35/1-521
CaHV-2_GP35/1-552
BHV_B35/1-571
TuHV-1_T35/1-567
Consensus
Occupancy

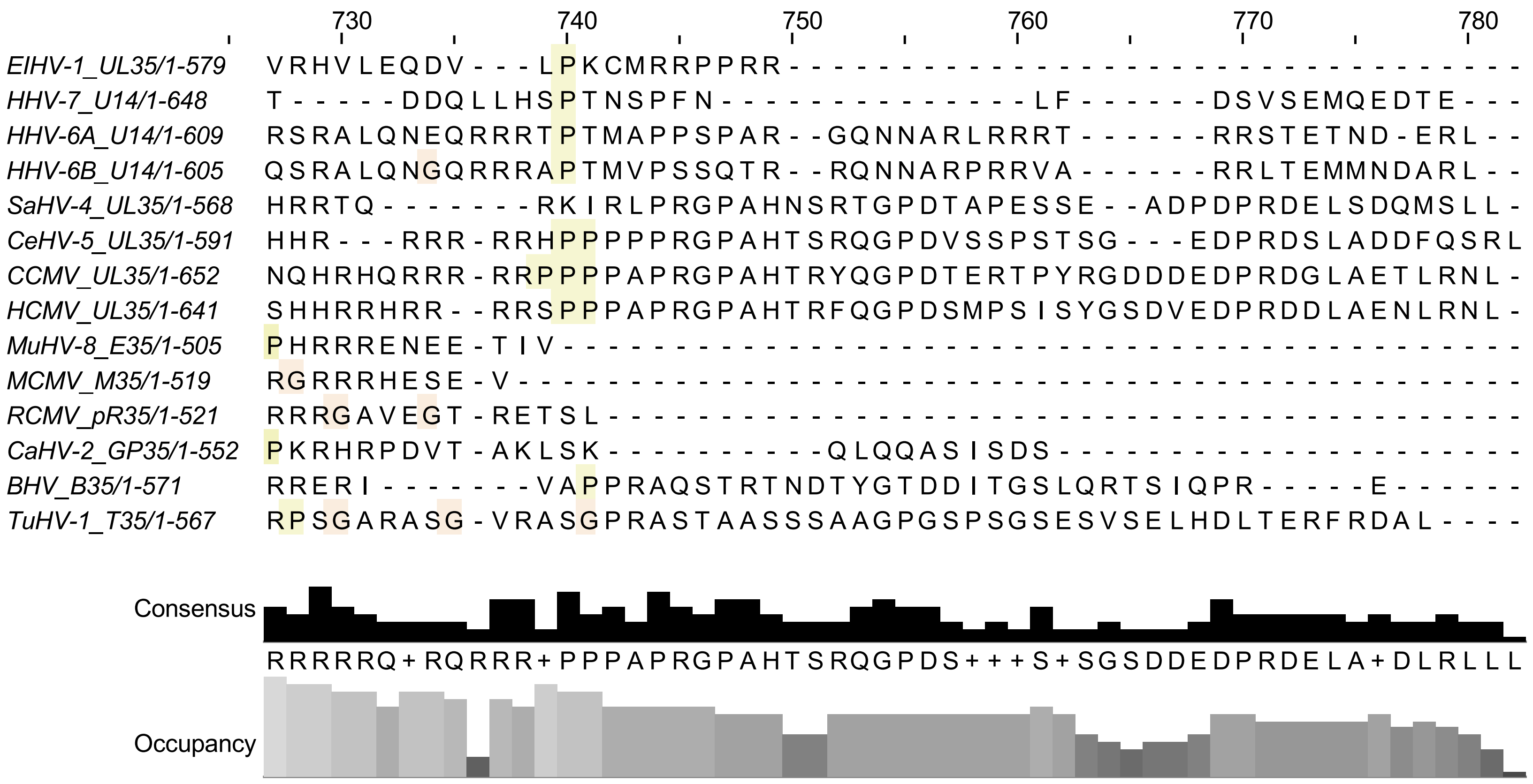

730
740
750
760
770
780
EIHV-1_UL35/1-579 VRHVLEQDV---LPKCMRRPPRR---------------------------------
HHV-7_U14/1-648 T-----DDQLLHSPTNSPFN--------------LF------DSVSEMQEDTE---
HHV-6A_U14/1-609 RSRALQNEQRRRTPTMAPPSPAR--GQNNARLRRRT------RRSTETND-ERL--
HHV-6B_U14/1-605 QSRALQNGQRRRAPTMVPSSQTR--RQNNARPRRVA------RRLTEMMNDARL--
SaHV-4_UL35/1-568 HRRTQ-------RKIRLPRGPAHNSRTGPDTAPESSE--ADPDPRDELSDQMSLL-
CeHV-5_UL35/1-591 HHR---RRR-RRHPPPPPRGPAHTSRQGPDVSSPSTSG---EDPRDSLADDFQSRL
CCMV_UL35/1-652 NQHRHQRRR-RRPPPPAPRGPAHTRYQGPDTERTPYRGDDDEDPRDGLAETLRNL-
HCMV_UL35/1-641 SHHRRHRR--RRSPPPAPRGPAHTRFQGPDSMPSISYGSDVEDPRDDLAENLRNL-
MuHV-8_E35/1-505 PHRRRENEE-TIV-------------------------------------------
MCMV_M35/1-519 RGRRRHESE-V---------------------------------------------
RCMV_pR35/1-521 RRRGAVEGT-RETSL-----------------------------------------
CaHV-2_GP35/1-552 PKRHRPDVT-AKLSK----------QLQQASISDS---------------------
BHV_B35/1-571 RRERI-------VAPPRAQSTRTNDTYGTDDITGSLQRTSIQPR-----E------
TuHV-1_T35/1-567 RPSGARASG-VRASGPRASTAASSSAAGPGSPSGSESVSELHDLTERFRDAL----
Consensus
RRRRRQ+RQRRR+PPPAPRGPAHTSRQGPDS+++S+SGSDDEDPRDELA+DLRLLL
Occupancy
